# Changes in the intracranial volume from early adulthood to the sixth decade of life: A longitudinal study

**DOI:** 10.1101/677898

**Authors:** Yaron Caspi, Rachel M. Brouwer, Hugo G. Schnack, Marieke E. van de Nieuwenhuijzen, Wiepke Cahn, René S. Kahn, Wiro J. Niessen, Aad van der Lugt, Hilleke Hulshoff Pol

## Abstract

Normal brain-aging occurs at all structural levels. Excessive pathophysiological changes in the brain, beyond the normal one, are implicated in the etiology of brain disorders such as severe forms of the schizophrenia spectrum and dementia. To account for brain-aging in health and disease, it is critical to study the age-dependent trajectories of brain biomarkers at various levels and among different age groups.

The intracranial volume (ICV) is a key biological marker, and changes in the ICV during the lifespan can teach us about the biology of development, aging, and gene X environment interactions. However, whether ICV changes with age in adulthood is not resolved.

Applying a semi-automatic in-house-built algorithm for ICV extraction on T1w MR brain scans in the Dutch longitudinal cohort (GROUP), we measured ICV changes. Individuals between the ages of 16 and 55 years were scanned up to three consecutive times with 3.32±0.32 years between consecutive scans (N=482, 359, 302). Using the extracted ICVs, we calculated ICV longitudinal aging-trajectories based on three analysis methods; direct calculation of ICV differences between the first and the last scan, fitting all ICV measurements of individuals to a straight line and applying a global linear mixed model fitting. We report statistically significant increases in the ICV in adulthood until the fourth decade of life (average change +0.03%/y, or about 0.5 ml/y, at age 20), and decreases in the ICV afterward (−0.09%/y, or about −1.2 ml/y, at age 55). To account for previous cross-sectional reports of ICV changes, we analyzed the same data using a cross-sectional approach. Our cross-sectional analysis detected ICV changes consistent with the previously reported cross-sectional effect. However, the reported amount of cross-sectional changes within this age range was significantly larger than the longitudinal changes. We attribute the cross-sectional results to a generational effect.

In conclusion, the human intracranial volume does not stay constant during adulthood but instead shows a small increase during young adulthood and a decrease thereafter from the fourth decade of life. The age-related changes in the longitudinal setup are smaller than those reported using cross-sectional approaches and unlikely to affect structural brain imaging studies correcting for intracranial volume considerably. As to the possible mechanisms involved, this awaits further study, although thickening of the meninges and skull bones have been proposed, as well as a smaller amount of brain fluids addition above the overall loss of brain tissue.

## Introduction

Aging is a natural process that occurs in most species of the animal kingdom (Jones et al., 2014). In *Homo Sapiens*, aging starts in adulthood and continues into old age (Lindle et al., 1997). Similar to the body, age-dependent structural changes in the brain, from the cellular level to the system one, result in brain aging (Lockhart and DeCarli, 2014; Kennedy and Raz, 2015). These structural brain-aging processes are, on average, accompanied by cognitive performances differences across the lifespan (Hartshorne and Germine, 2015).

The rate of brain-aging across different brain systems is, however, not constant. Instead, various brain regions show different aging trajectories (Hedman et al., 2011). For example, the cingulate atrophy rate is larger than that of the cerebellum cortex lobes (Pfefferbaum et al., 2013). In principle, regional structural aging-changes can reside within some accepted boundaries, in which case they characterize the normal aging process. By contrast, structural brain aging-changes can also occur beyond these normal boundaries, in which case they are the manifestation of disease-like degenerative processes. Considering the high impact of degenerative brain-processes on global population health, it is crucial to infer normal structural aging-trajectories of a wide range of brain systems and biomarkers and differentiate them from those in disease conditions.

One such biomarker is the total intracranial volume (ICV). Physiologically, the cranial bone is separated from the brain system by three membranal layers (meninges); the dura, the arachnoid, and the pia maters (Adeeb et al., 2012). Beneath these layers, within the ICV, all the central-nervous-system neuronal and glial cells, as well as the cerebrospinal fluid (CSF) are located (Jenkins et al., 2000). Since it is hard to delineate the meninges from the cranial bone in magnetic resonance imaging (MRI), in practice the ICV is mostly measured by identifying the sum-total volumes of the gray matter, white matter, and the CSF.

The ICV, demarcating the maximum size of the brain, has a long history for being utilized as a biomarker. Before the invention of advanced imaging techniques, such as MRI and computed tomography (CT), the size of the cranium served as an approximate measure for the ICV and the brain. At the beginning of the 20th century, it becomes the center of the efforts to show human bodily environmental plasticity (Boas, 1912; Sparks and Jantz, 2002; Gravlee et al., 2003). Later on, it was shown that extreme environmental conditions during the prenatal period influence the ICV at adulthood (Hulshoff Pol et al., 2000). It is also known that the ICV size is under the control of a large genetic component (Peper et al., 2007; Stein et al., 2012; Batouli et al., 2014; Adams et al., 2016). Recently, it was also found that the ICV correlates with several psychiatric conditions. For example, a small but significant correlation was found between schizophrenia disorder and a reduced ICV (Baaré et al., 2001; Haijma et al., 2012; van Erp et al., 2016; Smeland et al., 2017). Similarly, the ICV was the only brain volumetric biomarker that shows a genetic correlation with ADHD (Klein et al., 2019). The ICV is also strongly correlated with sex and head size. Thus, females, on average, have a smaller ICV (Ruigrok et al., 2014), and people with bigger heads have, on average, larger ICV (Wolf et al., 2003). In addition, probably due to its correlation with maximal brain size during the lifespan, the ICV is being used as a biomarker for the ‘brain reserve’ protective factor in dementia research (van Loenhoud et al., 2018). Finally, the ICV correlates with cognitive abilities (MacLullich et al., 2002) (although the effect size may have been overestimated (Pietschnig et al., 2015)). Similarly, genes for the whole brain volume and the ICV overlap with genes for general cognitive ability and with intelligence as is often measured by the intelligent quotient (IQ) (Posthuma et al., 2002; Deary et al., 2010).

Although the ICV serves as an elementary biomarker that correlates with a wide range of medical and physiological conditions, a comprehensive work that elucidates its exact aging-trajectory is still missing. Instead, many neuroimaging brain-aging studies use the ICV to regress out the influence of the head size on volumes or area of specific brain regions (Ikram et al., 2008; Dickie et al., 2013). In those cases, the ICV aging-trajectory is rarely reported for its own sake. We believe that the ICV aging-trajectory may have merits in itself. Thus, the work presented here aims at filling the gap in the understanding of the normal ICV aging during young and middle adulthood.

Several cross-sectional studies have reported ICV changes as a function of age. In general, the current scientific literature gravitates around two views. The first view suggests that detectable age-related ICV changes exist. The second view suggests that the ICV stays constant during adulthood. For the first view, a decline of the ICV of 1.45 ml/year for males and 1.82 ml/year for females was measured from the middle of the 4th to the 9th decade of life (0.1 %/year and 0.15 %/year for males and females respectively based on the ICV at age 44) in one study (DeCarli et al., 2005). In another study a reduction in the inner skull volume (which is roughly equal to the ICV) was also observed (Fillmore et al., 2015). The ICV volume decline started only around the fifth decade of life, and its reduction rate was not homogeneous. In total, in that study, from the age of ∼ 40 to the age of ∼ 75 years there was a decrease of about 10% in the inner skull volume (∼ 0.29%/year).

By contrast, other cross-sectional studies reported no changes in the ICV with age. For example, no changes in ICV were found in individuals from the age of 14 years to the 8th decade of life (Courchesne et al., 2000), and in young adulthood (20-25 years) compared to older adults (65-90 years)(Buckner et al., 2004). A similar negative finding was obtained when using a T2w MRI cohort of individuals between the ages of 15 and 59 years (Hasan et al., 2010), and in people between the ages of 18 and 95 using CT imaging (Ricard et al., 2010). Finally, in 208 individuals combined with the publicly available OASIS cohort between the ages of 35 to 75, no dependency of ICV on age was found (Schippling et al., 2017).

In some cases, when ICV age-related changes were observed, they were interpreted in the context of the secular trend of increasing head size in the last century. For example, Good et al. observed a quadratic decline in ICV with age for males, but not for females, and suggested that generational head growth is the cause of their findings (Good et al., 2001). Similarly, Kruggel has observed a small decrease in the ICV (0.84 ml/year between the ages of 16 and 70) that was significant at the 0.1 false-positive statistical level and interpreted the effect in the context of a generational growth (Kruggel, 2006). Finally, Kim et al. observed an average ICV decrease of 2.1 ml/year for males and females between a group of Korean people at an average age of 68 years and a second group at an average age of 24 years (Kim et al., 2018). They also suggested environmental factors as the cause of this ICV difference.

Since all these studies applied a cross-sectional analysis, it is hard to estimate what factor really represents a generational effect, and to what extent real individual age-dependent ICV changes exist. However, precisely this estimation is needed to infer the true aging-trajectory of the ICV. One way to obtain an accurate measure of the ICV aging-trajectory is by using a longitudinal approach. In a longitudinal approach, each person serves as its own normalization baseline, and thus the generational effect is removed. Indeed, it is known that applying cross-sectional and longitudinal analyses to the same data can give rise to substantial differences in the results (Pfefferbaum and Sullivan, 2015).

To the best of our knowledge, only two studies have used a longitudinal (or semi-longitudinal) approach to study ICV aging. In the first longitudinal study, 90 healthy subjects between the ages of 14 and 77 years had two MRI scans at an average interval of 3.5 years (Liu et al., 2003). To study longitudinal changes, the authors divided their cohort in three age epochs ≤ 34; 35 − 54; and *≥* 54 years). Their analysis showed a small increase in the ICV over repeated measurements for the youngest group (4.6 ml; 0.3 %). However, the ICV was not statistically significant different between the first and second measurements for the other two age epochs. In addition, the average ICV for the first age group was not different from that in the second and third age groups. By contrast, when a cross-sectional analysis was applied to the data, a statistically significant correlation with age was found, which was interpreted as the result of a secular growth of the head.

The second study compared two types of delineations of the ICV (Royle et al., 2013). First, an automatic ICV extraction in a cohort of 60 people between the ages of 71 years and 74 years was applied. Next, a manual expert delineation of the ICV plus the part of the inner skull table that was assessed by the human expert to results from age-related thickening was conducted. By comparing these two measurements, a ‘longitudinal’ ICV changes from the start of aging to the eighth decade of life was inferred. The results suggested an average of 101 ml (6.2 %) for males and 114 ml (8.3 %) for females individual ICV reduction during the course of life.

Assuming that the main effect that was detected in the cross-sectional ICV analyses was due to a generational growth, these two longitudinal studies raise a series of questions. Does the ICV indeed continue to enlarge at young adulthood? How come Liu et al. did not see ICV reduction at middle and advanced adulthood, while Royle et al. saw large longitudinal ICV decrease? How come the effect that Royle et al. measured is as large as the comparable cross-sectional effect that is generational by origin? Do these findings suggest a non-linear aging-trajectory for the ICV? In light of these questions, it is clear that additional work is needed regarding ICV and aging.

To assess ICV aging in young and middle adulthood, we measured ICV trajectories in a large longitudinal Dutch cohort with three repeated measurements. Using an in-house-built algorithm for ICV extraction, we assessed ICV changes over the repeated measurements. We analyzed the results using three longitudinal measures: the individual volume difference between the first and last measurement, individual fits to all available ICV measurements, and a global linear mixed model fitting. These analysis methods showed a small but statistically significant enlargement of the ICV at young adulthood that is replaced by a small but statistically significant ICV shrinkage at middle adulthood.

Despite its advantages, a longitudinal design can also introduce new confounding factors. In particular, in many cases, as a result of the lengthy period of the longitudinal study, changes occur in the scanner parameters (upgrade of software or hardware). Alternatively, it may become necessary to use different scanners for different waves of the study. Indeed, it is known that changes of scanner or scanner parameters can introduce confounding factors to segmentation analysis of brain images (Han et al., 2006; Jovicich et al., 2009). In general, these confounding factors are more significant when different scanners are used for the measurements than when the same scanner is used with different software or hardware parameters. Although it is essential to consider these confounding factors in cross-sectional studies, it is much more critical to consider them in longitudinal-designed studies. The reason for this importance stems directly from the fact that in longitudinal studies, each person serves as its own normalization baseline. Thus, the underlying assumption of longitudinal imaging studies is that any longitudinal changes in the image are due to changes in the measured property of the subject. This assumption is, of course, being violated by the confounding factors that are related to the scanner itself.

It should be noted that the magnitude of such confounding factors are reduced when the skull is taken into consideration in the segmentation algorithm (Takao et al., 2011). In our work, we have used a segmentation algorithm that takes the skull into account, and hence, scanner related confounding factor should be minimal. Yet, to increase precision, we have chosen to address the issue of scanner-introduced confounding factors overtly in the context of a linear mixed model analysis by directly using the scanner identification.

## Methods

### Data

We used the T1-weighted (T1w) MRI scans from the Utrecht site (The Netherlands) of the GROUP study (Korver et al., 2012). The GROUP study was originally designed to assess the risks and outcome of psychosis. Hence, it contains three groups of participants — namely — a control group, people that were diagnosed with schizophrenia and siblings of people that were diagnosed with schizophrenia. In general, in our analysis, we do not differentiate between these three groups of people. Altogether, the Utrecht GROUP Study contains T1w scans of 528, 378 and 309 individuals at the first, second and third time points respectively (see Table 1). The average period between consecutive scans is 3.32±0.32 years. The age span at the first time point is between 16 and 55 years. All scans were recorded using a 1.5-tesla Philips scanner at the University Medical Center of Utrecht. Note that during the acquisition phase of the GROUP study, two different scanners with the same field strength and the same acquisition protocol were used to collect the data between different waves of the study, as well as within each wave. Of some cases, scanner ID was missing. Altogether, we lack such knowledge for 22, 6, and 29 cases at the first, second, and third waves of the study. Out of these cases, we extracted ICV data for 14, 6, and 24 cases (see extraction method below.) For the exact acquisition protocol of the T1w scans see (Boos et al., 2011).

**Table 1:** Statistical Characteristics of the GROUP study. Participants were scanned on a 1.5 Tesla MRI machine (Philips) from one to three times. Top part - statistical characteristics of each wave of the study and the extracted ICV values. Bottom part - statistical characteristic of the longitudinal statistics of the study and the extracted ICV values. T_*x*_ - x wave of the study. Parenthesis - the number of females.

| Scan | # of subjects | # of subjects with ICV extracted | # of Schizophrenia diagnosed subjects with ICV extracted | # of siblings with ICV extracted | # of control subjects with ICV extracted | Average (±s.d.)age (years) |
| --- | --- | --- | --- | --- | --- | --- |
| **Cross-sectional** |  |  |  |  |  |  |
| T1 | 528 (224) | 482 (213) | 152 (34) | 214 (115) | 116 (64) | 27±7 |
| T2 | 378 (162) | 359 (156) | 110 (20) | 168 (90) | 81 (46) | 30±7 |
| T3 | 309 (131) | 302 (130) | 86 (16) | 135 (72) | 81 (42) | 34±8 |
| **Longitudinal** |  |  |  |  |  |  |
| T1-T2 | 339 (143) | 292 (130) | 82 (14) | 145 (77) | 65 (39) | — |
| T2-T3 | 254 (107) | 243 (104) | 73 (14) | 108 (55) | 62 (35) |  |
| T1-T3 | 260 (109) | 237 (103) | 62 (12) | 111 (56) | 64 (35) | — |
| T1-T2-T3 | 231 (97) | 207 (91) | 58 (11) | 97 (48) | 52 (32) | — |

### ICV Extraction

ICV extraction was performed using a self-written C++ pipeline based on the Minc-toolkit (version 1.0.08) for MRI image analysis (Vincent et al., 2016). We call the pipeline - IntracranialVolume (see https://www.neuromri.nl/2019/03/25/intracranial-volume/ to download the pipeline). For reasons of convenience, the pipeline was further wrapped using a python script within the Fastr environment (Achterberg et al., 2016) (see https://gitlab.com/bbmri/pipelines/fastr-resources). Similar to Buckner at al. (Buckner et al., 2004), our algorithm for ICV extraction makes use of several consecutive linear registrations steps (using the Minctracc program), followed by three non-linear registration steps (also using the Minctracc program). Such algorithm is a new implementation of a previously applied algorithm for ICV extraction used by our research group (see, e.g., (Scheewe et al., 2013)) that is of based on a clear segmentation rationale (Collins et al., 1995). In each registration instance, two MRI images are registered to an average MRI scan that was produced in the previous registration cycle. The first scan that is registered to the averaged scan is the T1w scan of the individual under consideration. The second scan that is registered to the same averaged scan is a model T1w MRI image that was created in our center by averaging several tens of T1w scans of different individuals. From the final registration matrices of the model MRI image and the subject scan to the averaged scan, one can calculate the transformation from the model T1w MRI image to the subject scan. Originally, adjacent to the T1w model MRI image, we also created an ICV mask, for that model brain, that was manually edited to assure the best correspondence between the model and its ICV mask. Applying the transformation that was calculated between the model MRI image and the individual scan to the model ICV mask, one obtains a new ICV mask for that individual. The total ICV of that subject is then merely the total number of voxels in that mask multiplied by the voxel volume. Note that, in principle, the dura matter is continuous between the brain and the spinal cord. Thus, it is hard to tell where exactly the brain ‘ends’ at its axial lower part. We have chosen to arbitrarily define the bottom part of the brain as the level below the cerebellum and the medulla.

We have adapted a semi-automatic procedure for the ICV extraction. The procedure amounts to running the ICV extraction pipeline with its default setting over a T1w MRI scan of a specific individual followed by manual inspection of an output figure file. The output figure file contains several sections of the axial, sagittal, and coronal directions of the brain overlaid by the outline of the ICV mask that was produced by the pipeline. If the ICV outer outline did not mark the real outline of the ICV, as judged by the first author, we have tried to run the ICV extraction tool with alternative flags to the Mintracc program. We have repeated this process several times until a satisfactory ICV mask was obtained, or else, it was decided that the pipeline is unable to produce an adequate ICV mask for that individual. The semi-automatic procedure was done blinded to participant age, gender, group status, or any other confounder.

In principle, in cases where the ICV mask was inadequate, two flags could be adjusted during the linear registration phase, and two could be adjusted during the non-linear registration phase to produce a better ICV mask. The two flags that could be adjusted during the linear registration phase are: (i) blurring the scans before the registration using the Mincblur command (usually, a kernel of 4-12 voxels helped), or (ii) not using the -est_translations flag of the Minctracc program. The two flags that can be adjusted during the non-linear registration steps are (i) setting the weighting factor for Minctracc optimization to a value of 0.01-0.4 (default 1.0); and (ii) setting the -similarity_cost_ratio flag of Minctracc to a value of 0.5-0.6 (default 0.3). Another measure that can help to achieve an appropriate ICV mask is to remove a large part of the neck in the scan of that individual, (as our model brain does not contain a neck moiety). In most of the cases, unsuccessful linear registration resulted in an ICV mask that is entirely unrelated to reality or that lacks a large portion of the cerebellum. Unsuccessful non-linear registration resulted in a part of the occipital region of the brain, or a small portion of the cerebellum, not being included in the ICV mask. In the majority of the cases (>90%), we obtained representative ICV masks after one or several rounds of applying the ICV extraction algorithm. However, in 3-9% of the images (depending on the time point), we were not able to obtain adequate ICV masks even after several rounds of running the ICV extraction tool. We have discarded these images form our analysis. Altogether, we extracted the ICV from 1143 T1 scans across all three data points that sum up to 94% of the available data. Statistical data regarding the number of scans that were used for the analysis out of the total number of scans in the cohort appear in Table 1.

We would like to stress that the re-iterative procedure of using different program registration flags did not result in an ICV bias, but was merely a technical procedure to assure that the ICV masks were adequate.

### Data Analysis

Data analysis was done within the statistical computing and graphics environment R (Team, 2018) using the packages: ggplot2 (Wickham, 2016), Matrix (Bates and Maechler, 2018), smoother (Hamilton, 2015), data.table (Dowle and Arun, 2018), matrixStats (Bengtsson, 2018), caTools (Tuszynski, 2018), gam (Hastie, 2018), and haven (Wickham and Miller, 2018).

Since our research hypothesis was that the expected longitudinal ICV changes will be extremely small and hard to detect (as the results indeed shows), and since we wanted to obtain high confidence in our results, we were not contented with analyzing the data using only one analysis method (even if the results obtained using this analysis method were statistically significant). Instead, as is described below, we used three complementary methods to analyze the data. These methods are detailed below.

In addition, to prevent any influence of residual physical growth for young subjects, we took into considerations only subjects with an average age between measurements larger than 20 years (unless otherwise specified). Moreover, in all cases, we did not detect any apparent differences between the longitudinal ICV behavior for females and males. Hence, in all cases, we analyzed the data for females and males separately, and together. In all cases, analyzing the data of females and males together resulted in an improved type-I error p-value for a fit to the data. These observations suggest that, in many cases, the limits on obtaining statistically significant p-values are mainly related to low statistical power in the face of a broad spread of the data. We report the statistical characteristics of the fits for females, males, and females and males together.

Note that, even after all precautions that we took, including the manual control of the ICV masks, our results still include outliers. These outliers could be identified by noting the existence of two groups in our results: one with very small longitudinal changes, and one with somewhat larger longitudinal changes (positive and negative). In other words, the distribution of longitudinal ICV changes included long tails from both sides of the distribution. The existence of the group with the more considerable absolute value longitudinal changes curtails our ability to reach statistical significant longitudinal ICV changes, and we consider them as outliers. To account for the existence of these outliers, we adopted a consistency filtering mechanism. We have used these consistency analysis methods to filter out cases with relatively large longitudinal ICV changes as we believe that individuals presenting these large absolute value longitudinal ICV changes are artifacts that do not represent real longitudinal ICV masks and consequently do not represent real ICV changes. The exact filtering mechanisms that were applied in each analysis case are detailed below.

### Differences between ICV values at different waves of the study

We calculated the ICV differences between different waves of the study (ΔICV_Tx,Ty_) using the formula:

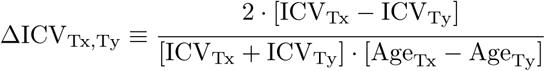

where Tx and Ty are different waves of the study. Ty - first, second or third wave of the study, Tx - second or third wave of the study. ICV_Tx_ is the calculated ICV at the first, second or third scan and Age_Tx_ is the corresponding age of the individual in years at the time of the scan.

The individual values of ΔICV_Tx,Ty_ where used to obtain a longitudinal trajectories of ICV change by fitting the set of {ΔICV_Tx,Ty_} results of the whole cohort to a straight line as a function of {Age_Tx,Ty_ ≡ 0.5 · [Age_Tx_ + Age_Ty_]}, where {…} represents a set of individual results.

To filter out points with large positive and negative ΔICV_Tx,Ty_ values, we have ordered the {ΔICV_Tx,Ty_} set according to their values, fitted the central part of the ordered ΔICV_TxTy_ sequence to a straight line for males and females separately, and maintained only individuals with longitudinal ICV change that deviated from the fitted line by less than 0.2 %/year (except for the case of females for ΔICV_T2,T1_ where the allowed deviation was of 0.25 %/year.) This method resulted in a reduced {ΔICV_Tx,Ty_} dataset that lacks the positive and negative heavy-tails parts of the {ΔICV_Tx,Ty_} set.

### Individual fit based analysis

Our longitudinal cohort contains three consecutive measurements. Since several error factors can confound MRI results and since computational measures of brain volumes from structural MRI scans are known to add additional error factors, we turned to minimize these effects by making use of the full three measurements that are in our hand. Thus, instead of taking into account only two ICV measurements between different waves and calculating ΔICV_Tx,Ty_, we have fitted the measured ICV as a function of the age to a straight line for each subject separately using the R function lm. In other words, for each individual (i), we fitted its ICV (ICV_Individual i_) to a straight line using the formula:

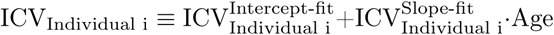

where 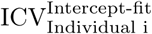 and 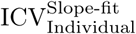 are the intercept and incline of the ICV data for that individual. The fitting was over three data points for those individuals which the ICV data existed for three consecutive measurements, or over two measurements in cases that only two ICV values existed.

This procedure resulted in a degenerate fitting of only two measurements for 30 cases and a meaningful fitting of three consecutive ICV measurements in 207 cases (87% of the cases, see Table 1). Next, we plotted the set of slopes for all the individual fits 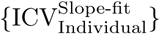 as a function of the average age of the subjects between these scans 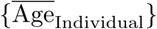.

Finally, we have used the 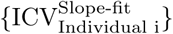 values for all individuals to obtain a global longitudinal ICV trajectory by fitting the 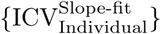 set as a function of 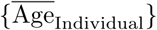 to a straight line.

To obtain a reduced dataset for 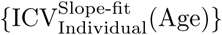, we have adopted the same approach as in the ΔICV_Tx,Ty_ case. By ordering the set of 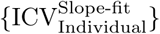 results according to their values for females and males separately and fitting the central part of the ordered datasets to straight lines, we created a reduced dataset that excludes the long tails in the data. The reduced 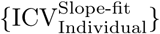 datasets discards individuals that their 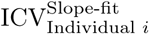 deviated from the linear fit to the central part of the ordered 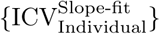 datasets (for females or for males) by more than 3 ml/year. Finally, to obtain a predicted longitudinal ICV trajectory for the reduced dataset case, we fitted the reduced dataset 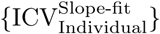 as a function of 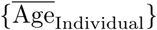 to a straight line.

### Linear mixed models fitting

The measured ICV dataset contains data points that are not entirely independent of each other, thus violating an essential assumption of a mathematical fit theory as was done for 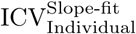 and ΔICV_Tx,Ty_ cases. The measurements in our cohort are not independent of each other in two respects. First, by the nature of the longitudinal design, they contain repeated measurements of the same individuals several times. Second, our cohort was designed to include conjoint family members (parents and siblings). As it is known that the ICV is influenced by genetic factors (Adams et al., 2016), the ICVs of related family members are not independent of each other.

To take these effects into account, we have adopted a third layer of analysis using a linear mixed model approach (LMM). LMMs are a general framework to fit continuous outcome variables in which the residuals may not be independent, as in the case of a longitudinal design (Bates et al., 2015). In principle, an LMM model is composed of a series of global effects that influence the whole dataset and a series of effects that are called ‘random’ and that are unique to each group of measurements. In that sense, an LMM is a sort of global analysis that profoundly reduced the number of degrees of freedom of the analysis relative to the ΔICV_Tx,Ty_ and 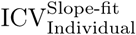 analyses.

To utilize the LMM approach, we have analyzed the ICV data using the R environment lme4 packaged (Bates et al., 2015) for all the participants in our cohort for which we were able to extract two or three measurements. That is, we have included in the LMM analysis all individuals that had their ICV measured at the first and second wave of the study, at the first and third wave of the study, at the second and third wave of the study, or at all three waves. As a primary LMM analysis model, we have constructed an LMM model with global effects of age and sex and random uncorrelated effects of intercept and age-dependent ICV for each participant as well as for all the member of a specific family. In the lme4 package notation the model that we have used is: ICV∼Age+Sex+(1+Age||Family:Subject) for the non-reduced complete ICV dataset. In this notation (1+x||y:z) means an individual z dependent random variable nested in factor y where the intercept (1) is not correlated with the x effect. Next, we used the predicted ICV values for each participant that were obtained from the LMM analysis at the corresponding age when the MRI scans were performed to calculate a predicted set of age-dependent ICV changes 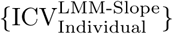, and fitted this set as a function of 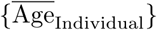 to a straight line. From this LMM-dependent straight line, one can predict the LMM based trajectory of longitudinal ICV changes.

Note that in many cases, when analyzing a longitudinal data using the framework of LMM, it is possible to study both the cross-sectional and longitudinal effects simultaneously by incorporating into the LMM the mean of the variable as an inter-subject effect. However, this is not the case when the cross-sectional inter-subjective differences are much larger than the longitudinal ones, as is the case here (see chapter 9 in (Verbeke and Molenberghs, 2000)). Indeed, incorporating the Mean ICV as a fixed effect resulted in a model that did not converge. Thus, we did not include the mean ICV term in the linear mixed model.

To further check for the accuracy of our LMM model, we have constructed a series of alternative LMM models and checked the statistical significance of an ANOVA comparison between the primary LMM model and the alternative models. Adding to the primary LMM model the group status of each subject (i.e., whether it belongs to the control group, the sibling group, or to the group of people that were diagnosed with schizophrenia) resulted in a highly insignificant result (ANOVA p-value=0.984). Also, adding a global group status*Age factor is not significant (ANOVA p-value=0.788). Similarly, adding to the LMM model the average measured IQ of the cohort’s participants as a random confounder effect resulted in a highly insignificant result (ANOVA p-value=1). Also, adding the identification of the MRI machine (one out of two MRI machines that were used for the GROUP cohort and a third identification code for cases where the MRI machine is unknown) resulted in an insignificant result (ANOVA p-value=0.551). By contrast, removing the global effect of the sex status showed that the original LMM was significantly better (ANOVA p-value=2.2e-16). Similarly, removing the random effect of family membership showed that the original LMM was significantly better (ANOVA p-value=1.97e-09). Finally, we have checked whether removing the global age effect from the LMM model will worsen or improve the primary model. The ANOVA result for removing the global age effect was statistically marginal (p-value=0.0504), meaning that the model with the global age effect is marginally better. In light of this marginal p-value, we still have kept this effect in the LMM model analysis.

To obtain a reduced dataset filtered from long-tailed extreme-cases of longitudinal ICV changes for LMM analysis, we have calculated the difference between the maximum and minimum measured ICV values (ΔIC*V*_max,min_ ≡ICV_max_− ICV_min_) for individuals, ordered the results according to their values, and fitted the 200 cases with the smallest value of ICV_max,min_ for females and males together to a straight line. By setting a limit on the allowed deviation from this straight line (5 ml/year), we obtained a reduced dataset for the LMM analysis. We have used this reduced dataset to construct an LMM model with longitudinal cases that have only small measured ICV change. For the reduced dataset is Lme4 notation the primary model that we used was: ICV∼Sex+MRI-Machine+(1+Age||Family:Subject).

Similar to the complete LMM dataset case, we constructed a series of LMM models and checked their applicability based on an ANOVA comparison. Importantly, unlike in the complete dataset case, for the reduced dataset case, the inclusion of the MRI machine code that was used to acquire the scans resulted in an improved model (ANOVA p-value 0.026). However, including the global age effect did not make the LMM model better (ANOVA p-value 0.146). By contrast, excluding the sex identification factor or family status, resulted in a statistically significant worse LMM model (ANOVA p-values - 2.2e-16 and 1.0e-07, respectively). Similarly, including the average IQ, or the patient status did not improve the LMM model (ANOVA p-values - 0.8857 and 1, respectively). Also, adding a global group status*Age factor resulted is not significant (ANOVA p-value=0.561). Thus, we chose an LMM model that includes the sex and the MRI-machine code as global effects, and the family, subject code, and age as random effects (but without a global age effect).

Finally, to obtain the predicted longitudinal ICV change trajectory, we have fitted the 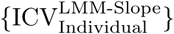 set as a function of 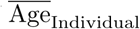 for females and males separately or together to a straight line,

To check the existence of higher ordered terms in the longitudinal ICV trajectories, we have applied two analysis methods. First, we studied the behavior of the 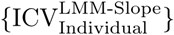 data by grouping males and females together and calculating a running average age dependency behavior of the 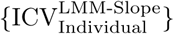 (minus the global age effect for the complete dataset). Second, in a more statistically controlled manner, we have used the Generalized Additive Model package of R (‘gam’). A generalized additive model (GAM) is a sort of spline fit that can include one or a higher number of degrees of freedom. In other words, it is a linear predictor model involving a sum of smooth functions of covariates (Hastie and Tibshirani, 1986). We have fitted the 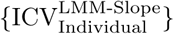 database as a function of age to a GAM model with one, two, and three degrees of freedoms.

### FreeSurfer Analysis

Alternative ICV measure to our home-based ICV extraction method was obtained by running the T1-weighted scans through the FreeSurfer recon-all pipeline (version 5.1.0) and obtaining the results from the aseg.stats file in the FreeSurfer directory as is explained in the FreeSurfer manual (see freesurfer manual).

### ICV as a function of IQ

We studied the relationship between ICV and the average IQ of individuals over the three waves of the study by plotting the values of the extracted ICV and the measured IQ and fitting the results to straight lines for females and males separately, as well as for the control group, siblings group, and the group of people that were diagnosed with schizophrenia separately (see the Supporting Text of the SI for more details.)

### Calculation of CSF, Gray Matter and White Matter

CSF, gray-matter (GM), white-matter (WM) were calculated by applying the ICV that was calculated for each subject as a mask on the T1w scan followed by separation of the CSF, GM, and WM using the FMRIB Software Library v6.0 FSL function FAST (Zhang et al. (2001)). Next, we extracted the volume values for these tissues (and the CSF) using the FSL function fslstats.

Relationship between the ICV, the CSF, the GM, and the WM were analyzed by defining ΔCSF_*T*3 .*T*1_, ΔGM_*T*3, *T*1_, and ΔWM_*T*3, *T*1_ similar to ΔICV_*T*3, *T*1_ and (a) calculating the Pearson correlation-coefficients (P_*Corr*_) using the statistical computing and graphics environment R function cor.test and using a filtration approach based on ΔICV_*T*3, *T*1_ as is described above; (b) fitting the ratio of ΔCSF_*T*3 .*T*1_ to ΔICV_*T*3, *T*1_ as function of age to a straight line and filtering the results based on the ratio of ΔCSF_*T*3 .*T*1_ to ΔICV_*T*3, *T*1_ to obtain an age-dependent reduced dataset (similar approach was used for the gray matter and the white matter); (c) fitting the relationship between the CSF (or the GM, or the WM) and the ICV for each individual separately using the equation:

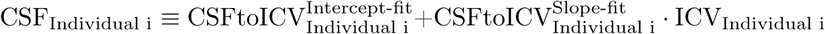

and fitting 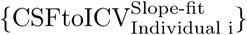 as a function of the average age between the scans to a linear function. To obtain the age-dependent relationships of the 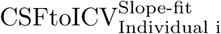 for a reduced dataset where the results with relatively large value of 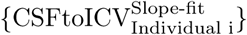 where filtered-out we used a similar analysis as is described above. Finally, we fitted the reduced 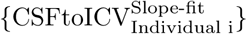 dataset to a linear function. A similar approach was used to analyze the 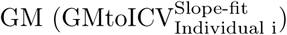 and the 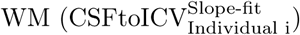.

### Cross-sectional Analysis

To check the cross-sectional behavior of the data, we fitted the extracted ICV set for each wave of the study separately for females and males as a function of the age of the individual (in years) at the time of the scan acquisition to a straight line. Next, we subtracted from the ICV of all females the intercept of the females fit (and similarly for males). We used the subtracted values to obtain a combined set of females and males. Finally, we fitted this combined set to a straight line as a function of age. We also repeated a similar analysis for the height of individuals for the first time they were scanned as a function of age.

To check the existence of higher-order terms in the cross-sectional results, we fitted the data for males and females separately to a general additive model (GAM) with one, two, or three degrees of freedoms. In addition, we also applied a running average for females and males separately over the data with a window of six years (similar to the period between the first and last waves of the study) and fitted the results to a straight line.

### Figures preparation

Figures and graphs for this article were prepared using the programs: SciDAVis (RRID:SCR_014643), Inkscape (RRID:SCR_014479), QtiPlot, and GIMP (RRID:SCR_003182).

## Results

### General ICV characteristics

Fig. 1 shows an example of several sections from a T1w MRI scan with an overlay of the ICV-mask outline that was obtained for that scan for one case of the GROUP cohort. As can be seen, the ICV mask that was produced by our computational tool encompasses the brain and the CSF and omits the two large dura mater reflections, namely the tentorium cerebelli and the falx cerebri (Adeeb et al., 2012). Thus, the computationally-obtained ICV-mask provides a direct measure of the conjointly total volumes of the brain and the CSF.

**Figure 1:**
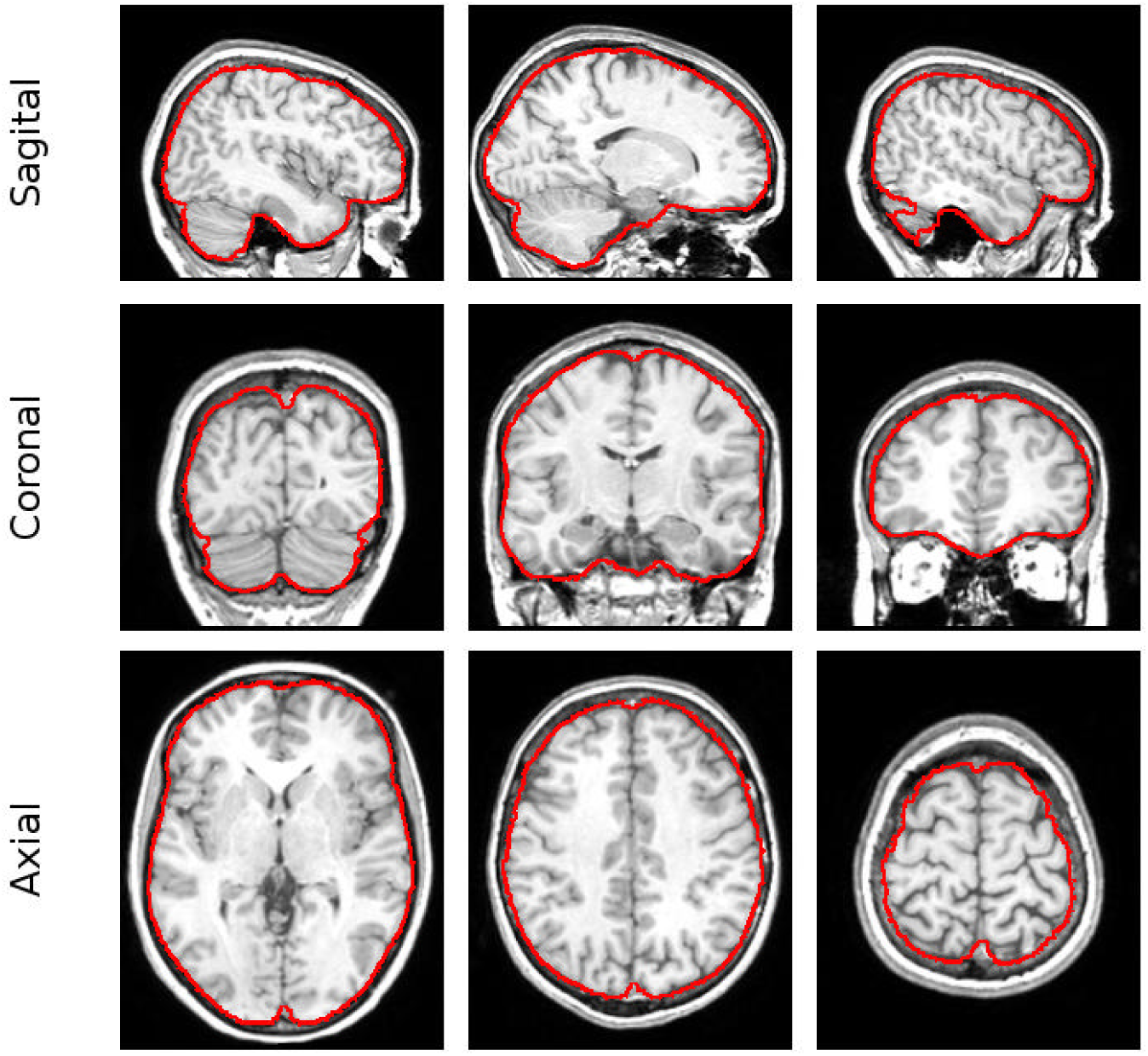
Extracted ICV. An example of an extracted ICV mask. Outline of the ICV mask (red line) for three sagittal, coronal, and axial slices superimposed on the original MRI scan. The ICV mask was calculated using our semi-automatic IntracranialVolume extraction tool.

A histogram plot of the extracted ICV values at the first time point shows a normal-like distribution (see Fig. S1a of the Supporting Information (SI)). The average (and standard deviations) of ICV value for females is 1.4±0.11 liters and for males 1.6 ± 0.13 liters (females/males ratio =0.875. For comparison see (Ruigrok et al., 2014)). The distributions of the ICV values at the second and third scanning periods show similar characteristics, and the average (±s.d.) ICV value stays constant (see Fig. S1b,c of the SI).

Since we have used a home-built-based method to extract the ICV, we wanted to validate our ICV extraction method using some external measures (unrelated to aging) before moving to study the relationship between ICV and age. We achieved this goal through a two-fold approach. First, we showed that cases that we were not able to extract the ICV are not statistically significant different from those that were successfully extracted (p-values for two-sided Wilcoxon Rank test for age distributions at the first, second and third waves of the study = 0.138, 0.303, 0.528; p-values for *χ*^2^ test for gender at the first, second and third waves of the study = 0.0094, 0.679, 0.116; p-values for *χ*^2^ test for patients, siblings, and control groups at the first, second and third waves of the study = 0.935, 0.673, 0.088; Bonferroni corrected statistically significant level - 0.05/9 = 0.0056). Second, we show that we were able to recapture the known characteristics of the ICV-IQ relationship for most of the cases (see Fig. S2 of the SI and supporting text for further details).

### Longitudinal ICV changes are extremely small

Fig. S3 of the SI shows the longitudinal trajectories of ICV values for all individuals in our cohort with two or more consecutive successful extraction of the ICV masks. Figures S4-S6 show the corresponding Spaghetti plots for the participants as a function of age, sex, and the scanner identification relationship of each two consecutive scans. Overall, no substantial changes occur in the individual ICV values from young adulthood to the sixth decade of life.

To assess longitudinal changes in the ICV, we have calculated ΔICV_T3,T1_ (see Methods section). A histogram of the results shows that ΔICV_T3,T1_ presents a distribution with average and standard error (s.e.) of the longitudinal ICV change equal to −3 · 10^−2^ ± 2.7 · 10^−2^ [%/year] for females and−3 · 10^−3^ ± 2 · 10^−2^ [%/year] for males (see Fig. 2a). In both cases, the ΔICV distributions do not follow a normal distribution. Instead, the ΔICV distributions have long tails from both sides of the distributions. Thus, this result suggests that if longitudinal changes in the ICV between early adulthood to the sixth decade of life exist, they are minimal and amount to a value of the order of 0.1 [‰/year] of the total ICV. Note that the age distributions for both females and males in our cohort are skewed toward people that are younger than 30 years (see Fig. S7 of the supporting information). Thus, the age distributions might mask the true trajectories of ICV changes as a function of age as they are assessed using the ΔICV distributions.

**Figure 2:**
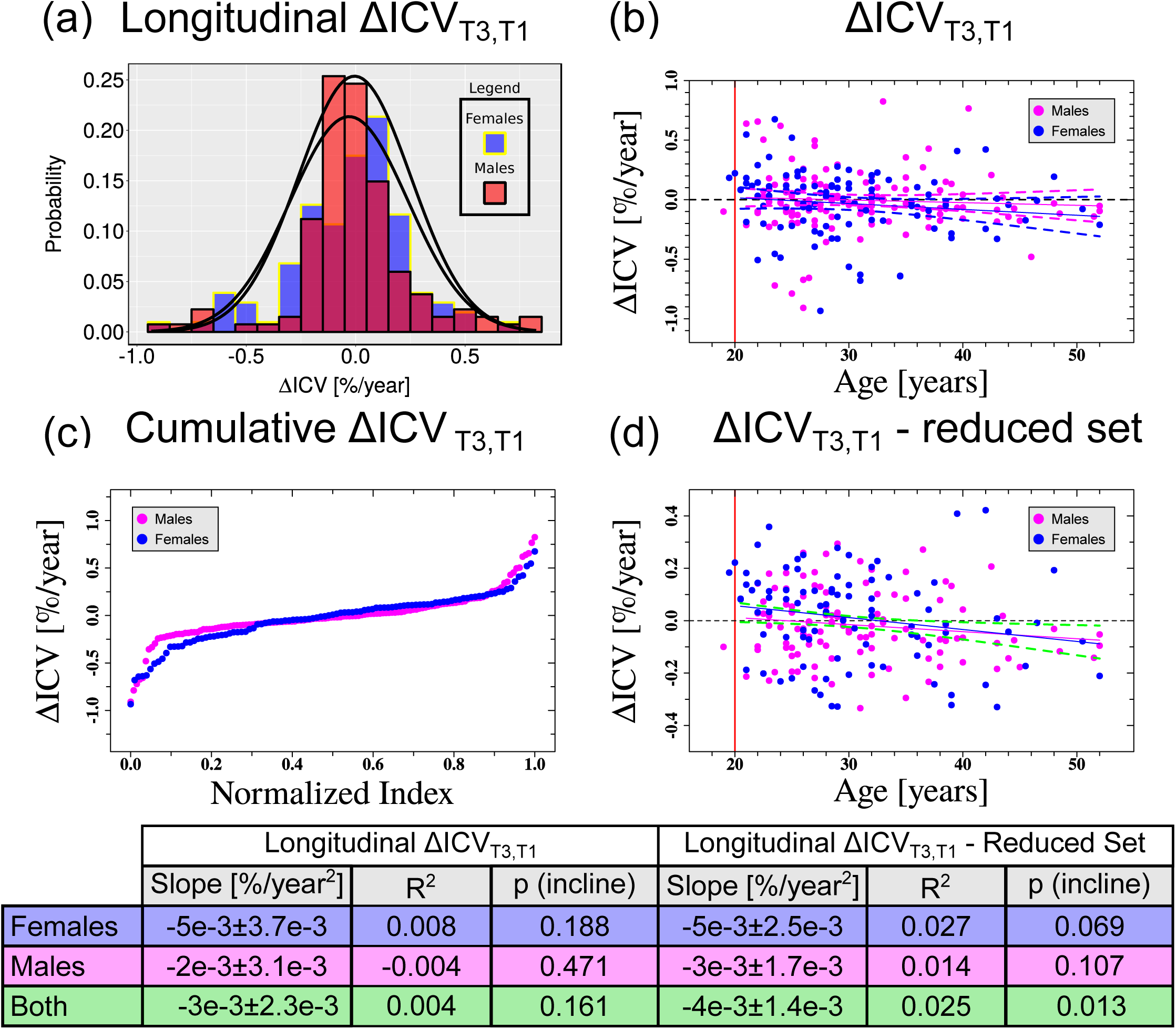
Changes in the ICV as a function of age. (a) Distribution of the measured ΔICV_T3,T1_ values for males (red) and females (blue). (b) ΔICV_T3,T1_ as a function of Age_T3,T1_. Each dot represents a calculated value for a specific individual in our dataset. Blue and pink lines are fits of the data to linear functions. Vertical red line represents the minimum age for inclusion in the fits (dots left to the line were not included in the fitting process). Dashed lines represent 95% confidence intervals of the fits. (c) Cumulative plot of ΔICV_T3,T1_ as a function of a normalized index between 0 and 1. (d) Same as (b) for the reduced {ΔICV_T3,T1_} dataset based on the middle part of the cumulative {ΔICV_T3,T1_} result. See also Fig. S9 of the SI for the precise selection criterion of the reduced {ΔICV_T3,T1_} dataset. In (b)-(d) blue color is for females and pink for males. In (d) dashed green lines are 95% confidence intervals of the fit for both females and males. In (d), for clarity, the fit itself for females and males together is not shown. The table below the graphs shows the fitting-values for the cases of females, males, and for the females and males together. *R*^2^-goodness of fit. p-value - statistical significance of the slope parameter of the fit. Error values - ±s.e.

### ICV differences between the first and last scanning waves

To assess the existence of small longitudinal ICV changes, we plotted {ΔICV_T3,T1_} as a function of {Age_T3,T1_}. The results of the analysis are shown in Fig. 2b. We fitted {ΔICV_T3,T1_} as a function of {Age_T3,T1_} for males and females separately. In both cases, the fitted lines had a negative slope. However, in both cases, the fits did not reach the statistically significant limit (see the table in Fig. 2). Inspection of the quantile-quantile plots (Q-Q plots) suggested the existence heavy-tail points in the residuals’ distribution (see Fig. S8a of the SI). Moreover, both fits of ΔICV_T3,T1_ may show signs of heteroskedasticity (see Fig. S8b of the SI). Taken together, these observations suggest problems with the linear models that were used for these fits.

Note that in Fig. 2b several points seems to be localized far away from the main pile of points. We were interested in checking if the existence of these points concealed a statistically significant result. To test this hypothesis, we have applied a filtering procedure to reject cases with large absolute value of ΔICV_T3,T1_ (see Fig. 2c). In this plot, the existence of two groups of heavy-tail points, one with a substantial negative ΔICV_T3,T1_ and one with a substantial positive ΔICV_T3,T1_, are clearly observed for both females and males. We used these data to create a reduced dataset (see Fig. S9a,b of the SI for the exact limits that were imposed on the ΔICV_T3,T1_ dataset to create the reduced dataset and the Materials and Methods section for details).

Next, we plotted {ΔICV_T3,T1_} as a function {Age_T3,T1_} for the reduced dataset and fitted the data to linear functions for females and males separately (see Fig. 2d). In this case, there was a large improvement of in the p-values of the linear fits. The p-value for the slope for the females fit reached the lesser stringent statistical-significance limit of 10% (p-value=0.069), but the slope p-value for males was above the statistical-significance limit (p-value=0.107). We could further reduce the slopes p-values by setting a somewhat stricter criterion for obtaining the ΔICV_T3,T1_ reduced dataset. With the stricter criterion, the slope p-value of the linear fit for females reaches the 0.05 statistical-significance limit, and the slope p-value for males reaches the 0.1 statistical-significance limit (data not shown). The fact that for the reduced ΔICV_T3,T1_ dataset the Q-Q plots for both females and males were linear (see Fig. S8c of the SI) and that the residual plots did not show signs of heteroskedasticity (see Fig. S8d of the SI) provides an additional layer of sanity-check to show that the process for obtaining the reduced dataset did not introduce a bias to the analysis.

When grouping both females and males together, the fit showed a negative slope, and the p-value was statistically significant (p-value=0.013, see table in Fig. 2). In addition, the Q-Q plot for the combined females and males fit showed a linear behavior, and no heteroskedasticity was observed (see Fig. S8e,f of the SI). A ‘leave-one-out’ analysis of the combined dataset of females and males together showed that the maximum influence of any point of the linear fit slope is 13%, well within the s.e. of the slope. Thus, the relationship that we detect between ΔICV_T3,T1_ and Age_T3,T1_ is not the result of changes in the ICV of any single individual.

It is interesting to note that similar results to these of ΔICV_T3,T1_ were also obtained for ΔICV_T2,T1_, but not for ΔICV_T3,T2_. For the ΔICV_T3,T2_ case, the results did not show statistically significant longitudinal ICV change. We believe that the reason for this discrepancy may be related to a lower statistical power as results of a smaller number of subjects in the ΔICV_T3,T2_ dataset relative to the ΔICV_T2,T1_ one in the face of a low *R*^2^ of the age dependency (See Fig. S10 of the SI and the Supporting Text of SI for further details).

Taken together, our analysis suggests that the ICV continues to grow in young adulthood. Later on, the ICV starts to shrink from the middle of the fourth decade of life (when ΔICV_T3,T1_=0; Age ≈ 34 years) and continues in this tendency well into the sixth decade of life (most probably also in the years after). Moreover, our data suggest that the decrease in ICV is not constant but accelerate with aging during the young and middle adulthood period. However, the size of ICV changes during this period is minimal, and even at the age of 55 (the end age of our analysis) amounts on average to less than 0.1 %/year.

### ICV trajectory based on all three measurements

The results of the Individual fit based analysis for all three measurements are shown in Fig. 3a. Similar to the analysis based on the ICV difference between different waves of the study, fitting 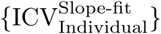 as a function of 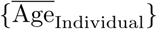 resulted in a non-statistically significant fit with a negative slope for both females and males. Grouping females and males together, improved the p-value of the fit a bit but did not cause it to reach statistical significance. Also, similar to the previous analysis method, the residuals Q-Q plots showed the existence of large tails (see Fig. S11a of the SI), and the residual vs. fitted values plot may have hinted to some heteroskedasticity.

**Figure 3:**
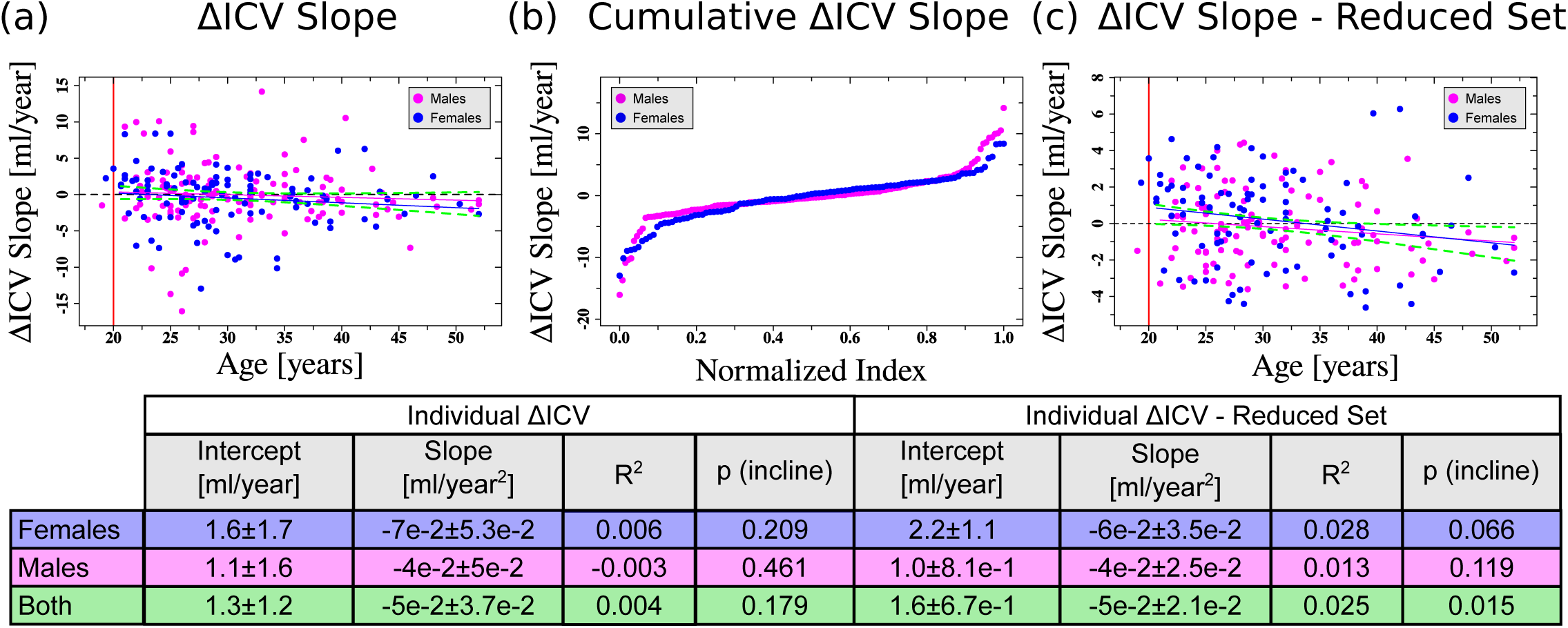
Individual longitudinal fits of ICV as a function of age. (a) Values of the slopes of the individual ICV fits as a function of the average age at which the scans were made. Each dot represents the age-dependent slope for an individual as was obtained from a linear model fitting in R to two or three (when available) consecutive ICV measurements. Lines are fits of the data to linear functions. Vertical red line represents the minimum age for inclusion in the fits. (b) Cumulative plot of 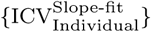 as a function of a normalized index between 0 and 1. (c) Same as (a) for the reduced ICV{age} dataset. See also Fig. S12 of the SI for the precise selection criterion of the reduced ICV{age} dataset. In (a)-(c) blue color is for females and pink for males. Dashed green lines are 95% confidence intervals of the fit for both females and males. The table below the graphs shows the fitting-values for the cases of females, males, and for the females and males together. For clarity, the fits for females and males together are not shown. *R*^2^- goodness of fit. p-value - statistical significance of the slope parameter of the fit. Error values - ±s.e.

To further analyze our data, we have created a reduced dataset by filtering out cases with a large absolute value of 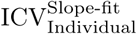 (See Fig. 3b, Fig. S12a,b of the SI and the Materials and Methods section). For the reduced dataset of 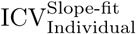, we have observed a substantial decrease in the p-value of the fit to 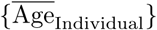 for both females and males (see Fig. 3c and the table therein). In that case, the females case reached the 0.1 statistical significance limit. The p-value for males was, however, above the statistical significance limit. Inspection of the residuals Q-Q plots (see Fig. S11c of the SI) showed good linearity of the fits for females and males, and the residuals vs. fitted values plots (Fig. S11d of the SI) showed no apparent indications of heteroskedasticity.

After grouping together females and males, the p-value of the slope of the fit was statistically significant (see table in Fig. 3), the residuals Q-Q plot was linear, and signed of heteroskedasticity were not observed (see Fig. S11e,f of the SI). These results provide additional support to our conclusion that the ICV shows a small but accelerated change during young and middle adulthood.

### Linear mixed model for ICV trajectory

To take into account the dependency in our data (repeated measurements of the same individual and family relatedness) we analyzed our data using an LMM approach. The results of the 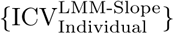 analysis predictions minus the global Age effect of the LMM model as a function of 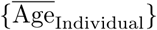 are shown in Fig. 4a. The statistical characteristics and the fitting-values for the corresponding fits are shown in the adjoined table to Fig. 4. From the results of the LMM model, we have obtained statistically significant age-dependent ICV change for females and males, as well as for females and males together (see table in Fig. 4).

**Figure 4:**
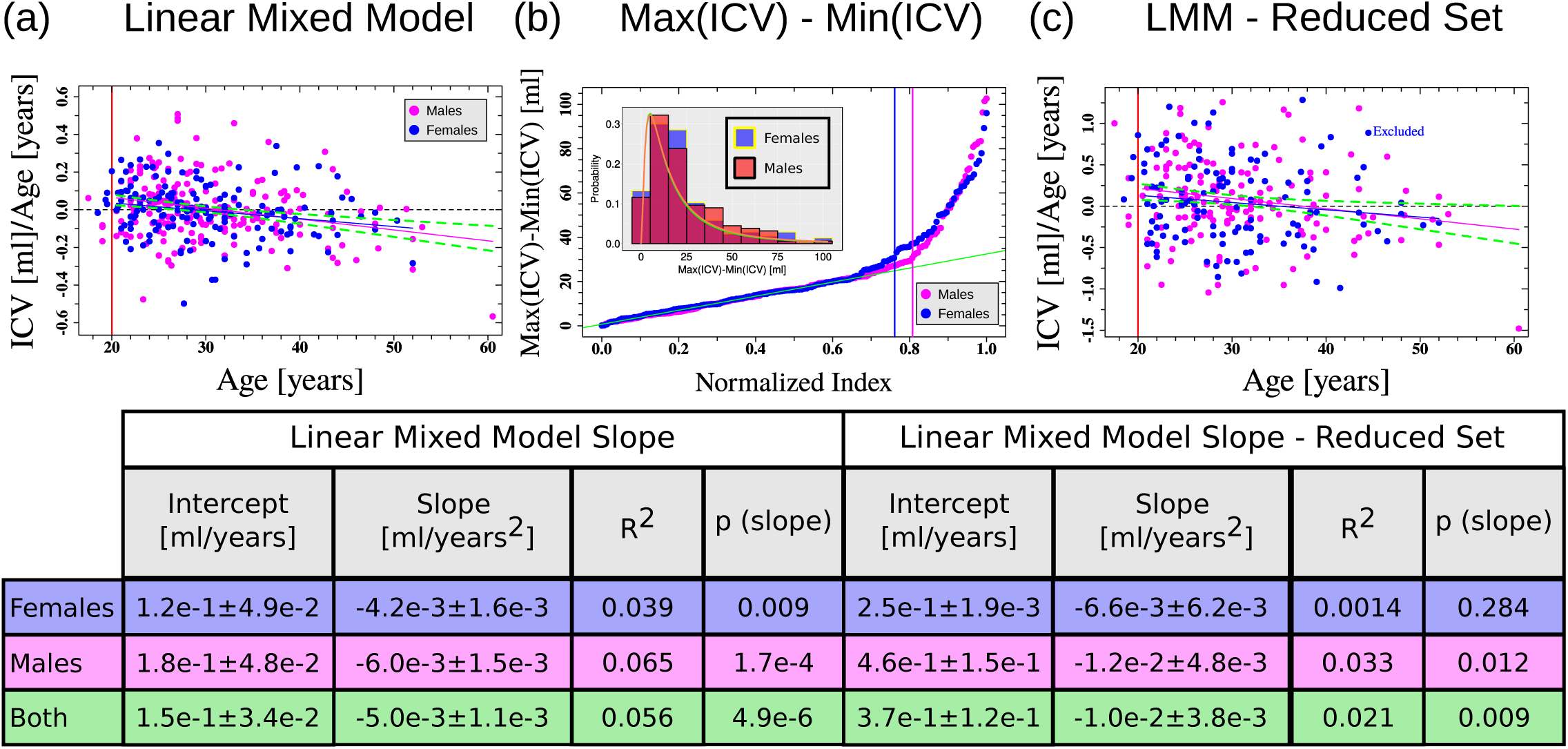
Linear mixed models for ICV as a function of age. (a) Predicted 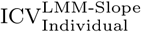 as a function of age. Each dot represents the age-dependent ICV slope for a specific individual. Lines are fits of the data to linear functions. Red line represents the minimum age for inclusion in the fits. (b) Cumulative measured individual ICV_max_ − ICV_min_ as a function of a normalized index between 0 and 1. Green line - fit of the 200 subjects with the smallest ICV_max_ − ICV_min_ (for both females and males) to a straight line. Vertical lines - limits of the normalized index criterion for inclusion in the linear mixed model reduced datasets for females and males. Inset - Distributions of ICV_max_ − ICV_min_. Green and Orange lines are fits of the distribution data to log-normal distributions for males and females respectively. (c) Same as (a) for the reduced 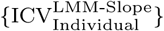 dataset. In (a)-(c) blue color is for females and pink for males. Dashed green lines are 95% confidence intervals of the fit for both females and males. The table below the graphs shows the fitting-values for the cases of females, males, and for the females and males together. For clarity, the fits for females and males together are not shown. In (c), ‘Excluded’ represents the females individual that most influenced the fit. Excluding it did not cause the fit to reach the statistical significance limit. Data in the table includes this individual. *R*^2^- goodness of fit. p-value - statistical significance of the slope parameter of the fit. Error values - ±s.e.

We were further interested in checking whether higher order terms in the longitudinal ICV trajectories exist behind a linear age-dependent change. A running average window analysis of the whole 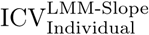 database did not show a clear-cut indication for the existence of higher order terms in the longitudinal ICV trajectories (see Fig. S13a of the SI). Similarly, an ANOVA result of the GAM models of one and two degrees of freedom had a p-value of 0.068, and an ANOVA result of the GAM models of one and three degrees of freedom had a p-value of 0.097. Thus, we did not obtain a statistical indication for the existence of higher order terms in the longitudinal ICV trajectories.

It should be noted, however, that the nature of the LMM as a global fit with small number of degrees of freedom (eight in our case), can cause it to be sensitive to the existence of extreme cases in the dataset that is fitted. To check this possibility, we again inspected the residuals of the LMM analysis. Indeed, the residuals Q-Q plot of the LMM model shows the existence of large non-linearities though it still has equal statistical variances across the whole fitting range (See Fig. S14a,b of the SI). Interestingly, the existence of large non-linearities in the LMM model, did not cause similar non-linearities or heteroskedasticity in the 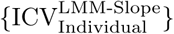 fit analysis for females and males (see Fig. S14c,d of the SI). Nor did it caused this effects to occur in the 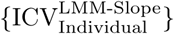 fit of both females and males (see Fig. S14e,f of the SI). Still, the residual analysis of the LMM model suggests that the extreme cases that were included in the LMM analysis may have skewed its results.

To account for these extreme cases, we filtered them out, constructed a reduced dataset, and again run a LMM analysis (see the Materials and Methods section). The results of the LMM analysis of the reduced dataset showed no signed of heteroskedasticity or non-linearity (see Fig. S15a,b of the SI). We have used the LMM model of the reduced dataset to calculate the slope of the ICV age dependency for individuals and to obtain a reduced dataset longitudinal ICV trajectory. The behavior of the reduced 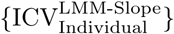 values as a function of age is shown in Fig. 4c. As expected, as a result of filtering out the individuals with large ICV_max,min_, we obtained a broader spread of the predicted LMM age-dependent slopes. A linear fit of the predicted individuals LMM age-dependent ICV changes as a function of age resulted in a statistically significant linear behavior for males (see Fig. 4c and the table therein). However, for females, the fit was not statistically significant (p-value=0.283). Excluding the subject with the most substantial influence on the linear fit slope did not increase the statistical significance of the fit substantially (p-value=0.157 see Fig. 4c). Since the residuals Q-Q plot and the residuals vs. fitted values did not show signs of non-linearity or heteroskedasticity (see, Fig. S15c,d of the SI), the lack of statistical significance for the female group is probably connected to a lack of statistical power for the females group relative to the large males group in the cohort. Indeed, lumping together the females and males and fitting all the predicted ICV age-dependent changes for the reduced dataset LMM model still resulted in a statistically significant longitudinal age-dependent linear behavior of the ICV change (p-value=0.029; see table in Fig. 4). Moreover, there were no signs of non-linearity or heteroskedasticity in this fit (see Fig. S15e,f of the SI).

Results of running average over the 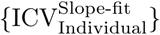 of the reduced dataset for both females and males together did not show signs of higher-order terms ICV acceleration above a linear one (see Fig. S13b of the SI). Similarly, a GAM analysis with one, two, and three degrees of freedom showed that one degree of freedom GAM model (meaning a linear one), is the most statistically significant (ANOVA p-values 0.1465 and 0.0816 for the one and two, and one and three degrees of freedom models, respectively).

Thus, consistent with the ΔICV_T3,T1_ and the 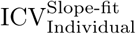 analyses, the LMM analyses, both for the complete ICV dataset and for the reduced one, showed an average linear accelerated change of the total ICV. Such linear accelerated ICV change results in a small but consistent net ICV reduction starting from the beginning of the fourth decade of life according to the complete dataset analysis or from the beginning of the fifth decade of life according to the reduced dataset analysis. It should be noted, however, that the LMM analyses predict a much smaller acceleration for the ICV change in comparison of to the ΔICV_T3,T1_ and the 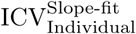 analyses. Yet, the critical finding of this research is that, on average, the ICV is not constant during young and middle adulthood.

### Analysis without patients

As mentioned in the Methods section, the cohort contains three groups (people that were diagnosed with schizophrenia, their relative, and a control group). A reservation regarding can be made regarding the inclusion of people that were diagnosed with schizophrenia in our analysis. These reservations might be related to differences in trajectories of brain tissue changes (but not necessarily is the skull) between people that were diagnosed with schizophrenia and control groups. We have tried to control for this factor by including the group status in the LMM analysis, which resulted in a high ANOVA p-value. To further check the possibility that the inclusion of people that were diagnosed with schizophrenia caused a bias in our results, we re-run the cross-sectional analysis while removing all members of this group. The longitudinal results for the three analysis methods are shown in Fig. S16-S18. In this case, for the linear-mixed-model analysis of the reduced dataset, we did not include the MRI machine status (ANOVA p-value 0.294). As can be seen, no substantial differences appear between the analyses with the people that were diagnosed with schizophrenia and the analyses without this group.

### FreeSurfer analysis

To obtain a further corroboration of our findings, we have recalculated ICV change using the FreeSurfer pipeline and reran the analyses. The results of the FreeSurfer eTIV estimation are shown in Fig. S19-S21. Again, this resulted in long tails with large ICV differences estimations (see Fig. S16b, S17b, and S18b). As can be seen from Fig. S19-S21, we were able to replicate our results for females, but not for males for ΔICV_T3,T1_, and 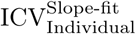. However, for the linear mixed model in the complete dataset, we also incorporated the MRI machine identification as this was statistically significant (ANOVA p-value 0.039). For the linear mixed model of the reduced dataset, we removed the MRI scanner identification as this effect was not significant (ANOVA p-value 0.21). Using linear mixed modelling, we obtained a change in the ICV of males and females, both in young and middle adulthood that is similar to the one that we obtained with our in-house ICV algorithm (with maybe a small disagreement whether the ICV continues to grow at young adulthood). Possibly, the discrepancy in the first two measures between the results of our in-house ICV extraction algorithm for males and these of FreeSurfer is related to the inherent bias of the FreeSurfer method (see (Klasson et al., 2018), see also the freesurfer manual). Similarly, the discrepancy between the results of the ΔICV_T3,T1_ and 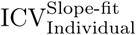 analyses and those of the linear-mixed model for the FreeSurfer analysis is probably related to the nature of the linear-mixed model fitting algorithm. Given the results of the FreeSurfer pipeline eTIV estimations, we tend to believe more to the results of the first two analysis methods of the same FreeSurfer result.

### Relationship to CSF, Gray Matter and White Matter

Having detected age-dependent ICV changes, it might be interesting to ask about the correlation between these age-dependent ICV changes and the trajectories of changes in the brain tissues and the CSF (see Supporting Text). We found moderate to low Pearson correlation-coefficients (P_*Corr*_) between ΔICV_*T*3, *T*1_ and ΔCSF_*T*3, *T*1_ (0.25, see table in Fig. S22), moderate to low P_*Corr*_ between ΔICV_*T*3, *T*1_ and ΔGM_*T*3, *T*1_ (0.31, see table in Fig. S23), and moderate to low P_*Corr*_ between ΔICV_*T*3, *T*1_ and ΔWM_*T*3, *T*1_ (0.25, see table in Fig. S24). Similar results were obtained by linear-fitting of ΔICV_*T*3, *T*1_ to ΔCSF_*T*3, *T*1_, to ΔGM_*T*3, *T*1_ or to ΔWM_*T*3, *T*1_ (see Fig. S22a-d, S23a-d, and S24a-d and tables therein).

However, when we studied the relationship between ΔICV_*T*3, *T*1_ and ΔCSF_*T*3 .*T*1_ or ΔWM_*T*3, *T*1_ as a function of age (see Supporting Text and Fig. S25-26 and S29-30), no meaningful relationships were observed (note that some correlation for males for the CSF case was detected, but only for one of the analysis methods). We concluded that changes in these two measures are unrelated to age-dependent changes in the ICV. By contrast, there might be some very small correlations between age-dependent changes in the ICV and those of the gray matter (see Fig. S27-28 and the supporting text), but it is questionable how significant are they.

### Cross-sectional analysis

Note that the rate of ICV change that we measure in the ΔICV_T3,T1_ and the 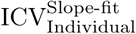 analyses (let alone in the LMM analyses) is not only accelerating but is also smaller than several previous reports that suggested an ICV reduction rate of 0.05-0.29 %/year during most of the young and middle adulthood period (DeCarli et al., 2005; Fillmore et al., 2015). For example, our ΔICV_T3,T1_ analysis shows that only at an average age of 57 the ICV reduction rate reaches a value of 0.1 %/year. To try to understand this discrepancy, we have calculated the cross-sectional ICV age dependency of our data with the same dataset that was used for the longitudinal analysis. The results of this analysis are shown in Fig. 5. We measured a statistically significant constant ICV value decrease with the cross-sectional approach for all time points for both males and females (see table in Fig. 5). Stratifying the cross-sectional data according to the group status in our cohort (control group, sibling group, and people that were diagnosed with schizophrenia), showed that in most of the cases we observed a statistically significant average cross-sectional age-dependent decreased ICV value (at least on the 0.1 significant level). The calculated ICV reduction rates in those cases are similar to the one that is observed in the general cross-sectional analysis (see Fig. S31 of the SI). Exception for this observation, such as for the females control group, where the p-value of the cross-sectional ICV analysis was always above 0.1, can be attributed to a lack of statistical power due to small number of participants.

**Figure 5:**
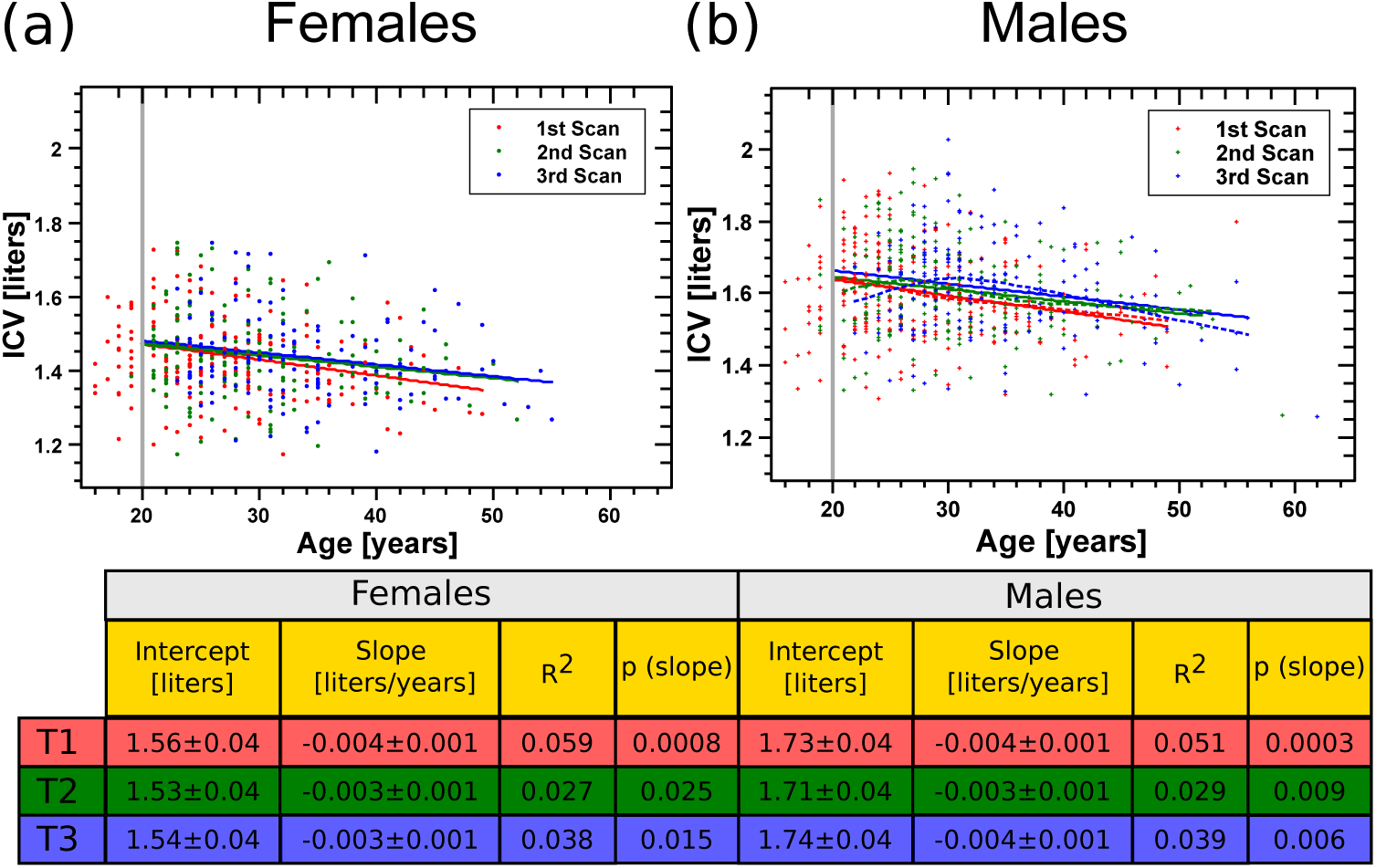
Cross-sectional results for the ICV. Cross-sectional relationship between the ICV and the age of the subject at the time of scan for females (a) and males (b). Each dot represents the ICV value for an individual. Lines are fit of the cross-sectional data to linear functions. Gray vertical lines indicate the minimum age that was used for the fitting. Red - first scan; Green - second scan; Blue - third scan. Dashed lines in (b) are the GAM fits with three degrees of freedoms. For clarity, the GAM fits with two degrees of freedom are not shown. Statistical characteristics of the fit are shown in the table below the graphs for the cases of females and males at the different time points. *R*^2^- goodness of fit. p-value - statistical significance of the slope parameter of the fit. Error values - ±s.e.

The calculated cross-sectional rate of ICV change for females and males based on the average slopes of the fits for the three time-points in Fig. 5 and the predicted average ICV at age 20 based on the same data is 0.21 %/years for males (3.8 ± 0.5 ml/year) and 0.22 %/year for females (3.5 ± 0.6 ml/year). We attribute this rate of ICV change to a generational effect rather than to a real ICV reduction with age.

Analysis of higher-order terms above the linear one in the cross-sectional data (see Material and Methods) is not statistically significant for females (p-values of ANOVA between GAM fits with one and two degrees of freedom = 0.62, 0.6, 0.35 for the first second and third waves of the study. p-values of ANOVA between GAM fits with one and three degrees of freedoms = 0.7, 0.7, 0.19 for the first second and third waves of the study). For males, we did obtain a statistically significant indication for higher-order terms at the second and third waves of the study, see Fig. 5a (p-values of ANOVA between GAM fits with one and two degrees of freedom = 0.078, 0.12, 0.01 for the first second and third waves of the study. p-values of ANOVA between GAM fits with one and three degrees of freedoms = 0.11, 0.035, 0.006 for the first second and third waves of the study). However, only for the third wave of the study, the data suggest an apparent quadratic or cubic behavior. Similarly, a running average analysis with a window of six years did not show deviation from linearity for females (see Fig. S32a of the SI), and only a marginal deviation from linearity for males at the second and third waves of the study (see Fig. S32b of the SI). Thus, the cross-sectional data may show some marginal non-linearity. However, we also attribute that non-linearity to a generational effect.

## Discussion

In this work, we have studied the aging-trajectory of the ICV by applying a longitudinal analysis. Using the data from our cohort, we have shown that the ICV does not stay constant during adulthood. Instead, our analysis suggests that, in the Dutch GROUP cohort, the ICV shows a non-linear aging pattern. To get a feeling for the amount of ICV change between the ages of 20 and 55 years according to different analysis modes that we applied, we present several possible ICV aging trajectories in Fig. 6a,b (assuming the average ICV at age 20). To the best of our knowledge, this study is the first extensive longitudinal ICV aging-analysis during early and middle adulthood.

**Figure 6:**
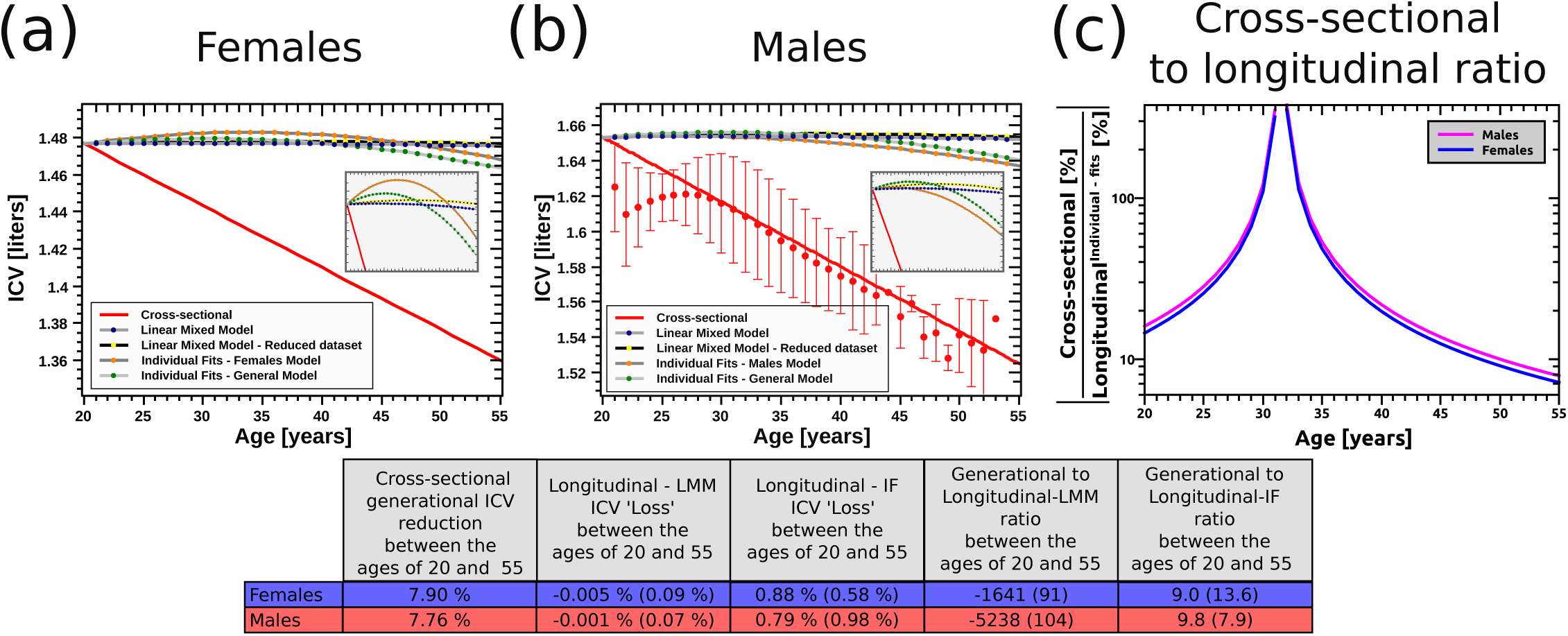
Cross-sectional vs. longitudinal results. Models for the cross-sectional and the longitudinal ICV trajectories for females (a) and males (b). Dark-gray lines with orange dots and light gray lines with green dots - ICV trajectories calculated based on the individual ICV fits for males and females separately or males and females together for the reduced dataset (see table in Fig. 3). Gray lines with yellow dots and dark lines with yellow dots - ICV trajectories calculated based on the linear mixed models for the reduced and the complete datasets receptively (see table in Fig. 4). Smooth red lines - predicted ICV effect calculated based on the average values of the cross-sectional linear fits for males and females separately (see table in Fig. 5). In (b), red dots with error bars - average and SD of the cross-sectional GAM model fits with three degrees of freedom. Insets - zoom-in of the corresponding graphs over the whole x-axis range and over the y-axis range of 1.46-1.4845 liters for females and 1.625-1.659 liters for males (graph background way grayed for emphasis) For clarity, the males cross-sectional GAM model results are emitted from the inset. (c) Absolute value ratio ([%]/[%]) of the cross-sectional generational effect based on the linear fits and the longitudinal ICV loss. For clarity, only the case for the 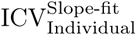 model for females and males together is shown. Note that the graph represent absolute values so that negative ratio (before the age of ≈31) and positive ones after that age are depicted with the same value. The table below the graphs shows, as an example, the predicted ICV loss between the of age 20 and 55 according the linear mixed models for females and males of the reduced dataset (see table in Fig. 4) and the general model for the 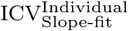 (see table in Fig. 3). In parentheses - the corresponding same values based on the 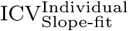 fits for females and males separately or for the LMM model for the complete dataset. The table also shows the predicted generational effect for females and males based on the linear models cross-sectional analysis (see table in Fig. 5). For the longitudinal case, trajectories was calculation by integrating the ICV rate of change equation: 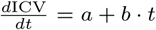, to obtain: 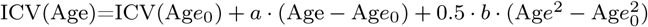. Since there was no substantial differences between the GAM model for males and the linear model, only the results of the linear fit are shown in the table.

All three analysis methods that we have used showed a consistence pattern of ICV changes, which amount to an increase in the ICV during young adulthood that is replaced by an ICV shrinkage later on at middle adulthood (see Fig. 2-4). Yet, using the three different modes of analysis, we detected somewhat different rates of age-dependent ICV changes. For example, the ΔICV_T3,T1_ analysis suggest an average ICV enlargement rate of +0.03 %/year immediately after age 20 years (0.5 ml/year for an averaged male with an ICV of 1.65 liters or 0.4 ml/year for an averaged female with an ICV of 1.48 liters) that evolves to an ICV reduction rate of −0.09 %/year at the age of 55 (1.5 ml/year for an averaged male with an ICV of 1.64 liters or 1.2 ml/year for an averaged female with an ICV of 1.46 liters). Similarly, the 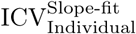 analyses for both males and females suggest an average addition of 0.5 ml/year to the ICV at age 20 years that is replaced by a decrease of 1.3 ml/year at age 55. In principle, these changes are at least one order of magnitude smaller than the ones that were detected for total brain change at this age range (Hulshoff Pol and Kahn, 2008).

Interestingly, the decade of life where we found the ICV to change direction between growth and decline corresponds well to the decade of life where the amount of white matter in the brain stops growing and starts shrinking or starts accelerated shrinking (Good et al., 2001; Kruggel, 2006; Schippling et al., 2017). This raises the possibility that these two processes result from a shared genetic cause. However, the fact that we did not detect age-dependent correlations between the change in the ICV and these of the WM, testify against this hypothesis. Further research is needed to clarify the relationship between the age-dependent trajectories of these two processes. Moreover, the small correlation between the age-dependent changes in the gray matter and those of the ICV calls for further research that will try to find out if aging of the ICV and aging of the gray matter are indeed very slightly correlated. And if they are, are they related one to the other by a common cause, or are they only non-causally correlated.

It is also notable that using the 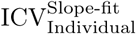 analysis we obtain a total increase of 2.9 ml from the age of 20 to the age of 35. This result is the same order of magnitude as in (Liu et al., 2003). Similarly, if we extrapolate the 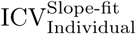 model until the age of 75, we obtain a total ICV reduction of 3.0% for males and 3.4% for females (based on the ICV at age 20). This estimate is the same order of magnitude as in (Royle et al., 2013). Thus, most probably, ICV continues to grow, or stays constant, until sometime during the fourth decade of life. Subsequently, it starts to decline in non-linear manner. We suggest that (Liu et al., 2003) did not observe such a decline in people older than 35 years due to a lack of statistical power. In addition, it is possible that at an older age than 55 the ICV decline depends on higher age powers than age^2^ that we did not detect. Higher order age dependency may be the reason that our predictions are somewhat smaller than in (Royle et al., 2013). It should be noted, however, that using the LMM models we predict a smaller positive change at age 20 and a smaller reduction at age 55 (an order of magnitude for the complete dataset analysis, and an ICV change that is in practice almost equal to zero with the reduced dataset analysis at the same age). These small inconsistencies between our different analysis models call for future longitudinal studies with a larger cohort, a longer duration between repetitive MRI scans, and a broader age range.

What could be the source of the ICV aging trajectory? In principle, we can think about three possible physical mechanisms that are not mutually exclusive as correlates with the ICV changes. The first possible physical mechanism is thickening of the meninges. Indeed, it is known the dura mater becomes thicker at old age (Adeeb et al., 2012). However, it is hard to believe that an ICV change of the size that our model predicts for old age will purely result out of meninges thickening. A possible second source of ICV age-dependent changes is thickening of the skull bone. In principle, if, to what extent, and at which age the skull bone gets thicker is still under dispute. The reason for this dispute is that two mechanisms operate on the skull bone. On the one hand, it is known that aging is accompanied by bone mass loss. On the other hand, changes in the ICV must be compensated by parallel changes in other tissues. Interestingly, thickening of the skull was detected in several cases (May et al., 2012; Royle et al., 2013). Thus, it is plausible that skull bone thickening is responsible for all, part of, or correlates with, the ICV changes. Finally, shape-changes transformations of the skull as a function of age (e.g., in the cephalic index) may also result in ICV changes (Albert et al., 2007; Urban et al., 2014). The elucidation of the causal mechanisms for the longitudinal ICV changes that we measure is outside the scope of this article and awaits future work.

When comparing the longitudinal analysis to the cross-sectional one (see Fig. 6a,b,c), the cross-sectional analysis suggests a much larger ICV difference in comparison to the longitudinal analysis between the ages of 20 and 55. What can be the source of this discrepancy? The most obvious answer is a population level of secular growth throughout the life of the cohort participants. Indeed, evidence for environmental-dependent secular growth in the last century was documented all around the globe (Cole, 2003; Simsek et al., 2005; Arcaleni, 2006; Marques-Vidal et al., 2008). In particular, the Dutch population has gone through a period of bodily enlargement since the middle of the 19th century probably due to an improvement of nutrient and health conditions during childhood (Fredriks et al., 2000). A large cross-sectional comparison has shown that, on average, between 1950 and 1997 Dutch adults’ height increased by roughly 8 cm. The rate of height increased was, however, subsidizing over these decades so that between 1980 and 1997 the rate was only 1.3 cm/decade. Moreover, recently, this trend has stopped (Schönbeck et al., 2013). The first wave of the GROUP cohort was acquired between 2004 and 2008 and participants in that wave were born between 1951 and 1991. To assess similar stature effect in our data, we note that the average height of the GROUP cohort participants correlates negatively with age (See Fig. S33 of the SI). The average rate of change in height that was measured in the GROUP cohort was 1 ± 0.5 cm/decade. Thus, it is clear that a generational effect subsists in our data. The fact that we, like others before us, attribute the cross-sectional result to a generational effect, can also explain why in different cross-sectional based studies different rates of ICV change were found, sometimes reaching the statistically significant limit and sometimes not. Most probably, these different cross-sectional rates are the results of different secular growth rates.

These results also call for caution when comparing different groups in MRI research. For the Dutch case, even if the control and study group age characteristics deviate one from another by three years, there can be an average difference of ≈ 0.6 % in their ICVs. This difference may bias the conclusions of such studies. One way, to get over this age matching problem is to normalize all brain measures to the ICV. However, this procedure assumes a one-to-one correspondence between ICV to other brain biomarkers extent. Thus, it is important to keep strict age matching characteristics in case-control comparative cross-sectional studies.

It is interesting to note that the rate of cross-sectional ICV change that we detected (≈ 0.2 %/year) is larger than the corresponding cross-sectional secular height growth rate in the same cohort (0.05 − 0.06 %/year based on the height at age 20, see Fig. 15 of the SI). Indeed, additional to secular growth in height, there is also evidence for cranial changes over the last decades in different parts of the world (Buretic-Tomljanovic et al., 2006; Little et al., 2006; Cymek et al., 2015; Kim et al., 2018). Since there need not exist a necessary one-to-one correspondence between height and cranial dimensions, it is possible that the relative secular ICV growth was more significant than that of the height. The fact that 4 out of 7 genetic loci for ICV are known genetic height loci, but that in some cases these loci act discordantly (Adams et al., 2016), stands in accord with this suggestion. In this context and in the light of the correlation between the ICV and the IQ, it is interesting to speculate about the relationships between secular growth rates and the Flynn effect. The Flynn effect is the worldwide increase in phenotypic IQ in the last decades (Flynn, 1987). It is usually assumed that the source of the Flynn effect is purely sociological, i.e., results from better education. However, the possibility that part of this secular rise in IQ is also physical by origin was also suggested (Mingroni, 2004). Thus, it is interesting to ask whether the more significant rate of ICV growth relative to the body (height) liberate more brain material for cognitive abilities that are measured by the IQ scale, which results in the secular rise in the phenotypic IQ. Though outside the scope of this work, we believe that such hypothesis merits additional future inspection.

This study has several limitations that need to be addressed when interpreting its finding. First, for the two individual analysis modes that we used (ΔICV_T3,T1_ and 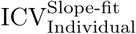), we obtained statistically significant ICV changes only after applying a filtering procedure over the data. Similarly, for the global LMM analysis, the age-related ICV changes that were calculated using the reduced dataset are somewhat smaller than those that were obtained from the complete dataset. One may question our filtering procedure.

We believe that we are not imposing arbitrary filtering criteria to obtain a statistically significant result. On the contrary, using the filtering mechanism, we are exposing a genuine relationship between ICV and age that was hidden under the noise. Two reasons lead us to believe that this is indeed the case. First, we have filtered out results with the rather large effect (more substantial ICV difference). In general, one would expect that keeping cases with a rather large ICV change will enhance the false positive probability and not the other way around. Second, a post hoc additional manual inspection of several MRI scans that were filtered out by our criteria show that, in many cases, small cranium or meninges segments were included in the ICV mask. These segments were the reason that the data for these individuals showed substantial ICV changes. However, these segments were not detected during the original manual inspection of the ICV masks. It is easy to understand why this is the case. The most considerable ICV difference that we measured was around 100 ml (see Fig. 4b). It can be expected that small non-ICV segments sometimes cannot be detected by a human inspection in a large cohort when the researcher inspects only a limited number of slices and the non-ICV tissue is distributed over a large part of the ICV surface. A second limitation of this study is related to the field strength of the scanners that was used for the T1w acquisition. We have used MRI scans that were acquired on a 1.5 Tesla scanners. However, modern scanners usually operate at 3 Tesla and sometimes at higher field strengths. The reason that the data was acquired with a rather low field strength is tightly related to its longitudinal acquisition protocol which started with the recruitment of first individuals at 2004, when not many 3 Tesla scanners were operational, and ended at 2014. Since we are interested only in a global measure of the total ICV and not in detailed regional gray or white matter trajectories, we believe that this limitation is less crucial. MRI studies that used similar field strength were commonly used to study brain aging before the advance for higher field strength scanners, and many of their findings were corroborated by later research (see, e.g., (Good et al., 2001; Buckner et al., 2004)). Yet, future studies will probably use higher field strength when corroborating our findings.

A third limitation of our study is related to the fact that the acquisition was performed on two scanners and not one. It is known that different scanners introduce a systemic error and that can lead to a bias in our results (Jovicich et al., 2009). Indeed, when applying the linear mixed model analysis to the reduced dataset, we found that incorporating the scanner identifier as a confounder into the model made the linear mixed model statistically significant better. Yet, in that case, we also found changes in ICV in young and middle adulthood. Thus, we believe that even if the acquisition protocol that included two scanners introduced some bias to our results. It did not change their nature. Still, despite an extended longitudinal acquisition period, future studies will be better using only a single scanner to minimize bias.

A fourth limitation of our study is related to the fact that we do not have information regarding childhood conditions. Aging and development are related, and each one of these processes is also related to environmental conditions. For example, it is known that the puberty timing is genetically related to risk for prostate and breast cancer at old age (Day et al., 2017). Similarly, the age of menarche is associated with mortality from cardiovascular disease at old age (Lakshman et al., 2009). In addition, the age of menarche changed over the last 50 years in the Netherlands (Talma et al., 2013). Thus, it is possible that childhood conditions that influenced development and growth may also influence ICV changes in adulthood. It should also be noted that our cohort is primarily a Dutch one, and specific conditions of the Dutch population (genetic or environmental) could have influenced the rate and nature of the ICV aging trajectory. Thus, future longitudinal studies will be better to incorporate childhood data as an additional confounder. Moreover, it is crucial to study ICV aging in populations that are not from European and European-North American origin to obtain a better representation of ICV and brain aging processes over a wide range of human conditions (Okonkwo et al., 2012; Falk et al., 2013).

A fifth limitation of our study is related to head motion during scanning. It is known that head motion can hinder brain imaging measures (Reuter et al., 2015). These head motions were related to genetic factors that characterize each individual (Zeng et al., 2014) but also become more apparent for older people (especially above the age of 50, see (Savalia et al., 2017)). In principle, head motions can act as a confounder for the estimate of age effects since what is estimated as a longitudinal age effect may be due to more substantial head motion during the scan. There are a couple of ways to address this problem in the future, including collecting multiple T1w scans for the same subject at each wave of the study or utilizing orthogonal scan protocols such as fMRI (Reuter et al., 2015; Savalia et al., 2017). Yet, we believe that there is a high chance that our result represents a real longitudinal ICV effect for two reasons. First, previous studies suggested that increase head motion is more pronounced at old age while our study sample is composed of young and middle adults. Second, brain measures such as the ICV, when they are obtained by global registration of the scan with high contrast should be less sensitive to the ‘smearing’ effect of the gray levels in the scans due to head motion.

The sixth limitation of this study might be related to the type of cohort that was used that includes people that were diagnosed with schizophrenia, their relative, and a control group. Including the first group of these three in the study may place an unwarranted confounding factor. Indeed, as stated in the introduction, it is known that people who suffer from psychosis have, on average, smaller ICV (Mean Weighted Cohen’s d of −0.14 see (Haijma et al., 2012)). Nevertheless, this fact does not necessarily mean that, besides the small overall ICV difference, also the trajectory of ICV change in young and middle adulthood is different among people that were diagnosed with schizophrenia and those that were not. Indeed, we were not able to detect a difference in the behavior of the ICV changes between these three groups. Incorporation of the group status to the linear mixed model analysis resulted in a high p-value. Similarly, the exclusion of this group of people from our analysis did not change the results. One can still claim that the incorporation of relatives of people that were diagnosed with schizophrenia might have biased our result. We did not carry an analysis the control group alone, mainly due to statistical power issues. However, it is hard to conceive that the relative group will substantially bias the results as these people are not recognized as possessing a sickness in any medical sense. Still, more subtle issues regarding the relative group might operate here (see the discussion in the next paragraph). Hence, future corroboration of our results in other cohorts is advisable.

Finally, a seventh limitation of the study is that we cannot tell for sure if our finding represents normal aging or underlying aging processes that are related to the lifestyle or genetics of the participants. Today, it is known that several brain pathological processes start well before any clinical signed are observed (e.g., for Alzheimer disease see (Filippini et al., 2009; Berti et al., 2011; Okonkwo et al., 2012; Dowell et al., 2016)). Thus, it can be the case that the longitudinal changes that we detected are related to some environmental factors, genetic factors, or their interactions. Yet, the fact that the group status (patient, sibling, control) did not come up significantly in our linear mixed model suggests that, as long as normal aging is defined as the average degenerative processes in some general population due to solely to shared conditions (environmental or genetic) in that population, our study at least indicate such behavior in the Dutch population.

To sum up the main result of this work, in this manuscript we present an extensive longitudinal analysis of ICV aging-trajectory that shows a non-linear behavior. We hope that this work will inspire future longitudinal analysis of the ICV and other brain biomarkers aging-trajectories, especially using big data cohorts. We believe that conducting longitudinal-designed studies over multiple conditions and in multiple populations is the most trust-worthy method to obtain accurate measures of brain aging and its dependency on various genetic and environmental factors.

## Acknowledgments

The authors would like to thank Hakim Achterberg, Marcel Koek, Adriaan Versteeg, Thomas Phil, Thomas Kroes, Baldur van Lew, Marcel Zwiers and Seyed Mostafa Kia for a collaboration within the BBMRI-NL work-package 3.

This work was supported by the Netherlands Organization for Scientific Research (NWO 184.033.111), Biobanking and BioMolecular resources Research Infrastructure The Netherlands (BBMRI-NL2.0), and by the ENIGMA World Aging Center grant (NIH 1R56AG058854-01, subaward 112068003).

The infrastructure for the GROUP study is funded through the Geestkracht programme of the Dutch Health Research Council (Zon-Mw, grant number 10-000-1001), and matching funds from participating pharmaceutical companies (Lundbeck, AstraZeneca, Eli Lilly, Janssen Cilag) and universities and mental health care organizations (Amsterdam: Academic Psychiatric Centre of the Academic Medical Center and the mental health institutions: GGZ Ingeest, Arkin, Dijk en Duin, GGZ Rivierduinen, Erasmus Medical Centre, GGZ Noord Holland Noord. Groningen: University Medical Center Groningen and the mental health institutions: Lentis, GGZ Friesland, GGZ Drenthe, Dimence, Mediant, GGNet Warnsveld, Yulius Dordrecht and Parnassia psycho-medical center The Hague. Maastricht: Maastricht University Medical Centre and the mental health institutions: GGzE, GGZ Breburg, GGZ Oost-Brabant, Vincent van Gogh voor Geestelijke Gezondheid, Mondriaan, Virenze riagg, Zuyderland GGZ, MET ggz, Universitair Centrum Sint-Jozef Kortenberg, CAPRI University of Antwerp, PC Ziekeren Sint-Truiden, PZ Sancta Maria Sint-Truiden, GGZ Overpelt, OPZ Rekem. Utrecht: University Medical Center Utrecht and the mental health institutions Altrecht, GGZ Centraal and Delta.

## Authors contribution

YC and HHP conceived the study and wrote the article. RB and HS developed the ICV extraction algorithm. YC implemented the ICV algorithm, preformed the experiment, and analyzed the data. MVDN helped in the initial stages of the data analysis. WC and RK contributed data. AVDL and WN contributed expertise. All authors critically reviewed the manuscript drafting.

