## Supporting Information for "Changes in the intracranial volume from early adulthood to the sixth decade of life: A longitudinal study"

### Supporting Information for the article - Changes in the intracranial volume from young adulthood to the sixth decade of life: A longitudinal study

Yaron Caspi[a], Rachel Brouwer[a], Hugo Schnack[a], Marieke E. van de Nieuwenhuijzen[a], Wiepke Cahn[a], René S. Kahn[a,b], Wiro J. Niessen[c], Aad van der Lugt[c], Hilleke Hulshoff Pol[a]

[a]*Brain Center Rudolf Magnus, University Medical Center Utrecht, The Netherlands*
[b]*Department of Psychiatry, Icahn School of Medicine at Mount Sinai, New York, NY, USA*
[c]*Department of Radiology, Erasmus MC: University Medical Center Rotterdam, The Netherlands*

#### Supporting Text

*ICV differences between adjacent scanning waves*

To explore the longitudinal ICV changes between adjacent scanning waves, we studied ICV differences between the second and first scanning waves ($\Delta\mathrm{ICV}_{\mathrm{T2,T1}} \equiv \frac{2\cdot[\mathrm{ICV}_{\mathrm{T2}} - \mathrm{ICV}_{\mathrm{T1}}]}{[\mathrm{ICV}_{\mathrm{T2}} + \mathrm{ICV}_{\mathrm{T1}}]\cdot[\mathrm{Age}_{\mathrm{T2}} - \mathrm{Age}_{\mathrm{T1}}]}$), and the corresponding differences between the third and second scanning waves ($\Delta\mathrm{ICV}_{\mathrm{T3,T2}}$). For $\Delta\mathrm{ICV}_{\mathrm{T2,T1}}$ we indeed obtained a similar behavior to the one we observed for $\Delta\mathrm{ICV}_{\mathrm{T3,T1}}$. Namely, we did not obtain a statistically significant age dependence behavior of the ICV change for the complete dataset in our cohort (see Fig. SS10a of the SI and the corresponding table). However, after filtering out individuals with a high absolute value of measured ICV change (see Fig. SS10c,d of the SI), we obtained statistically significant age-dependent $\Delta\mathrm{ICV}_{\mathrm{T2,T1}}$ for males but not for females (see Fig. SS10b of the SI and the corresponding table). Combining the females and males groups resulted in a statistically significant age-dependent $\Delta\mathrm{ICV}_{\mathrm{T2,T1}}$ fit. The values of the slope, in that case, was consistent with the value that was calculated for $\Delta\mathrm{ICV}_{\mathrm{T3,T1}}$. By contrast, we did not obtain statistically significant $\Delta\mathrm{ICV}_{\mathrm{T3,T2}}$ even after applying the filtering procedure and grouping females and males together (data not shown). Note, however, that the longitudinal changes that we detected are minimal, and the age dependency explains only a small portion of the total variance of the data. Hence, one explanation for the fact that we did not obtain statistically significant $\Delta\mathrm{ICV}_{\mathrm{T3,T2}}$ as a function of age may be related to a lower statistical power as results of a smaller number of subjects in the $\Delta\mathrm{ICV}_{\mathrm{T3,T2}}$ dataset relative to the $\Delta\mathrm{ICV}_{\mathrm{T2,T1}}$ one in the face of a low $R^2$ of the age dependency (see Table 1 of the main article). An alternative explanation is that our data contains too much noise to detect age dependency $\Delta$ICV in that case. Several control analyses that we have performed points to the second explanation as the correct one (data not shown).

*ICV and IQ*

We assessed the known relation between the ICV and the IQ [1, 2]. In light of the predicted negative relationship between IQ and the risk for Schizophrenia [3, 4], we have studied the IQ vs. ICV relationship for the three groups in our cohort separately. For this analysis, we have calculated the average IQ from the three waves of the study and plotted it as a function of the ICV for each wave of the study separately.

For the females control group, we detect statistically significant IQ vs. ICV relationship for the first two waves of the study. For the third wave of the study, the statistical significance was somewhat below the 0.05 limit, probably due to the small size of this group (see Fig. SS2a of the SI). Similar results were obtained for the groups of female siblings and females that were diagnosed with schizophrenia. However, in these cases, the IQ vs. ICV relationship was statistically significant for all the three consecutive measurements (see Fig. SS2b,c of the SI). Similarly, for the males control group, the IQ vs. ICV relationship was statistically significant for the first two waves of the study, but reached only the 0.1 p-value limit for the third wave of the study (see Fig. SS2d of the SI). However, by contrast with the females case, the males' sibling group showed a statistically significant IQ vs. ICV relationship only for the first wave of the study. No correlation between IQ and ICV was found for the other two waves of the study as well as for the males that were diagnosed with schizophrenia (See Fig. SS2e,f of the SI). However, the fact that we obtained the expected linear correlation between the IQ and the ICV for the control group, as well as for the other female groups in our cohort strengthen our conclusion regarding the longitudinal trajectories of the ICV with age as was in the main text. Cases where no correlation between IQ and ICV was found, such as for the groups of males siblings and males that were diagnosed with schizophrenia, probably represent factors that are related to the diagnosis state rather than to a malfunction of our ICV extraction procedure.

*Changes in the ICV and changes in the gray matter, white matter, or the CSF*

To check the relationships between the changes in the ICV and changes in the gray matter (GW), white matter (WM), and the CSF, we first calculated the Pearson correlation coefficients between the changes in the $\Delta ICV_{T3,T1}$ and these of the other measures. As a complementary analysis, we also fitted the relationship between $\Delta ICV_{T3,T1}$ and the other measures to a straight line. As can be seen in figures S22a, S3a, S4a, and the tables therein, we obtained high correlations between $\Delta ICV_{T3,T1}$ and all other measures. Despite the fact that these correlations were significant and the fits looked unbiased (Fig. S22b, S3b, S24b), these correlations might be skewed by the existence of cases with relatively high values of $\Delta ICV_{T3,T1}$ (which we suspected as unreal along with the rest of the article). This is especially evident for the CSF and the WM (see Fig. S22a and S24a), where it looks as if the plot is composed out of a central blob and and two disperse point from both of its sides. Hence, we filtered the results based on $\Delta ICV_{T3,T1}$, similar to the procedure that is described in the main text (see Fig. S9). Indeed, the result of the reduced datasets shows a much lower correlation (moderate to low correlations - see Fig. 22c, 23c, 24c, and tables therein) of the order of 0.25-0.3. These correlations were similar for males and for females, and did not changes by much when the people that were diagnosed with schizophrenia were emitted (for CSF for both males and females $P_{Corr} = 0.32$, 0.95 CI=0.17-0.46, p=4.2e-5, GM for both males and females $P_{corr} = 0.3$, 0.95 CI=0.15-0.44, p=0.0002, for WM for both males and females $P_{corr} = 0.21$, 0.95 CI=0.06-0.36, p=0.008). Note that the fit for the CSF looks good (Fig. 22d), less good for the GM (Fig. S23d), and not so good for the WM (Fig. S24d)

However, the fact that we detected significant correlations between $\Delta ICV_{T3,T1}$ and changes in the GM, WM, and CSF does not testify that the age-dependent trajectories that we detected for $\Delta ICV_{T3,T1}$ are also manifested in correlated age-dependent changes in the three other measures. The reason for this putting forward this query is related to the fact that in general, there was a large spread both for the correlation coefficients and for the age-related $\Delta \mathrm{ICV}_{T3,T1}$ trajectories (in other words, in both cases, the calculated linear fits explain only a small portion of the variations in the data). Thus, it can be the case that there are significant correlations between $\Delta \mathrm{ICV}_{T3,T1}$ and changes in the other three measures, but these correlations are independent form the aging processes.

To check this possibility we applied two methods. First, we analyzed the relationship between $\Delta \mathrm{CSF}_{T3,T1}/\Delta \mathrm{ICV}_{T3,T1}$ as a function of age (and similar for the GM and the WM). Second, we fitted the relationship between $CSF_{T3,T1}$ and $\mathrm{ICV}_{T3,T1}$ to a straight line for each individual and analyzed the changes in the slope of these fits as a function of age (and similar for the GM and the WM). The results of the analysis of the changes between the first and the last epochs of the study are shown in Fig. S5a, S7a, and S9a for the CSF, GM, and WM, respectively (and the tables therein). The results of the analysis individual slopes of the fits are shown in Fig. S6a, S8a, and S30a for the CSF, GM, and WM, respectively (and the tables therein). As can be seen, no significant relationships were observed for most of these cases. Only for the GM for females and only in one of the analysis methods the fit reached the 0.05 significant level (but was above the Bonferroni corrected values 0.05/3=0.017). Similarly, for the WM, only in one of the analysis methods, we observed p-values below 0.017 (for females and both females and males together - see Fig. S29a).

However, these lack of significant relationships (or their existence) might be skewed by cases with large absolute values. Thus, we used the same approach that we took in the rest of this manuscript and filtered out results with large absolute vales for $\Delta \mathrm{CSF}_{T3,T1}/\Delta \mathrm{ICV}_{T3,T1}$, see Fig. 25b (and similarly for the GM and WM, see Fig. S27b, S29b). Similarly, we filtered out cases with large absolute values for $\{\mathrm{CSFtoICV}^{\text{Slope-fit}}_{\text{Individual i}}\}$, see Fig. 26b (and similarly for the GM and WM, see Fig. S28b, S30b).

The results of the linear fits for the reduced datasets are shown in Fig. S25c for $\Delta \mathrm{CSF}_{T3,T1}/\Delta \mathrm{ICV}_{T3,T1}$, S27c for $\Delta \mathrm{GM}_{T3,T1}/\Delta \mathrm{ICV}_{T3,T1}$, and S29c for $\Delta \mathrm{WM}_{T3,T1}/\Delta \mathrm{ICV}_{T3,T1}$ (and the tables therein). The results of the linear fits for the reduced datasets are shown in Fig. S26c for $\{\mathrm{CSFtoICV}^{\text{Slope-fit}}_{\text{Individual i}}\}$, S28c for $\{\mathrm{GMtoICV}^{\text{Slope-fit}}_{\text{Individual i}}\}$, and S30c for $\{\mathrm{WMtoICV}^{\text{Slope-fit}}_{\text{Individual i}}\}$. For the WM, the reduced dataset showed no significant age-dependent relationships for both data analysis methods. Removing the people that were diagnosed with schizophrenia from the analysis did not change these results (data not shown). For the CSF, similar results were obtained.

For the GM, the age-dependent trajectories show some small linear relation for males and both females and males using the $\Delta \mathrm{GM}_{T3,T1}/\Delta \mathrm{ICV}_{T3,T1}$analysis methods. Similar results, but less significant,were obtained for females only. For the $\{\mathrm{GMtoICV}^{\text{Slope-fit}}_{\text{Individual i}}\}$ analysis method, the linear fits showed smaller p-values compared to the other analysis method, but these fits probably show the same direction of the effect (especially for females and females and males together). Emitting the people that were diagnosed with schizophrenia from the analysis for both males and females together did not change the results of the analysis for both data analysis methods significantly (data not shown). Thus we can conclude that our data might suggest some very small correlation between the age-dependent trajectory of the ICV and this of the GM. However, the fact that the linear fits were not good for both analysis methods (see Fig. 27d and 28d) and the fact that the correlation between the age-dependent trajectory of the ICV and this of the GM were rather minute call for caution with the interpretation of this result. Further research is warranted to analyze the relationship between the aging of the ICV and the aging of the GM.

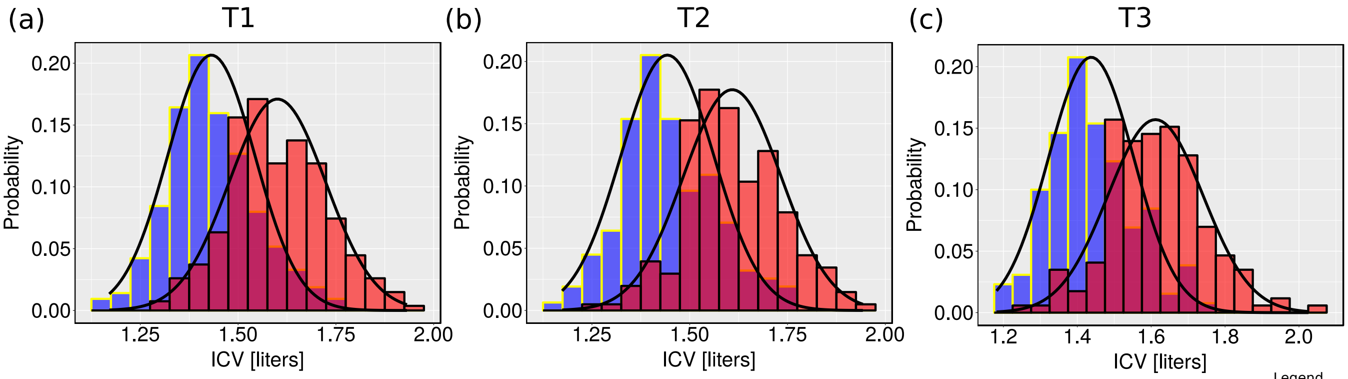


| | ICV T1 [Liters] | ICV T2 [Liters] | ICV T3 [Liters] |
|---|---|---|---|
| Females | 1.4±0.11 | 1.4±0.12 | 1.4±0.12 |
| Males | 1.6±0.12 | 1.6±0.13 | 1.6±0.13 |

Figure S1: **ICV Distributions.** Bar graph distributions for the measured ICV values during the first wave (a), the second wave (b) and the third wave of the study (c). Red - Males, blue - Females. Solid lines - plots of normal distributions with the same mean and s.d. as were calculated for the data. Average and s.d. parameters for the three scanning periods are shown in the table below the graphs.

### Females

(a) Control Group (b) Siblings Group (c) Patients Group

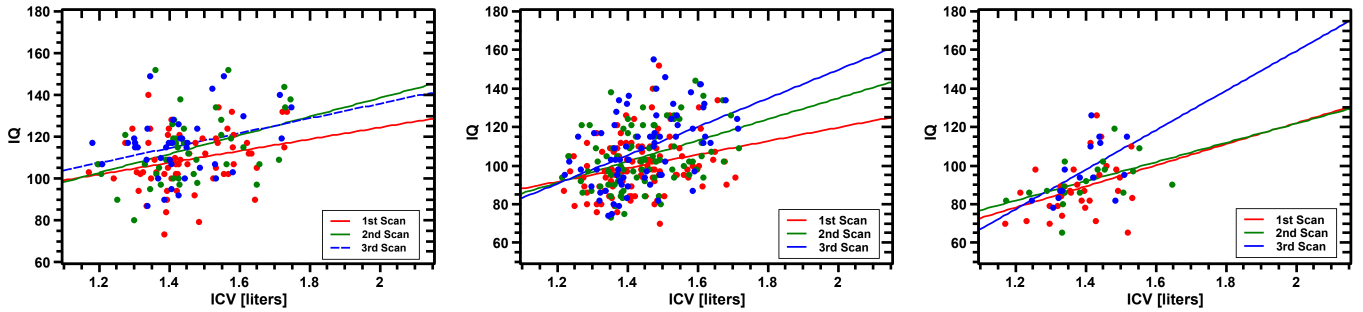


| | Control Group | | | | Siblings Group | | | | Patients Group | | | |
|---|---|---|---|---|---|---|---|---|---|---|---|---|
| | Intercept | Slope [Liters$^{-1}$] | $R^2$ | p (slope) | Intercept | Slope [Liters$^{-1}$] | $R^2$ | p (slope) | Intercept | Slope [Liters$^{-1}$] | $R^2$ | p (slope) |
| T1 | 63±19 | 28±13 | 0.053 | 0.037 | 49±18 | 35±12 | 0.058 | 0.006 | 12±35 | 55±25 | 0.106 | 0.039 |
| T2 | 49±24 | 45±16 | 0.124 | 0.009 | 25±21 | 55±14 | 0.133 | 0.0003 | 22±30 | 50±22 | 0.195 | 0.033 |
| T3 | 75±26 | 36±18 | 0.069 | 0.057 | 2±26 | 74±18 | 0.192 | 7.8e-5 | -46±61 | 103±44 | 0.25 | 0.035 |

### Males

(d) Control Group (e) Siblings Group (f) Patients Group

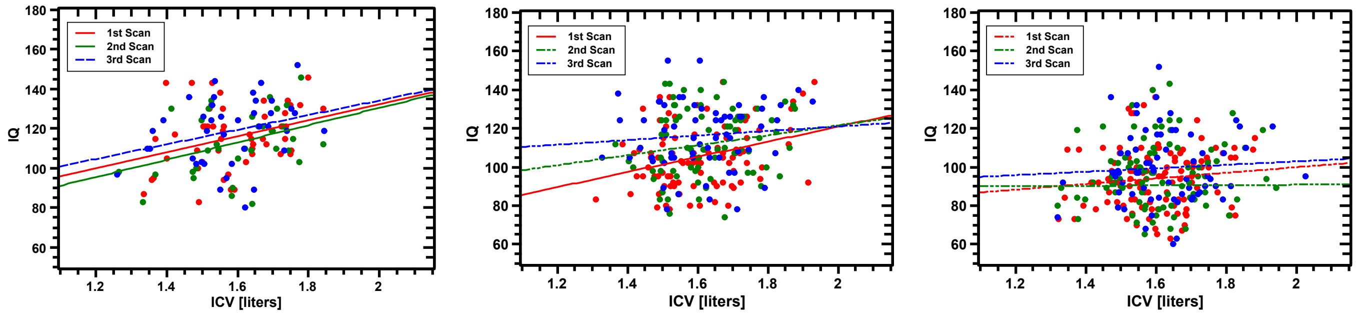


| | Control Group | | | | Siblings Group | | | | Patients Group | | | |
|---|---|---|---|---|---|---|---|---|---|---|---|---|
| | Intercept | Slope [Liters$^{-1}$] | $R^2$ | p (slope) | Intercept | Slope [Liters$^{-1}$] | $R^2$ | p (slope) | Intercept | Slope [Liters$^{-1}$] | $R^2$ | p (slope) |
| T1 | 52±27 | 40±17 | 0.089 | 0.019 | 43±20 | 39±12 | 0.088 | 0.002 | 71±20 | 14±12 | 0.004 | 0.236 |
| T2 | 43±31 | 44±19 | 0.111 | 0.029 | 70±27 | 26±16 | 0.019 | 0.122 | 76±22 | 13±14 | -0.002 | 0.365 |
| T3 | 60±32 | 37±20 | 0.060 | 0.072 | 97±28 | 12±17 | -0.009 | 0.494 | 85±29 | 22±31 | -0.011 | 0.622 |

Figure S2: **IQ versus ICV**. IQ vs. ICV for the different groups in our study for (a-c) females and (d-e) males. Each dot represents the data for one individual. (a,d) control group; (b,e) group of siblings of subjects that were diagnosed with schizophrenia; (c,f) group of subjects that were diagnosed with schizophrenia. Red - first scan, Green - second scan, Blue - third scan. In all panels, lines are fits of the different datasets to linear functions. Full lines - fits with a slope that is statistically significant at the $p < 0.05$ level. Striped lines - fits with a slope that is statistically significant at the $p < 0.1$ level. Striped-dotted lines - fits with a slope that is not statistically significant. The table below the graphs shows the fitting values for the cases of females or males. $R^2$- goodness of fit. p-value - statistical significance of the slope parameter of the fit. Error values - ±s.e.

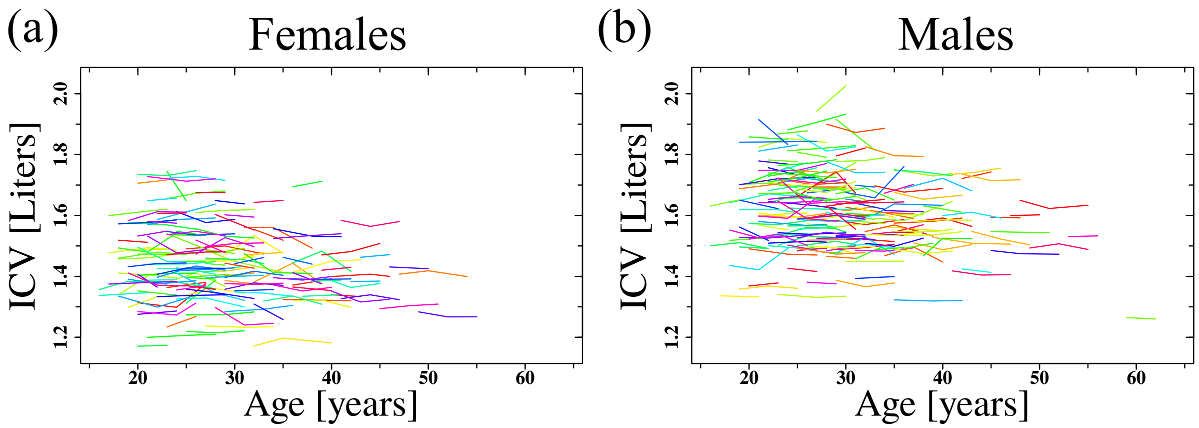


Figure S3: **ICV Longitudinal trajectories.** Longitudinal trajectories of the ICV for females (a) and males (b). Only cases with two or three consecutive measurements are shown. The ICV trajectories of individuals are drawn in separate colors.

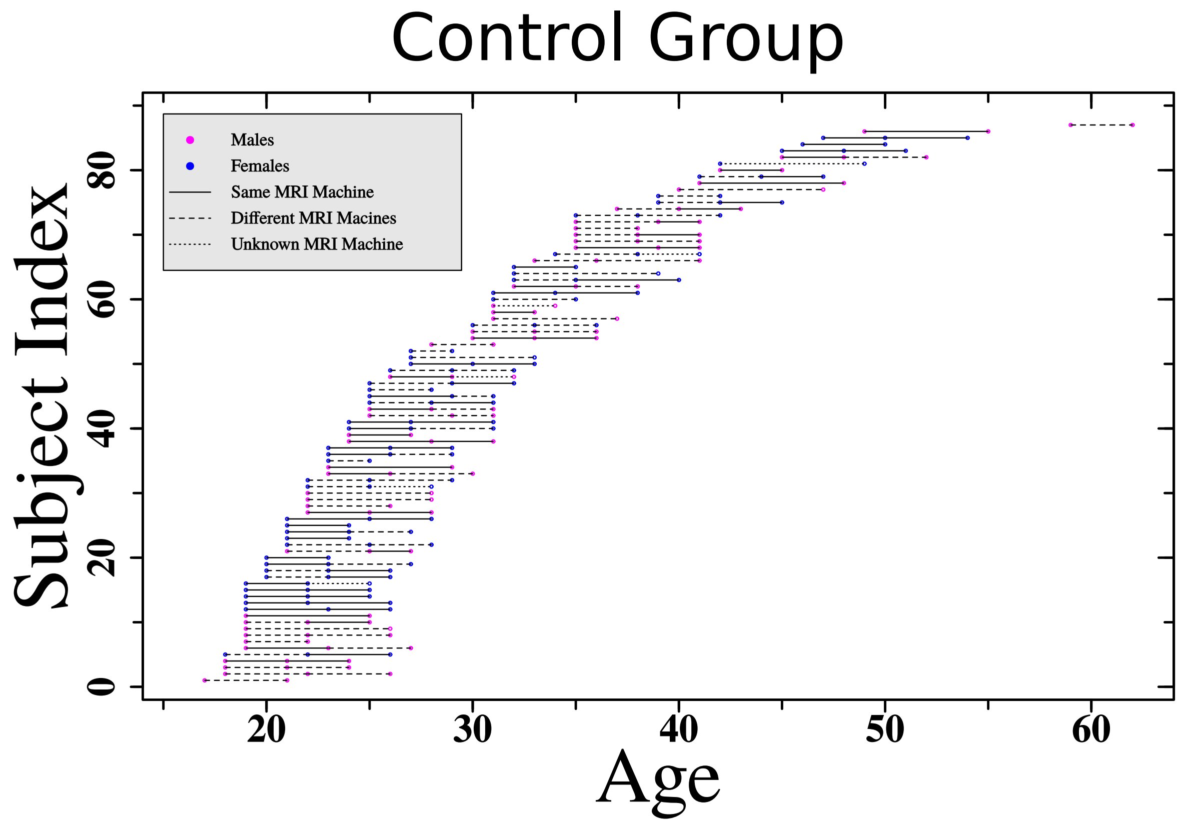


Figure S4: **Spaghetti plot for the control group.** A Spaghetti plot of the control group relative to age, sex and scanner identifier. Males are depicted in pink. Females are depicted in blue. Full lines represent a case where two consecutive scans were carried in the same scanner. Dashed lines represent two consecutive scans that were carried on two different scanners. Dotted lines represents two consecutive scans where the scanner identifier of one of the two scans is unknown. This plot is an adjacent plot to plots S5 and S6. The Spaghetti plots are presented according to the group status of the participants only to make the plots clear relative to the considerable number of participants (that can hardly fit one graph).

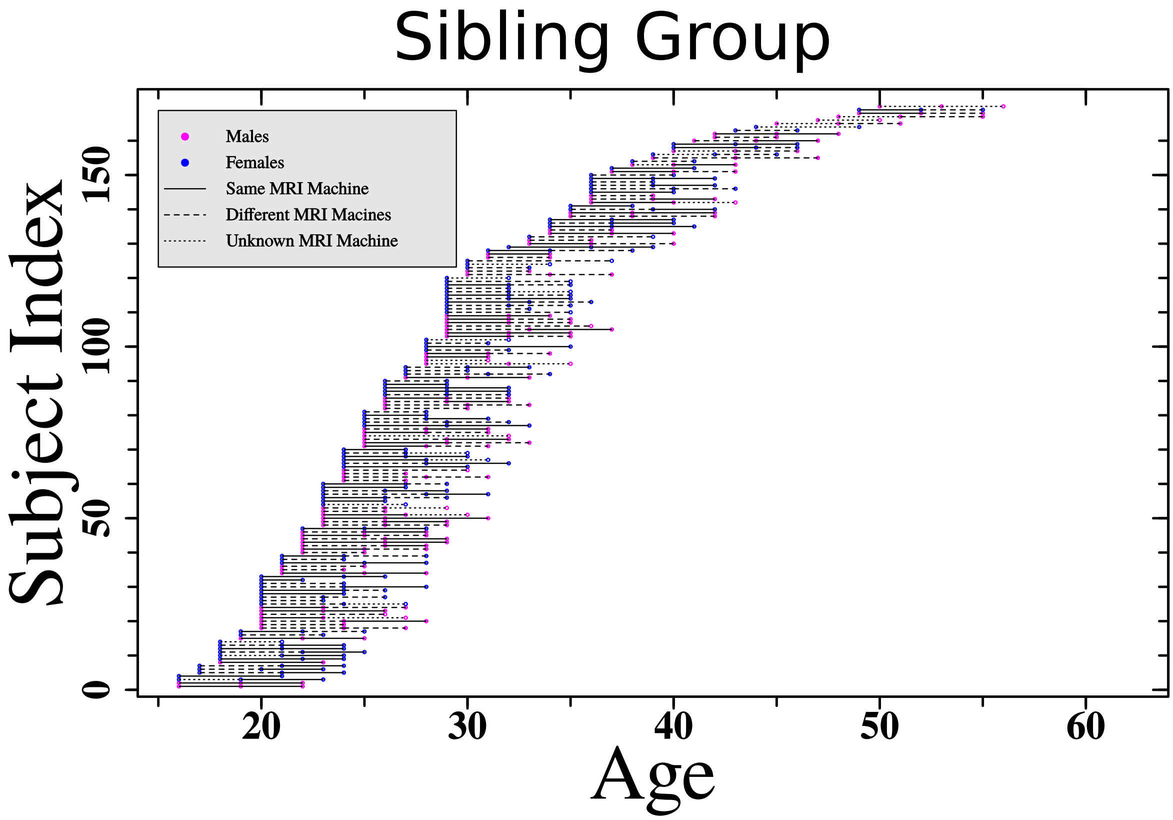


Figure S5: **Spaghetti plot for the sibling group.** A Spaghetti plot of the sibling group relative to age, sex and scanner identifier. Males are depicted in pink. Females are depicted in blue. Full lines represent a case where two consecutive scans were carried in the same scanner. Dashed lines represent two consecutive scans that were carried on two different scanners. Dotted lines represents two consecutive scans where the scanner identifier of one of the two scans is unknown. This plot is an adjacent plot to plots S4 and S6. The Spaghetti plots are presented according to the group status of the participants only to make the plots clear relative to the considerable number of participants (that can hardly fit one graph).

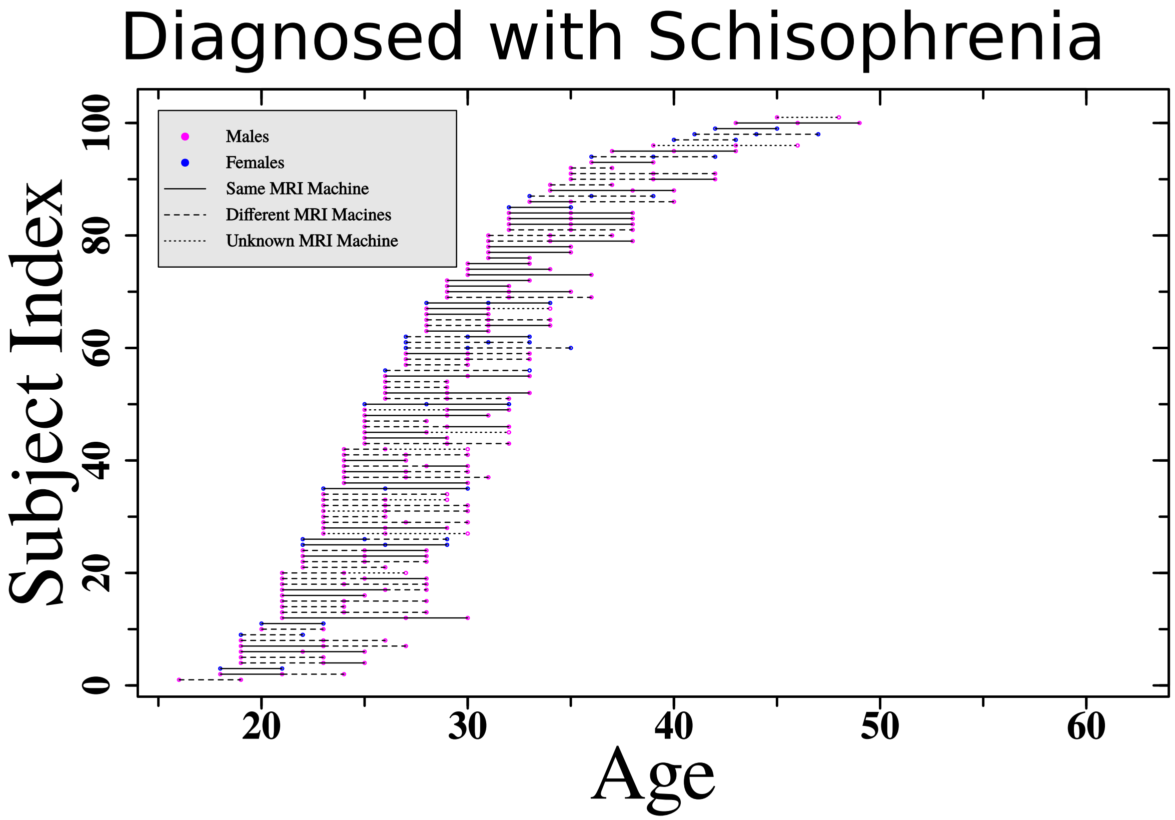


Figure S6: **Spaghetti plot for the control group.** A Spaghetti plot of the control group relative to age, sex and scanner identifier. Males are depicted in pink. Females are depicted in blue. Full lines represent a case where two consecutive scans were carried in the same scanner. Dashed lines represent two consecutive scans that were carried on two different scanners. Dotted lines represents two consecutive scans where the scanner identifier of one of the two scans is unknown. This plot is an adjacent plot to plots S4 and S5. The Spaghetti plots are presented according to the group status of the participants only to make the plots clear relative to the considerable number of participants (that can hardly fit one graph).

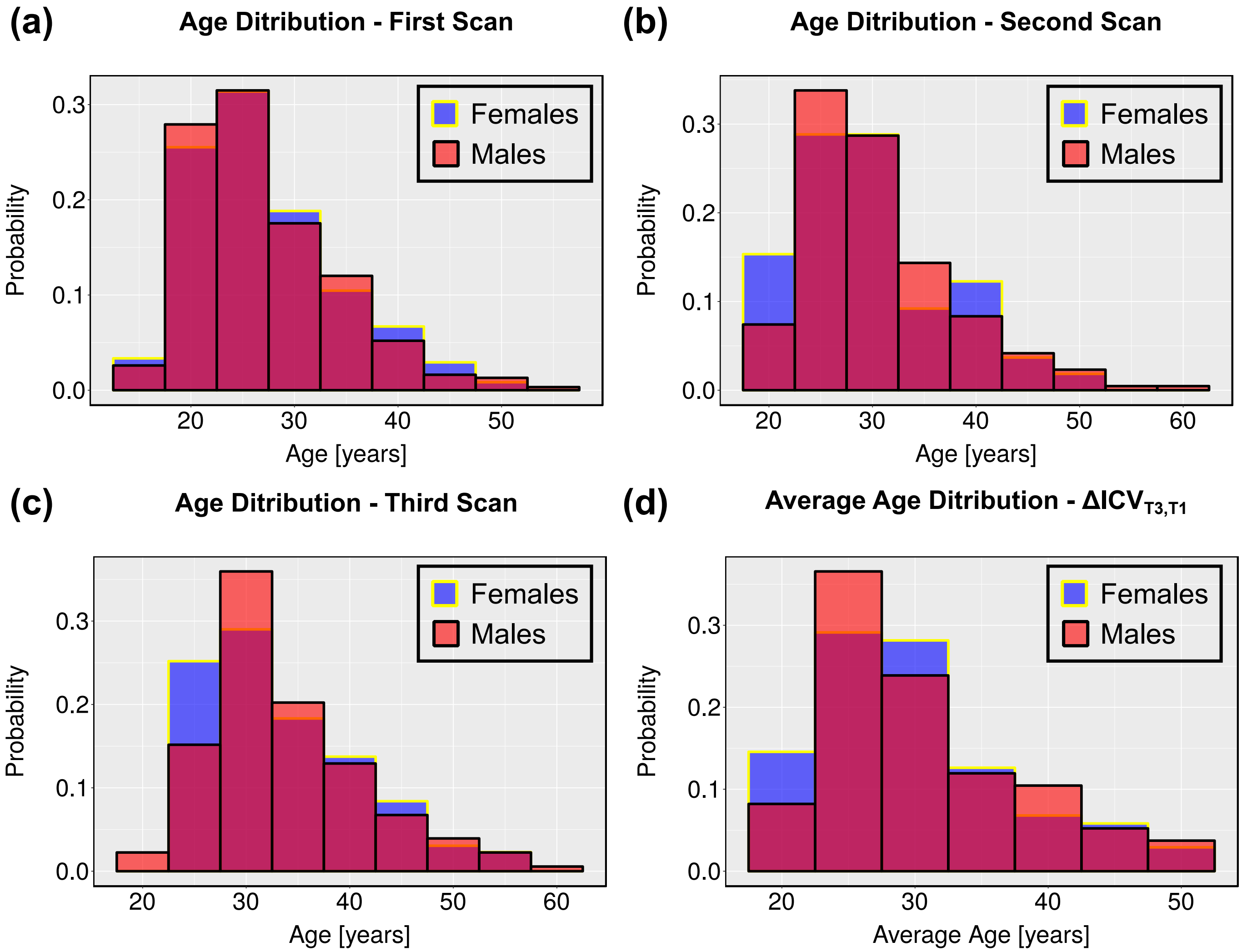


Figure S7: **Age distributions.** Age distributions for subjects that their ICV was measured at the first wave of the study(a); second wave of the study (b) and third wave of the study (c). (d) Distribution of the average age for subjects that their ICV was repeatedly measured at the first and third scans. Red - Males; Blue - Females.

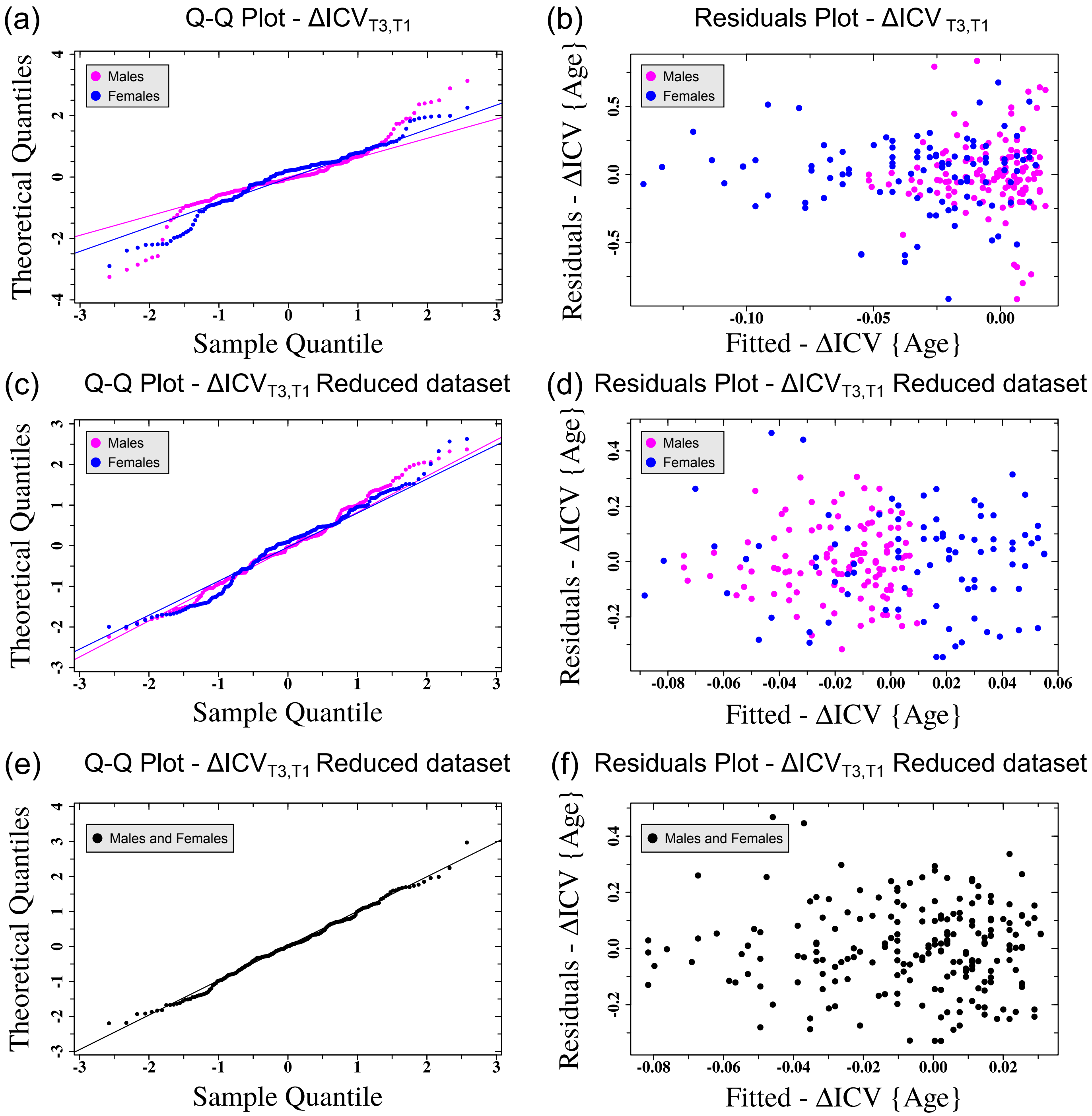


Figure S8: **Residuals analysis of $\Delta\mathrm{ICV}_{\mathrm{T3,T1}}$ as a function of age** (accompanying Fig. 2(b,d) of the main text). (a,c,e) Quantile-Quantile residuals plots and (b,d,f) residuals vs. fitted values plots for the linear fits of $\{\Delta\mathrm{ICV}_{\mathrm{T3,T1}}\}$ as a function of age. (a,b) Residuals analysis of $\{\Delta\mathrm{ICV}_{\mathrm{T3,T1}}\}$ age dependency fits for the complete dataset for females (blue) and males (pink). (c,d) Resuduals analysis of $\{\Delta\mathrm{ICV}_{\mathrm{T3,T1}}\}$ age dependency fits for the reduced dataset for females (blue) and males (pink). (e,f) Resuduals analysis of $\{\Delta\mathrm{ICV}_{\mathrm{T3,T1}}\}$ age dependency fit for the reduced dataset for males and females together.

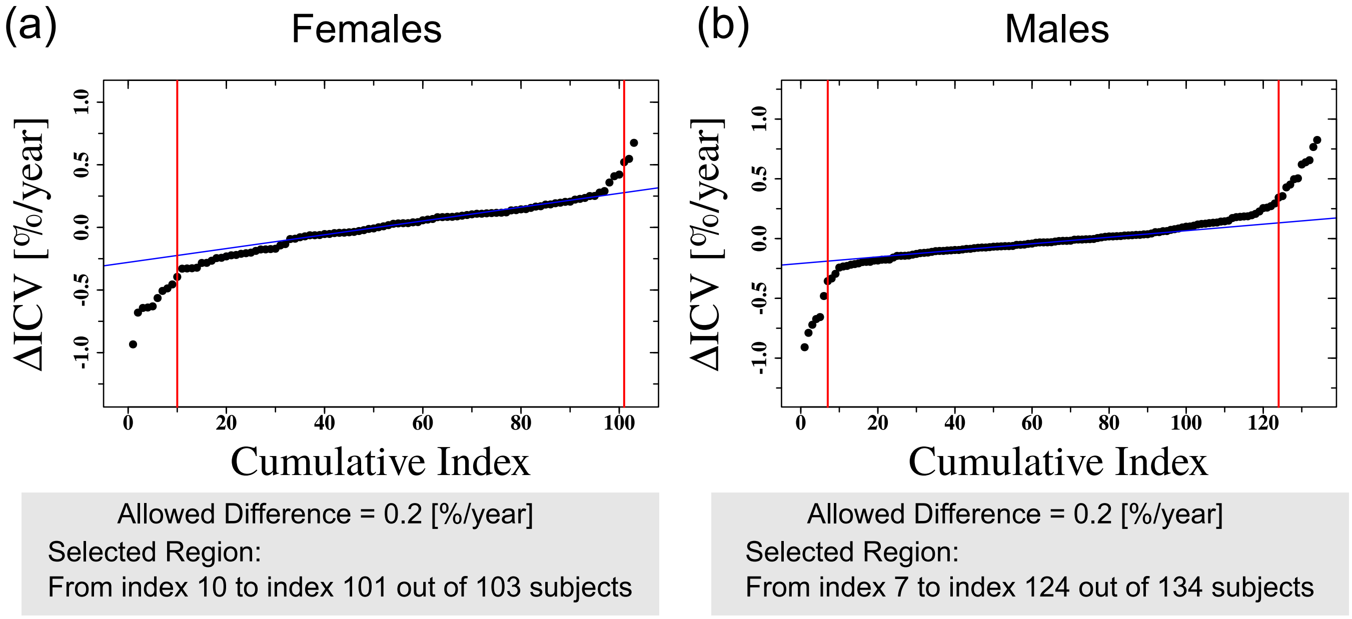


Figure S9: **Cumulative analysis of $\{\Delta \mathbf{ICV_{T3,T1}}\}$**. Cumulative results for $\{\Delta \mathrm{ICV_{T3,T1}}\}$ according to value (accompanying Fig. 2(c) of the main text). Blue lines - linear fits to the central part of the cumulative results. Red lines represents the start and end indexes of the reduced $\{\Delta \mathrm{ICV_{T3,T1}}\}$ dataset. (a) Females; (b) Males.

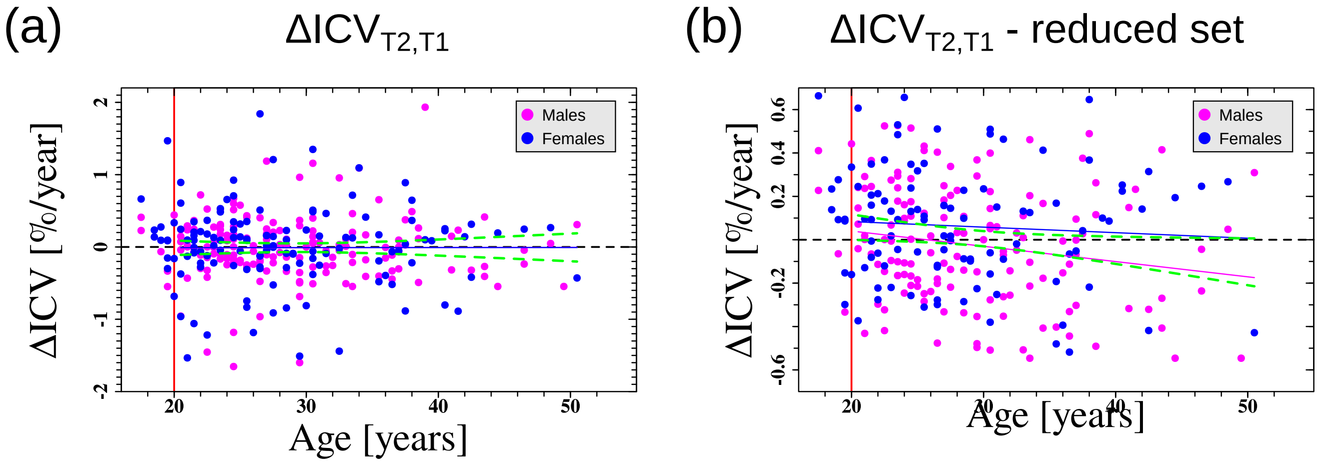


| | Longitudinal ΔICV | | | Longitudinal ΔICV - Reduced Set | | |
|---|---|---|---|---|---|---|
| | Slope [%/year$^2$] | $R^2$ | p (incline) | Slope [%/year$^2$] | $R^2$ | p (incline) |
| Females | 7e-5±7e-3 | -0.009 | 0.992 | -3e-3±4e-3 | -0.006 | 0.520 |
| Males | 3e-4±5e-3 | -0.006 | 0.955 | -7e-3±3.2e-3 | 0.026 | 0.031 |
| Both | 2e-4±4e-3 | -0.004 | 0.964 | -5e-3±2.5e-3 | 0.015 | 0.037 |

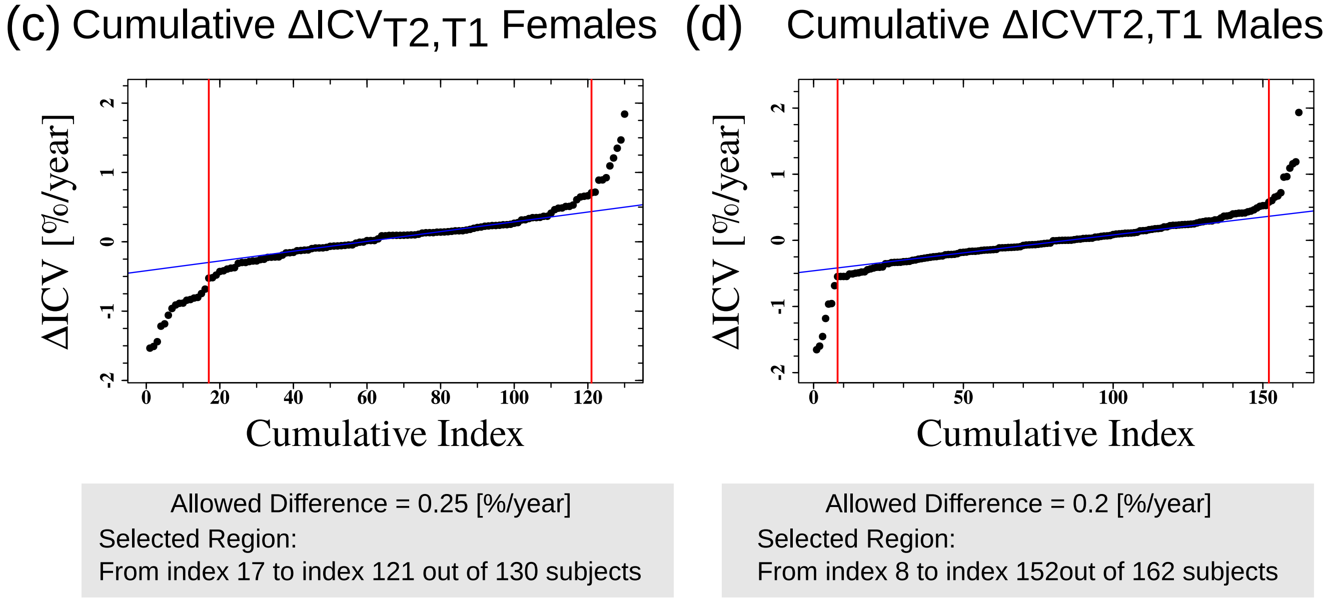


Figure S10: **Changes in the ICV between the first and second waves of the study.** $\Delta ICV_{T2,T1}$ as a function of age for the complete $\{\Delta ICV_{T2,T1}\}$ dataset (a) and the reduced $\{\Delta ICV_{T2,T1}\}$ dataset (b). Each dot represents a calculated value for a specific individual in our dataset. Blue and pink lines are fits of the data to linear functions. Vertical red lines represent the minimum age for inclusion in the fits (dots left to the line were not included in the fitting process). Dashed green lines are 95% confidence intervals of the fit for both females and males. For clarity, the fit for females and males together is not shown. Table below the graphs shows the fitting values for the females, males, and both females and males cases. $R^2$- goodness of fit. p-value - statistical significance of the slope parameter of the fit. Error values - ±s.e. (c) and (d) Cumulative results for $\{\Delta ICV_{T2,T1}\}$ according to value. Blue lines - linear fits to the central part of the cumulative results. Red lines - the start and end indexes of the reduced $\{\Delta ICV_{T2,T1}\}$ dataset. (c) Females; (d) Males.

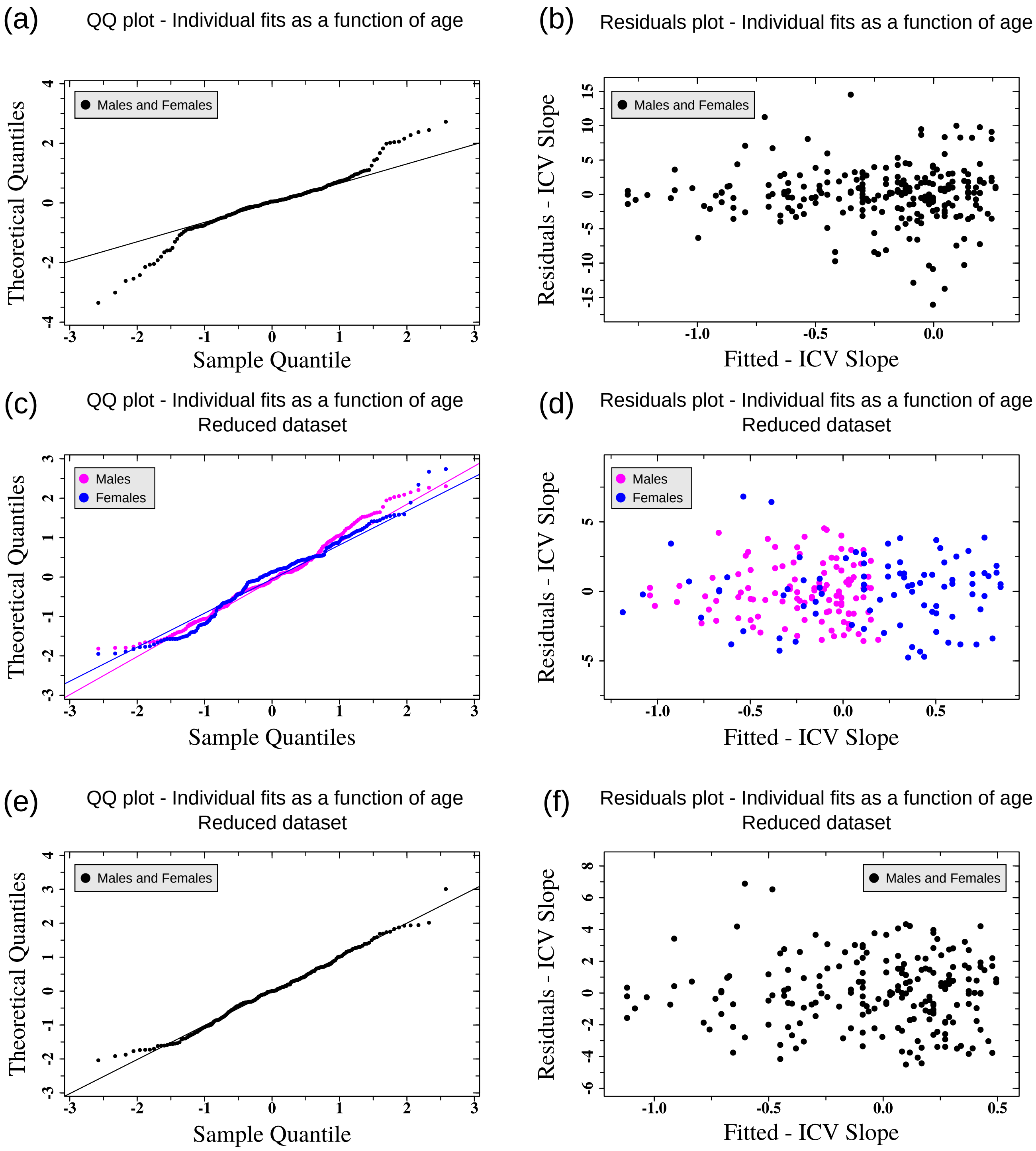


Figure S11: **Residuals analysis for the individual longitudinal ICV fits** (accompanying Fig. 3(a,c) of the main text). (a,c,e) Quantile-Quantile residuals plots and (b,d,f) residuals vs. fitted values plots for the $\{\mathrm{ICV}^{\text{Slope-fit}}_{\text{Individual}}\}$ as a function of age. (a,b) Residuals analysis of the $\{\mathrm{ICV}^{\text{Slope-fit}}_{\text{Individual}}\}$ age dependency fits for the complete dataset for females (blue) and males (pink). (c,d) Residuals analysis of the $\{\mathrm{ICV}^{\text{Slope-fit}}_{\text{Individual}}\}$ age dependency fits for the reduced dataset for females (blue) and males (pink). (e,f) Residuals analysis of the $\{\mathrm{ICV}^{\text{Slope-fit}}_{\text{Individual}}\}$ age dependency fit for the reduced dataset for males and females together.

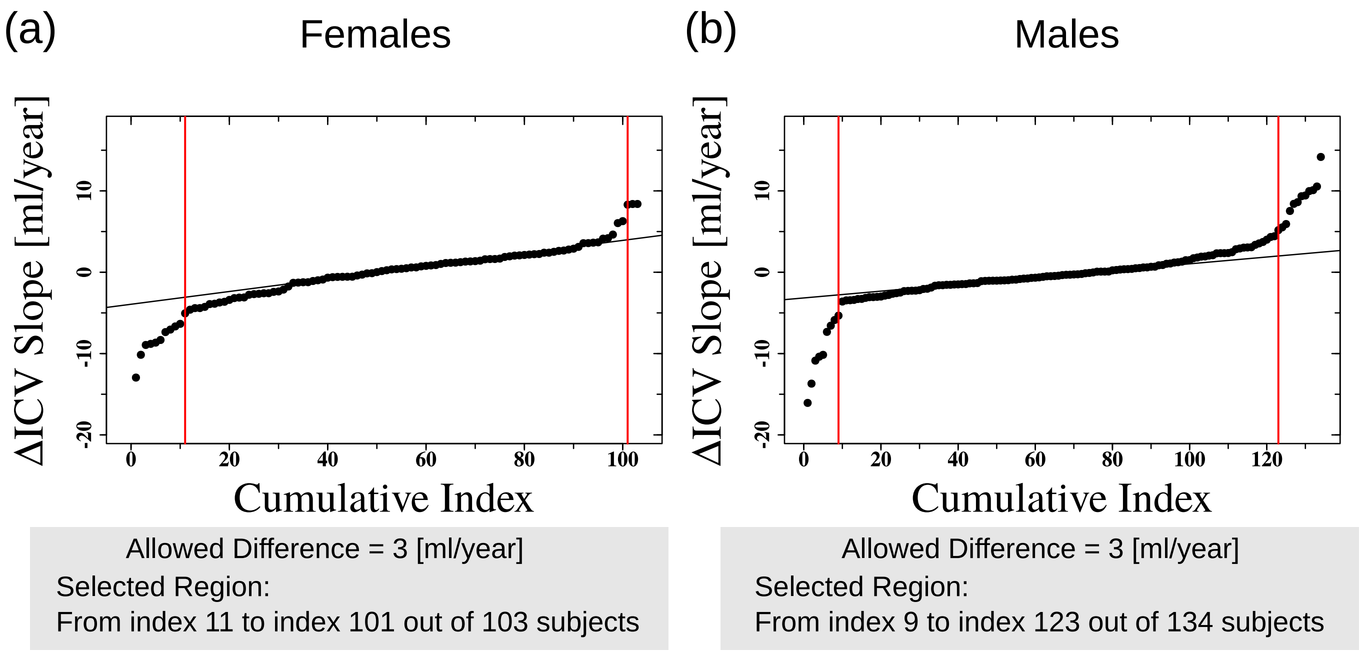


Figure S12: **Cumulative analysis of $\{\mathbf{ICV}^{\text{Slope-fit}}_{\text{Individual}}\}$.** Cumulative results for $\{\text{ICV}^{\text{Slope-fit}}_{\text{Individual}}\}$ according to value (accompanying Fig. 3(b) of the main text). Blue lines - linear fits to the central part of the cumulative results. Red lines represents the start and end indexes of the reduced $\{\text{ICV}^{\text{Slope-fit}}_{\text{Individual}}\}$ dataset. (a) Females; (b) Males.

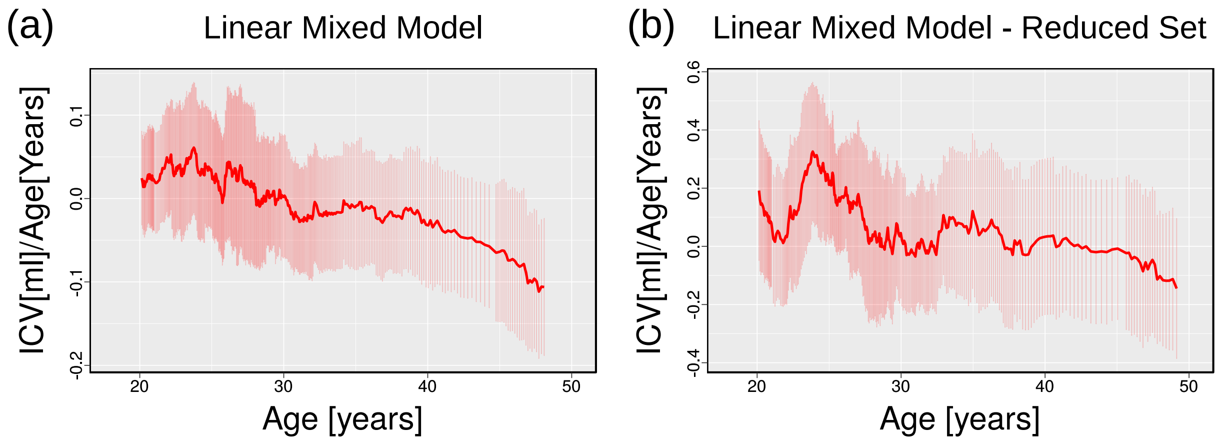


Figure S13: **Running-average results of the linear-mixed-models.** Running average graphs for $\{\text{ICV}^{\text{LMM-Slope}}_{\text{Individual}}\}$ for the complete dataset (a) and the reduced dataset (b). For the complete dataset, a sliding window of 45 subjects over the results that are shown in Fig. 4(a) of the main text was used to produce the graph. For the reduced dataset (Fig. 4(c) of the main text), a sliding window of 36 subjects was used to produce the graphs (sliding window size was changed to account for the size differences in these two datasets). Red emphasized lines - average of the sliding window. Thin red vertical lines - ±one s.d. of the sliding window.

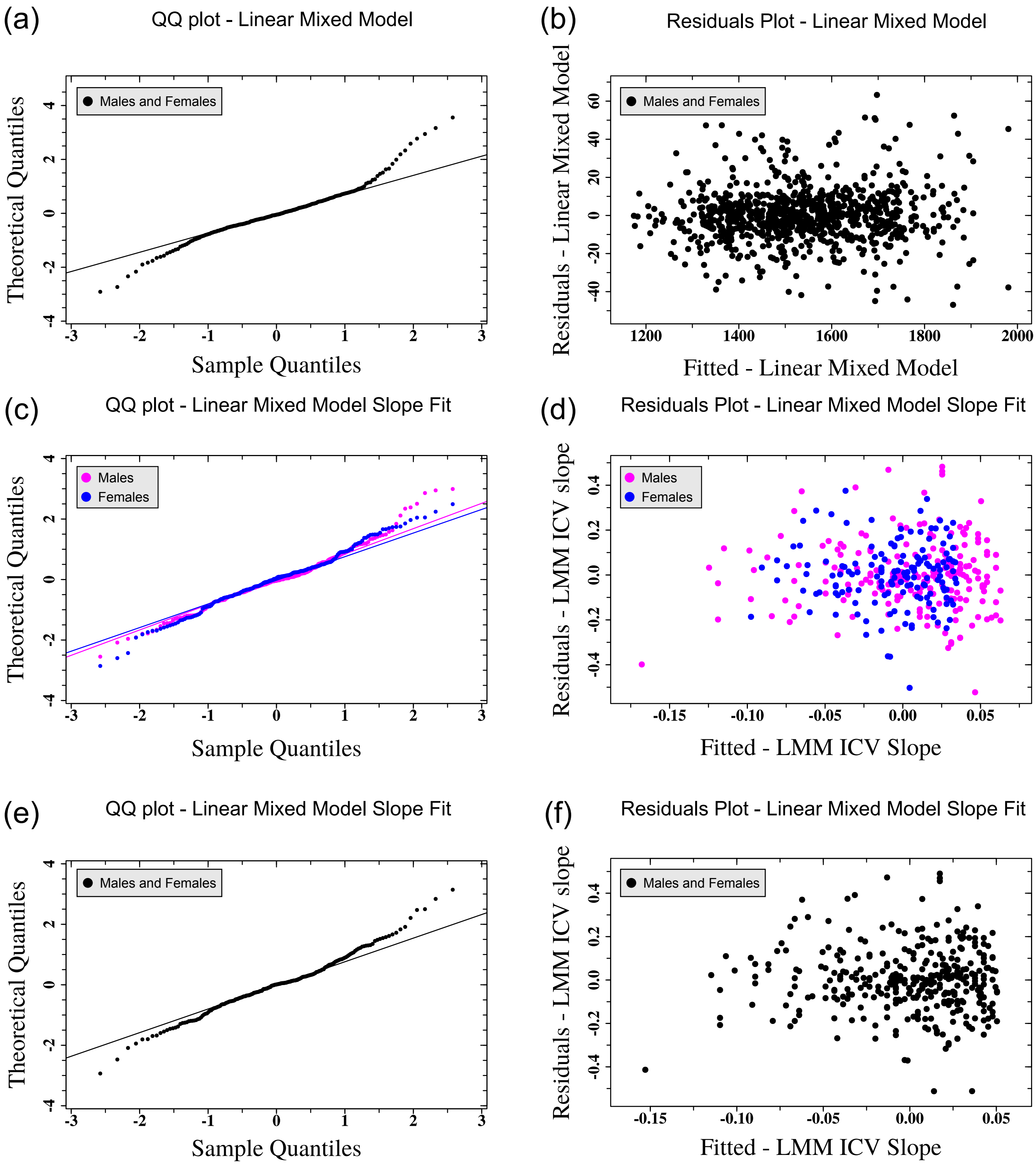


Figure S14: **Residuals analysis for the linear mixed model** (accompanying Fig. 4(a) of the main text). (a) Quantile-Quantile residuals plot and (b) residuals vs. fitted values plot for the Predicted-$\mathrm{ICV}^{\mathrm{Longitidinal}}_{\mathrm{LMM}}$ model. (c,e) Quantile-Quantile residuals plots and (d,f) residuals vs. fitted values plots for the fit of the $\{\mathrm{ICV}^{\mathrm{LMM\text{-}Slope}}_{\mathrm{Individual}}\}$ as a function of age for females (blue) and males (pink) separately (c,d), and for females and males together (e,f).

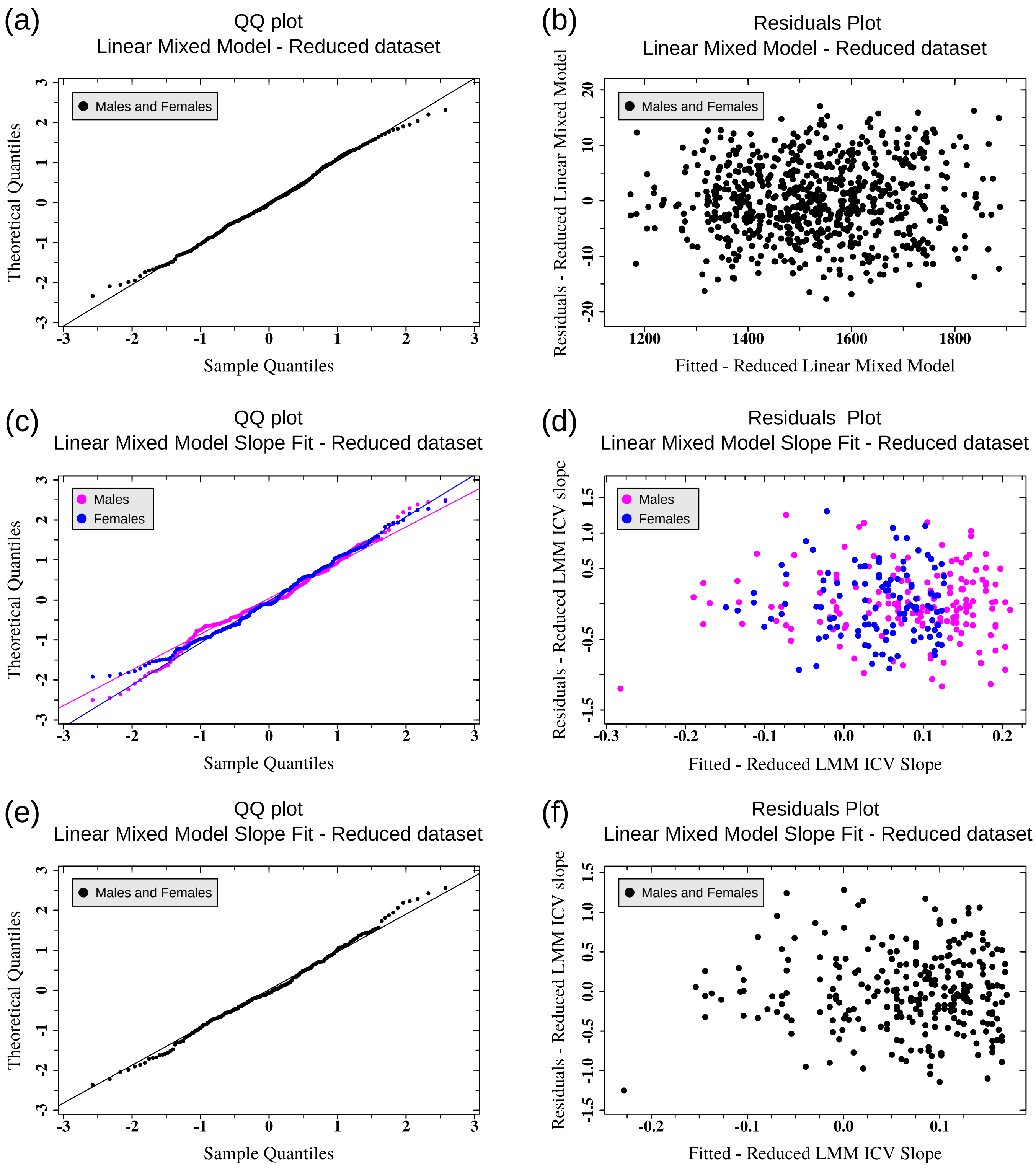


Figure S15: **Residuals analysis for the linear mixed model of the reduced dataset** (accompanying Fig. 4(c) of the main text). (a) Quantile-Quantile residuals plot and (b) residuals vs. fitted values plot for the Predicted-ICV$^{\text{Longitidinal}}_{\text{LMM}}$ model for the reduced dataset. (c,e) Quantile-Quantile residuals plots and (d,f) residuals vs. fitted values plots for the fit of the $\{\text{ICV}^{\text{LMM-Slope}}_{\text{Individual}}\}$ for the reduced dataset as a function of age for females (blue) and males (pink) separately (c,d), and for females and males together (e,f).

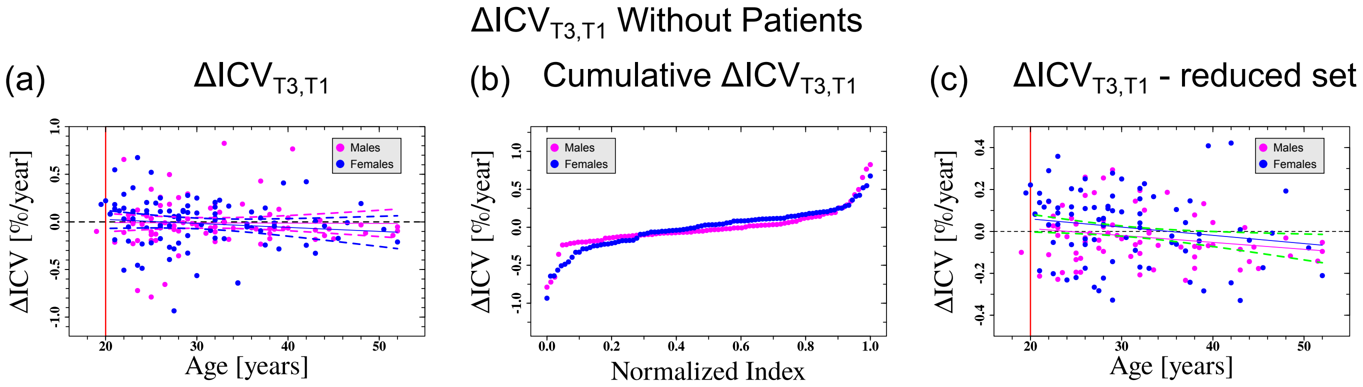


| | Longitudinal $\Delta ICV_{T3,T1}$ | | | Longitudinal $\Delta ICV_{T3,T1}$ - Reduced Set | | |
|---|---|---|---|---|---|---|
| | Slope [%/year$^2$] | $R^2$ | p (incline) | Slope [%/year$^2$] | $R^2$ | p (incline) |
| Females | -4e-3±3.8e-3 | 0.003 | 0.269 | -3.9e-3±2.5e-3 | 0.018 | 0.128 |
| Males | -3e-4±3.0e-3 | -0.012 | 0.933 | -3.3e-3±1.8e-3 | 0.034 | 0.064 |
| Both | -2e-3±2.5e-3 | -0.002 | 0.402 | -3.8e-3±1.5e-3 | 0.032 | 0.015 |

Figure S16: **Changes in the ICV as a function of age without the patient group** (accompanying Fig. 2 of the main text)**.** (a) $\Delta ICV_{T3,T1}$ as a function of $Age_{T3,T1}$ excluding the patient group. (b) Cumulative plot of $\Delta ICV_{T3,T1}$ excluding the patient group as a function of a normalized index between 0 and 1. (c) Same as (a) for a reduced dataset that is obtained by keeping only the central part of the cumulative graph in (b). Each dot represents a calculated value for a specific individual in our dataset. Blue and pink dots are for females and males respectively. blue and pink lines are fits of the data to linear functions. Vertical red line represents the minimum age for inclusion in the fits (dots left to the line were not included in the fitting process). In (c), dashed green lines are 95% confidence intervals of the fit for both females and males. For clarity, the fit itself for females and males together is not shown. The table below the graphs shows the fitting-values for the cases of females, males, and for the females and males together. $R^2$- goodness of fit. p-value - statistical significance of the slope parameter of the fit. Error values - ±s.e.

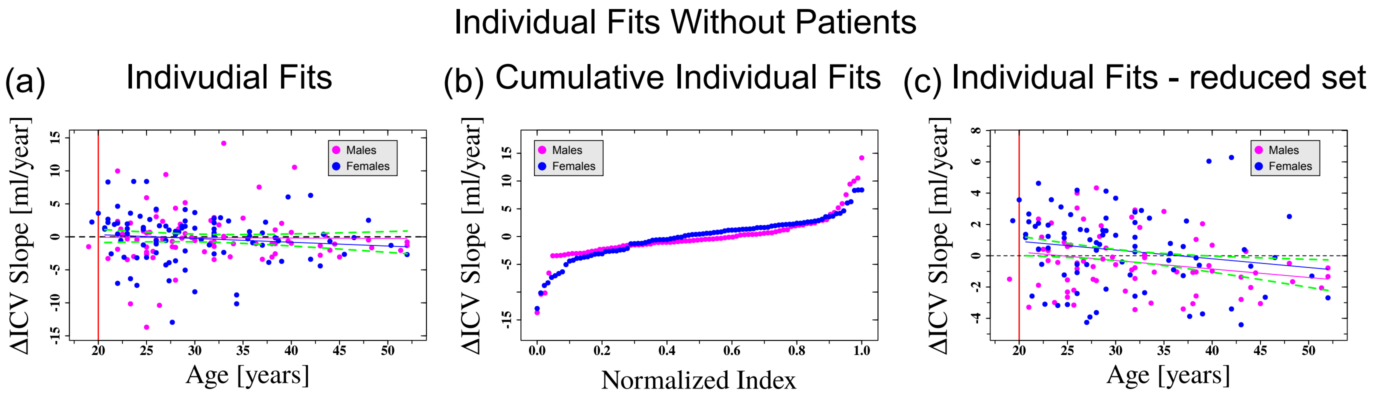


| | Individual ΔICV | | | | Individual ΔICV - Reduced Set | | | |
|---|---|---|---|---|---|---|---|---|
| | Intercept [ml/year] | Slope [ml/year$^2$] | $R^2$ | p (incline) | Intercept [ml/year] | Slope [ml/year$^2$] | $R^2$ | p (incline) |
| Females | 1.5±1.7 | -6e-2±5.5e-2 | 0.0013 | 0.293 | 2.0±1.1 | -6e-2±3.6e-2 | 0.019 | 0.128 |
| Males | -0.06±1.8 | -6e-3±6e-2 | -0.012 | 0.913 | 1.3±0.9 | -5e-2±2.6e-2 | 0.043 | 0.045 |
| Both | 0.7±1.2 | -3e-2±3.9e-2 | -0.0023 | 0.436 | -1.8±0.7 | -6e-2±2.2e-2 | 0.038 | 0.01 |

Figure S17: **Individual longitudinal fits of ICV as a function of age excluding the patient group** (accompanying Fig. 3 of the main text). (a) Values of the slopes of the individual ICV fits as a function of the average age at which the scans were made excluding the patient group. Each dot represents the age-dependent slope for an individual as was obtained from a linear model fitting in R to two or three (when available) consecutive ICV measurements. Lines are fits of the data to linear functions. Vertical red line represents the minimum age for inclusion in the fits. (b) Cumulative plot of $\{\mathrm{ICV}^{\text{Slope-fit}}_{\text{Individual}}\}$ as a function of a normalized index between 0 and 1. (c) Same as (a) for the reduced ICV{age} dataset that is obtained by keeping only the central part of the cumulative graph in (b). In (a)-(c) blue color is for females and pink for males. Dashed green lines are 95% confidence intervals of the fit for both females and males. The table below the graphs shows the fitting-values for the cases of females, males, and for the females and males together. For clarity, the fits for females and males together are not shown. $R^2$- goodness of fit. p-value - statistical significance of the slope parameter of the fit. Error values - ±s.e.

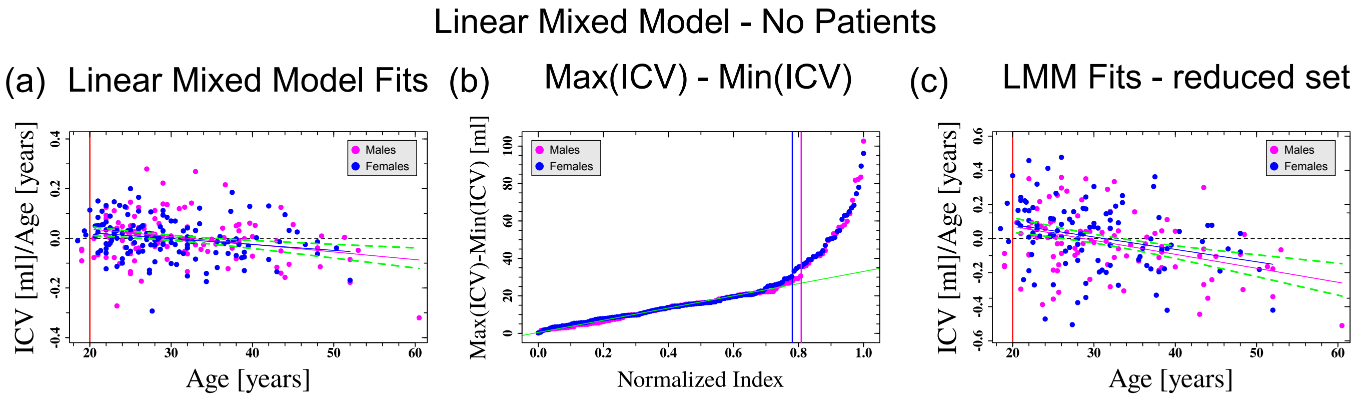


| | Linear Mixed Model Slope | | | | Linear Mixed Model Slope - Reduced Set | | | |
|---|---|---|---|---|---|---|---|---|
| | Intercept [ml/year] | Slope [ml/year$^2$] | $R^2$ | p (incline) | Intercept [ml/year] | Slope [ml/year$^2$] | $R^2$ | p (incline) |
| Females | 7.0e-2±3.9e-2 | -2e-3±1e-3 | 0.039 | 0.013 | 2e-1±8e-2 | -7e-3±2.5e-3 | 0.066 | 0.005 |
| Males | 9.8e-2±3.2e-2 | -3e-3±0.9e-3 | 0.069 | 0.002 | 2e-1±7e-2 | -8e-3±2e-3 | 0.144 | 9.8e-5 |
| Both | 8.0e-2±2.1e-2 | -3e-3±0.6e-3 | 0.057 | 8.6e-5 | 2e-1±5e-2 | -8e-3±1.5e-3 | 0.11 | 1.1e-6 |

Figure S18: **Linear mixed models for ICV as a function of age excluding the patient group** (accompanying Fig. 4 of the main text) (a) Predicted $\mathrm{ICV}^{\text{LMM-Slope}}_{\text{Individual}}$ as a function of age excluding the patient group. Each dot represents the age-dependent ICV slope for a specific individual. Lines are fits of the data to linear functions. Red line represents the minimum age for inclusion in the fits. (b) Cumulative measured individual $\mathrm{ICV}_{\max} - \mathrm{ICV}_{\min}$ as a function of a normalized index between 0 and 1 excluding the patient group. Green line - fit of the 200 subjects with the smallest $\mathrm{ICV}_{\max} - \mathrm{ICV}_{\min}$ (for both females and males) to a straight line. Vertical lines - limits of the normalized index criterion for inclusion in the linear mixed model reduced datasets for females and males. (c) same as (a) for a the reduced $\{\mathrm{ICV}^{\text{LMM-Slope}}_{\text{Individual}}\}$ dataset that was obtained by keeping only these cases left to the pink and blue lines in (b). In (a)-(c) blue color is for females and pink for males. Dashed green lines are 95% confidence intervals of the fit for both females and males. The table below the graphs shows the fitting-values for the cases of females, males, and for the females and males together. For clarity, the fits for females and males together are not shown. $R^2$- goodness of fit. p-value - statistical significance of the slope parameter of the fit. Error values - ±s.e.

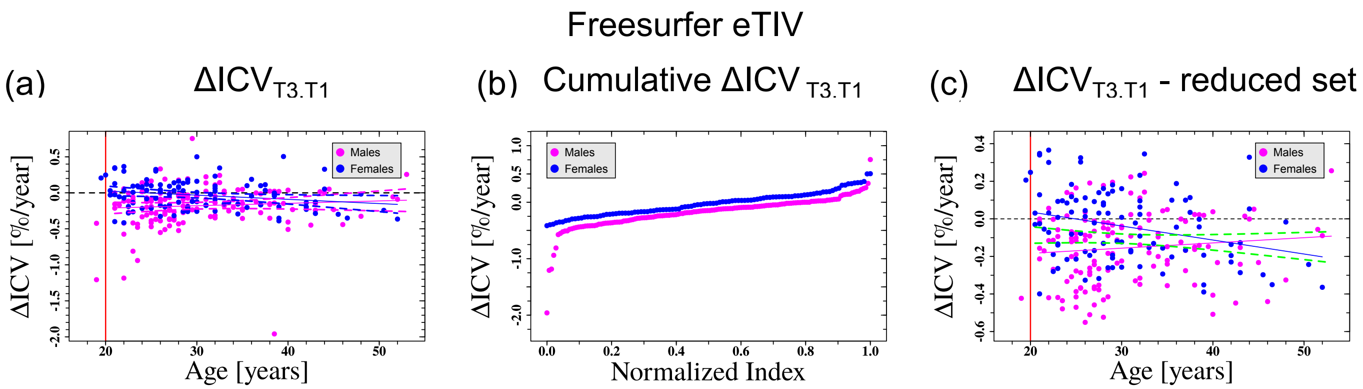


| | Longitudinal $\Delta ICV_{T3,T1}$ | | | Longitudinal $\Delta ICV_{T3,T1}$ - Reduced Set | | |
|---|---|---|---|---|---|---|
| | Slope [%/year$^2$] | $R^2$ | p (incline) | Slope [%/year$^2$] | $R^2$ | p (incline) |
| Females | -6e-3±2.6e-3 | 0.037 | 0.032 | -7e-3±2.5e-3 | 0.077 | 0.003 |
| Males | 3e-3±3.3e-3 | -7e-5 | 0.321 | 3e-3±2.2e-3 | 0.004 | 0.218 |
| Both | -0.7e-3±2.3e-3 | -0.004 | 0.740 | -2e-3±1.8e-3 | 0.001 | 0.269 |

Figure S19: **Changes in the ICV as a function of age based on the FreeSurfer eTIV values.** (a) $\Delta \mathrm{ICV}_{\mathrm{T3,T1}}$ as a function of $\mathrm{Age}_{\mathrm{T3,T1}}$ based on FreeSurfer eTIV values. (b) Cumulative plot of $\Delta \mathrm{ICV}_{\mathrm{T3,T1}}$ based on the FreeSurfer eTIV values as a function of a normalized index between 0 and 1. (c) Same as (a) for a reduced dataset that is obtained by keeping only the central part of the cumulative graph in (b). Each dot represents a calculated value for a specific individual in our dataset. Blue and pink dots are for females and males respectively. blue and pink lines are fits of the data to linear functions. Vertical red line represents the minimum age for inclusion in the fits (dots left to the line were not included in the fitting process). In (c), dashed green lines are 95% confidence intervals of the fit for both females and males. For clarity, the fit itself for females and males together is not shown. The table below the graphs shows the fitting-values for the cases of females, males, and for the females and males together. $R^2$- goodness of fit. p-value - statistical significance of the slope parameter of the fit. Error values - ±s.e.

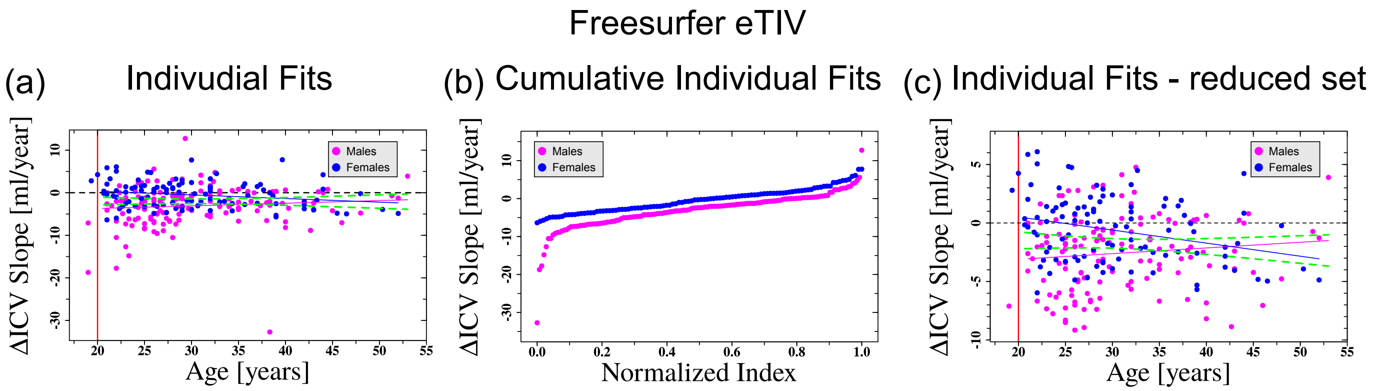


| | Individual ΔICV | | | | Individual ΔICV - Reduced Set | | | |
|---|---|---|---|---|---|---|---|---|
| | Intercept [ml/year] | Slope [ml/year$^2$] | $R^2$ | p (incline) | Intercept [ml/year] | Slope [ml/year$^2$] | $R^2$ | p (incline) |
| Females | 2.1±1.3 | -9e-2±4.1e-2 | 0.033 | 0.038 | 2.8±1.2 | -1e-1±3.7e-2 | 0.003 | 0.003 |
| Males | -4.9±1.8 | 6e-2±5.6e-2 | 0.001 | 0.274 | -4.1±1.2 | 5e-2±3.9e-2 | 0.004 | 0.210 |
| Both | -1.9±1.2 | -4e-3±3.8e-2 | -0.004 | 0.902 | -1±9.0e-1 | -3e-2±2.8e-2 | -0.0008 | 0.365 |

Figure S20: **Individual longitudinal fits of ICV as a function of age based on the FreeSurfer eTIV values.** (a) Values of the slopes of the individual ICV fits as a function of the average age at which the scans were made based on the FreeSurfer eTIV values. Each dot represents the age-dependent slope for an individual as was obtained from a linear model fitting in R to two or three (when available) consecutive ICV measurements. Lines are fits of the data to linear functions. Vertical red line represents the minimum age for inclusion in the fits. (b) Cumulative plot of $\{\mathrm{ICV}^{\text{Slope-fit}}_{\text{Individual}}\}$ as a function of a normalized index between 0 and 1. (c) Same as (a) for the reduced ICV{age} dataset that is obtained by keeping only the central part of the cumulative graph in (b). In (a)-(c) blue color is for females and pink for males. Dashed green lines are 95% confidence intervals of the fit for both females and males. The table below the graphs shows the fitting-values for the cases of females, males, and for the females and males together. For clarity, the fits for females and males together are not shown. $R^2$- goodness of fit. p-value - statistical significance of the slope parameter of the fit. Error values - ±s.e.

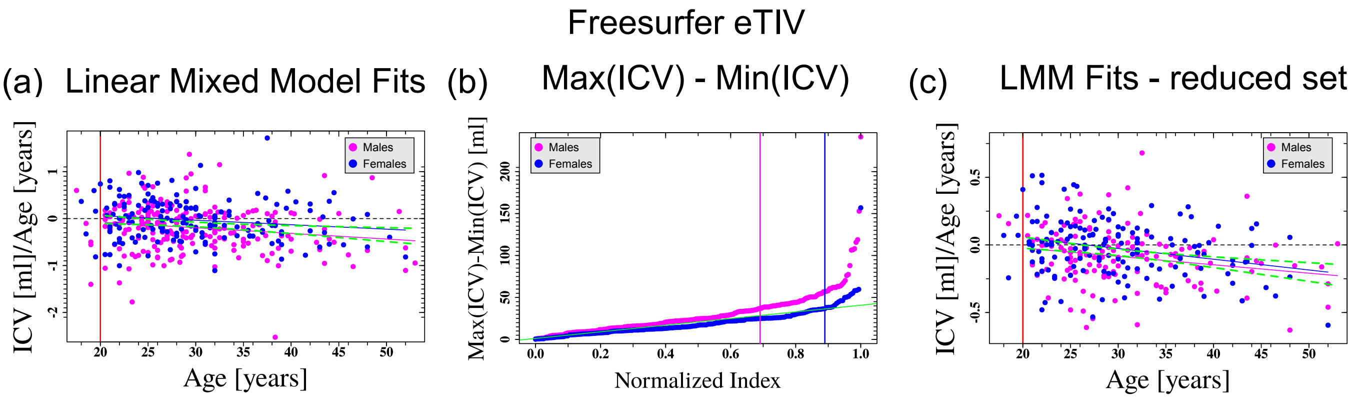


| | Linear Mixed Model Slope | | | | Linear Mixed Model Slope - Reduced Set | | | |
|---|---|---|---|---|---|---|---|---|
| | Intercept [ml/year] | Slope [ml/year$^2$] | $R^2$ | p (incline) | Intercept [ml/year] | Slope [ml/year$^2$] | $R^2$ | p (incline) |
| Females | 2.5e-1±1.5e-1 | -1e-2±0.4e-2 | 0.018 | 0.050 | 2e-1±7e-2 | -8e-3±2e-3 | 0.067 | 0.001 |
| Males | 1.7e-1±1.5e-1 | -1.2e-2±0.5e-2 | 0.023 | 0.017 | 1e-1±7e-2 | -6e-3±2.3e-3 | 0.041 | 0.009 |
| Both | 2e-1±1.1e-1 | -1e-2±0.3e-2 | 0.022 | 0.025 | 1.6e-1±5.3e-2 | -7e-3±1.6e-3 | 0.059 | 2.3e-5 |

Figure S21: **Linear mixed models for ICV as a function of age based on the FreeSurfer eTIV vaules.** (a) Predicted $\mathrm{ICV}^{\text{LMM-Slope}}_{\text{Individual}}$ as a function of age based on the FreeSurfer eTIV vales. Each dot represents the age-dependent ICV slope for a specific individual. Lines are fits of the data to linear functions. Red line represents the minimum age for inclusion in the fits. (b) Cumulative measured individual $\mathrm{ICV}_{\mathrm{max}} - \mathrm{ICV}_{\mathrm{min}}$ as a function of a normalized index between 0 and 1 based on the FreeSurfer eTIV values. Green line - fit of the 200 subjects with the smallest $\mathrm{ICV}_{\mathrm{max}} - \mathrm{ICV}_{\mathrm{min}}$ (for both females and males) to a straight line. Vertical lines - limits of the normalized index criterion for inclusion in the linear mixed model reduced datasets for females and males. (c) same as (a) for a the reduced $\{\mathrm{ICV}^{\text{LMM-Slope}}_{\text{Individual}}\}$ dataset that was obtained by keeping only these cases left to the pink and blue lines in (b). In (a)-(c) blue color is for females and pink for males. Dashed green lines are 95% confidence intervals of the fit for both females and males. The table below the graphs shows the fitting-values for the cases of females, males, and for the females and males together. For clarity, the fits for females and males together are not shown. $R^2$- goodness of fit. p-value - statistical significance of the slope parameter of the fit. Error values - ±s.e.

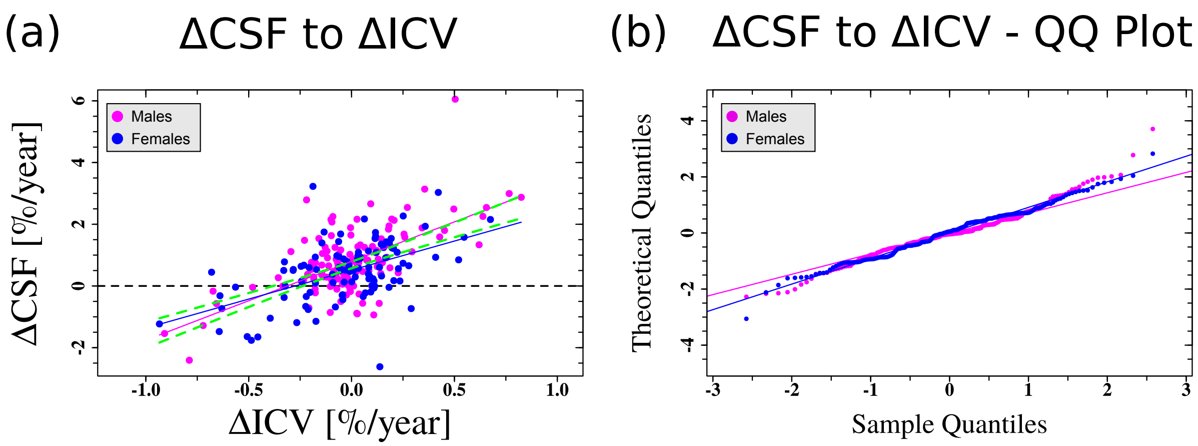


| | ΔCSF - ΔICV | | | | Pearson Correlation | | |
|---|---|---|---|---|---|---|---|
| | Intercept [%/year] | Slope | $R^2$ | p | $P_{corr}$ | 95% CI - $P_{corr}$ | p($P_{corr}$) |
| Females | 0.52±0.08 | 1.9±0.31 | 0.260 | 2.3e-8 | 0.518 | 0.359-0.646 | 2.3e-8 |
| Males | 0.80±0.07 | 2.6±0.27 | 0.390 | 4.7e-16 | 0.628 | 0.513-0.721 | 4.7e-16 |
| Both | 0.68±0.05 | 2.3±0.21 | 0.322 | <2.2e-16 | 0.579 | 0.587-0.658 | <2.2e-16 |

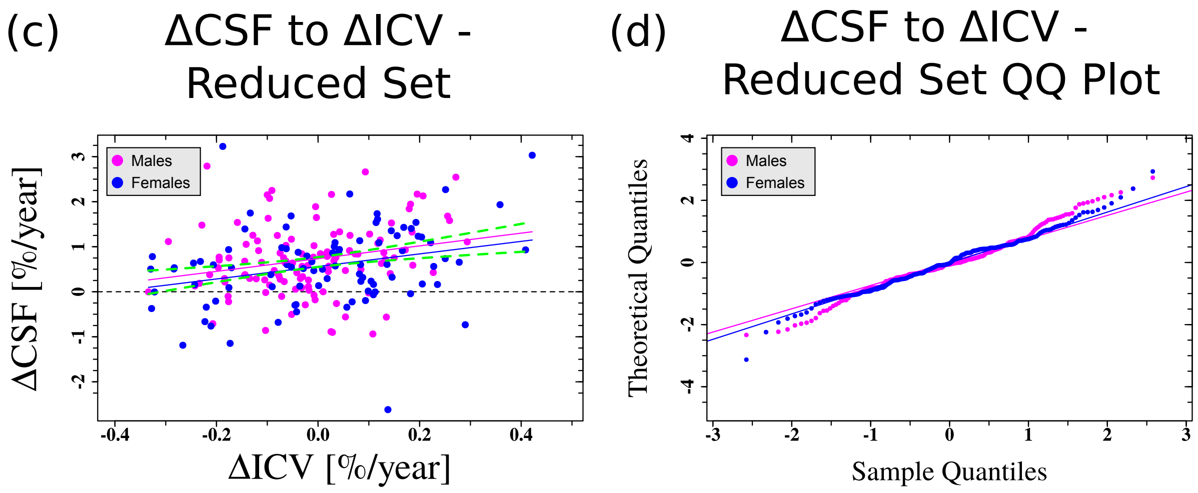


| | ΔCSF - ΔICV - Reduced Set | | | | Pearson Correlation - Reduced Set | | |
|---|---|---|---|---|---|---|---|
| | Intercept [%/year] | Slope | $R^2$ | p | $P_{corr}$ | 95% CI - $P_{corr}$ | p($P_{corr}$) |
| Females | 0.58±0.09 | 1.4±0.51 | 0.0682 | 0.0074 | 0.281 | 0.078-0.461 | 0.0074 |
| Males | 0.74±0.07 | 1.4±0.74 | 0.0486 | 0.0098 | 0.239 | 0.059-0.404 | 0.0098 |
| Both | 0.66±0.06 | 1.3±0.36 | 0.0581 | 0.0003 | 0.250 | 0.118-0.374 | 0.0003 |

Figure S22: **Correlations between the CSF and ICV**. Pearson correlation-coefficients and linear fit between $\Delta\text{CSF}_{T3,T1}$ and $\Delta\text{ICV}_{T3,T1}$. (a) Linear fit for the complete dataset. (b) Quantile-Quantile plot of the linear fit of the complete dataset. (c) Linear fit for a reduced dataset that was constructed based on the criteria of Fig. S9. (d) Quantile-Quantile plot of the linear fit of the reduced dataset. Blue color is for females and pink for males. Dashed green lines are 95% confidence intervals of the fit for both females and males. The table below the graphs shows the fitting-values for the cases of females, males, and for the females and males together. For clarity, the fits for females and males together are not shown. $R^2$- goodness of fit. p-value - statistical significance of the fit (or the Pearson correlation coefficient - $P_{corr}$). CI - confidence interval. Error values - ±s.e.

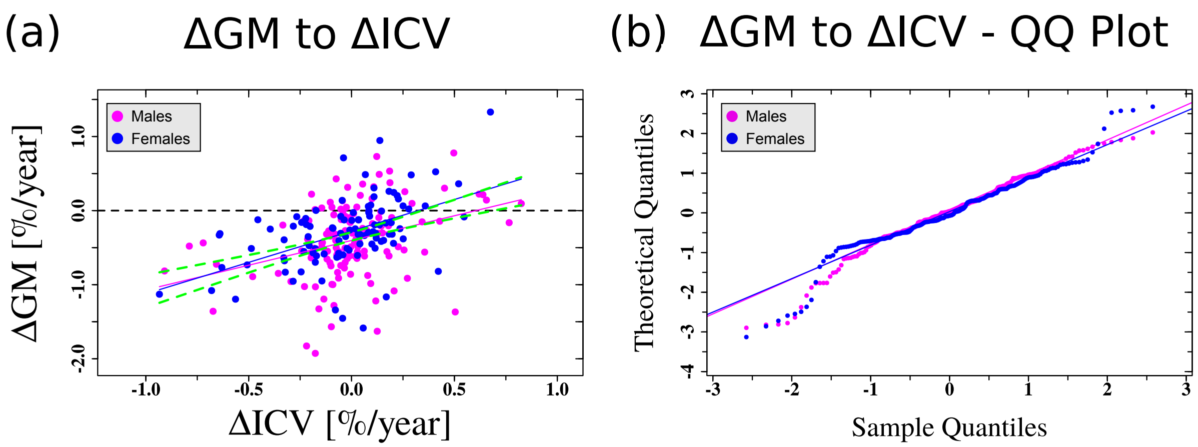


| | ΔGM - ΔICV | | | | Pearson Correlation | | |
|---|---|---|---|---|---|---|---|
| | Intercept [%/year] | Slope | $R^2$ | p | $P_{corr}$ | 95% CI - $P_{corr}$ | $p(P_{corr})$ |
| Females | -0.3±0.04 | 0.8±0.15 | 0.241 | 8.5e-8 | 0.498 | 0.337-0.631 | 8.5e-8 |
| Males | -0.4±0.04 | 0.7±0.16 | 0.114 | 3.9e-5 | 0.348 | 0.189-0.488 | 3.9e-5 |
| Both | -0.3±0.03 | 0.7±0.11 | 0.159 | 1.1e-10 | 0.403 | 0.291-0.505 | 1.1e-10 |

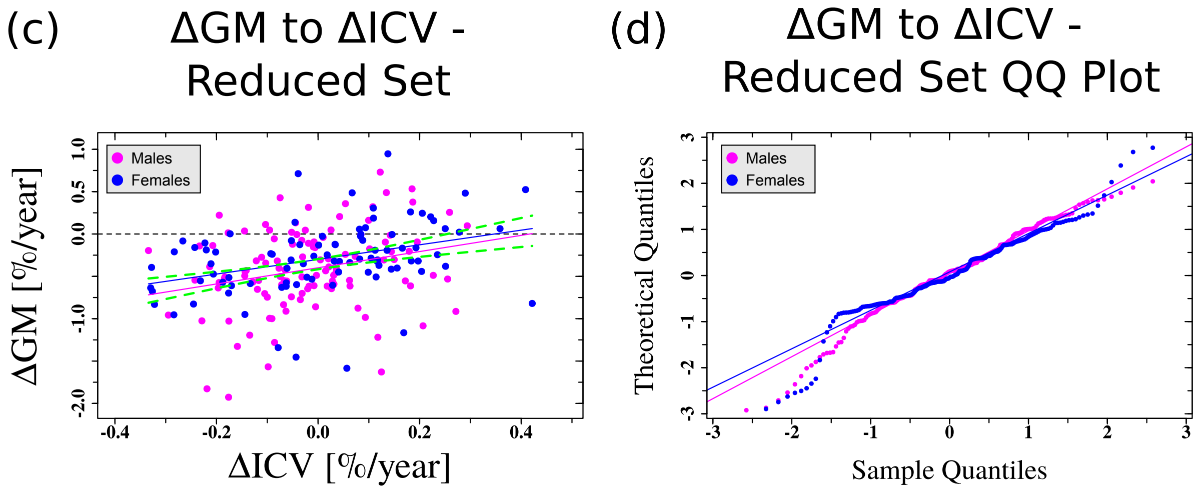


| | ΔGM - ΔICV - Reduced Set | | | | Pearson Correlation - Reduced Set | | |
|---|---|---|---|---|---|---|---|
| | Intercept [%/year] | Slope | $R^2$ | p | $P_{corr}$ | 95% CI - $P_{corr}$ | $p(P_{corr})$ |
| Females | -0.3±0.04 | 0.9±0.24 | 0.121 | 0.0005 | 0.361 | 0.166-0.529 | 0.0005 |
| Males | -0.4±0.04 | 1.0±0.33 | 0.0586 | 0.0051 | 0.258 | 0.080-0.421 | 0.0051 |
| Both | -0.4±0.03 | 0.9±0.20 | 0.0918 | 5.7e-6 | 0.310 | 0.181-0.429 | 5.7e-6 |

Figure S23: **Correlations between the gray matter and ICV**. Pearson correlation-coefficients and linear fit between $\Delta GM_{T3,T1}$ and $\Delta ICV_{T3,T1}$. (a) Linear fit for the complete dataset. (b) Quantile-Quantile plot of the linear fit of the complete dataset. (c) Linear fit for a reduced dataset that was constructed based on the criteria of Fig. S9. (d) Quantile-Quantile plot of the linear fit of the reduced dataset. Blue color is for females and pink for males. Dashed green lines are 95% confidence intervals of the fit for both females and males. The table below the graphs shows the fitting-values for the cases of females, males, and for the females and males together. For clarity, the fits for females and males together are not shown. $R^2$- goodness of fit. p-value - statistical significance of the fit (or the Pearson correlation coefficient - $P_{corr}$). CI - confidence interval. Error values - ±s.e.

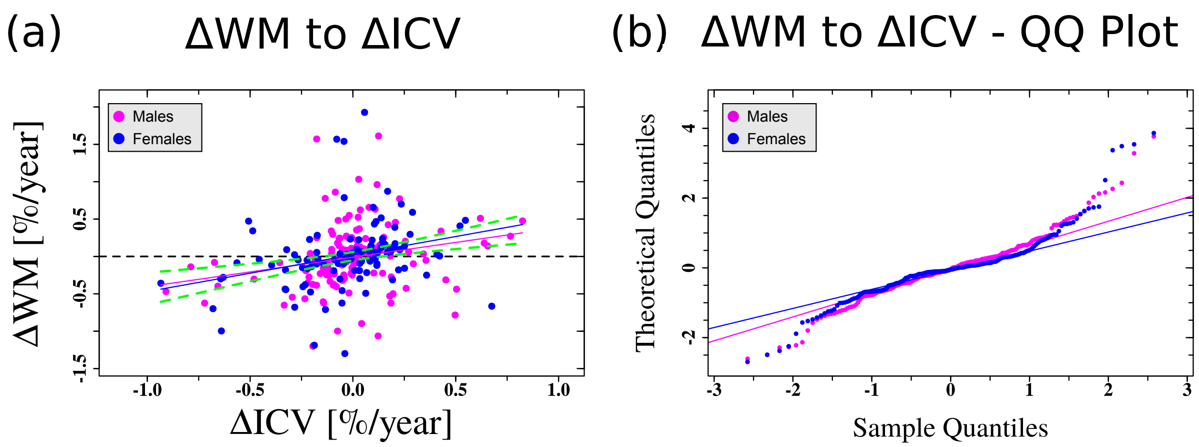


| | ΔWM - ΔICV | | | | Pearson Correlation | | |
|---|---|---|---|---|---|---|---|
| | Intercept [%/year] | Slope | $R^2$ | p | $P_{corr}$ | 95% CI - $P_{corr}$ | $p(P_{corr})$ |
| Females | 0.02±0.04 | 0.5±0.16 | 0.072 | 0.004 | 0.284 | 0.096-0.453 | 0.004 |
| Males | -0.01±0.04 | 0.4±0.14 | 0.047 | 0.007 | 0.234 | 0.066-0.387 | 0.007 |
| Both | 0.03±0.3 | 0.4±0.11 | 0.061 | 7e-5 | 0.255 | 0.132-0.371 | 7e-5 |

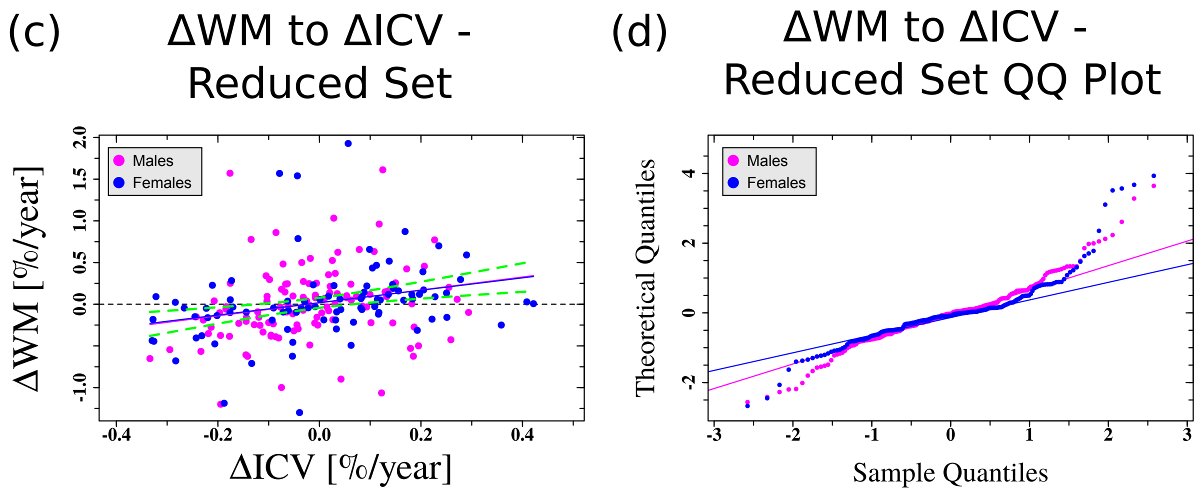


| | ΔWM - ΔICV - Reduced Set | | | | Pearson Correlation - Reduced Set | | |
|---|---|---|---|---|---|---|---|
| | Intercept [%/year] | Slope | $R^2$ | p | $P_{corr}$ | 95% CI - $P_{corr}$ | $p(P_{corr})$ |
| Females | 0.02±0.05 | 0.7±0.27 | 0.0692 | 0.0070 | 0.282 | 0.080-0.462 | 0.0070 |
| Males | 0.02±0.04 | 0.8±0.31 | 0.0427 | 0.0147 | 0.226 | 0.046-0.392 | 0.0147 |
| Both | 0.02±0.03 | 0.8±0.20 | 0.0603 | 0.0002 | 0.255 | 0.122-0.378 | 0.0002 |

Figure S24: **Correlations between the white matter and ICV**. Pearson correlation-coefficients and linear fit between $\Delta WM_{T3,T1}$ and $\Delta ICV_{T3,T1}$. (a) Linear fit for the complete dataset. (b) Quantile-Quantile plot of the linear fit of the complete dataset. (c) Linear fit for a reduced dataset that was constructed based on the criteria of Fig. S9. (d) Quantile-Quantile plot of the linear fit of the reduced dataset. Blue color is for females and pink for males. Dashed green lines are 95% confidence intervals of the fit for both females and males. The table below the graphs shows the fitting-values for the cases of females, males, and for the females and males together. For clarity, the fits for females and males together are not shown. $R^2$- goodness of fit. p-value - statistical significance of the fit (or the Pearson correlation coefficient - $P_{corr}$). CI - confidence interval. Error values - ±s.e.

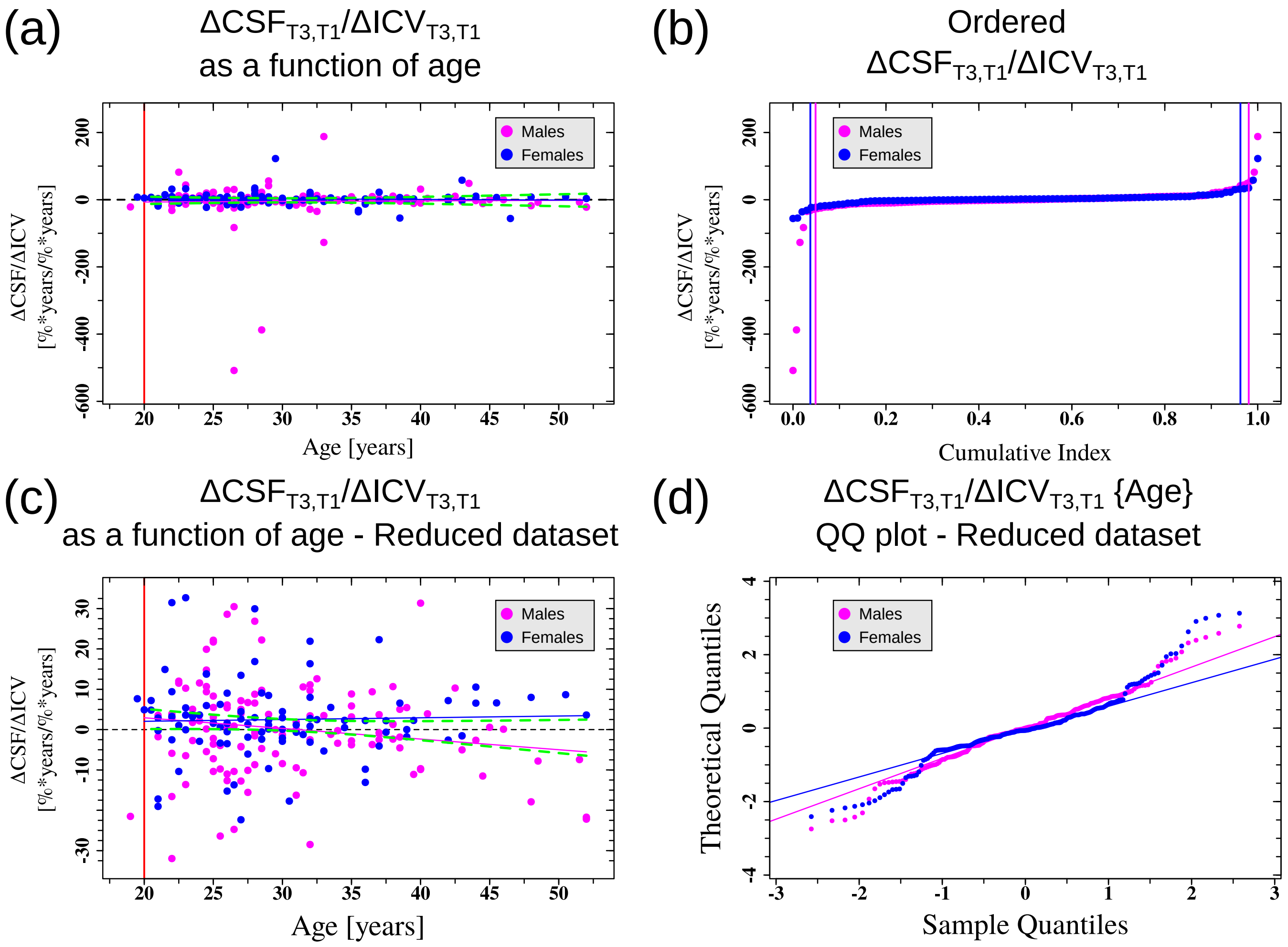


| | ΔCSF /ΔICV as a function of age | | | | ΔCSF /ΔICV as a function of age for the reduced dataset | | | |
|---|---|---|---|---|---|---|---|---|
| | Intercept | Slope [1/year] | $R^2$ | p | Intercept | Slope [1/year] | $R^2$ | p |
| Females | 8±8 | 0.2±0.3 | -0.006 | 0.521 | 1±4.3 | 0.04±0.14 | -0.001 | 0.757 |
| Males | -10±23 | 0.2±0.7 | -0.007 | 0.820 | 8±4.4 | -0.3±0.14 | 0.021 | 0.589 |
| Both | -2±14 | 0.008±0.4 | -0.004 | 0.986 | 6±3.1 | -0.1±0.1 | 0.005 | 0.146 |

Figure S25: **$\Delta$CSF$_{T3,T1}$/$\Delta$ICV$_{T3,T1}$ as a function of age. Relationships between $\Delta$CSF$_{T3,T1}$ and $\Delta$ICV$_{T3,T1}$ as a function of age.** (a) Linear fit for the complete dataset. (b) Cumulative plot of $\Delta\mathrm{CSF}_{\mathrm{T3,T1}}/\Delta\mathrm{ICV}_{\mathrm{T3,T1}}$ as a function of a normalized index between 0 and 1. Vertical lines represent the range of inclusion in the reduced dataset. (c) Linear fit for a reduced dataset that was constructed based on the criteria of panel (b). (d) Quantile-Quantile plot of the linear fit of the reduced dataset. Blue color is for females and pink for males. Dashed green lines are 95% confidence intervals of the fit for both females and males. The table below the graphs shows the fitting-values for the cases of females, males, and for the females and males together. For clarity, the fits for females and males together are not shown. $R^2$- goodness of fit. p-value - statistical significance of the fit. Error values - ±s.e.

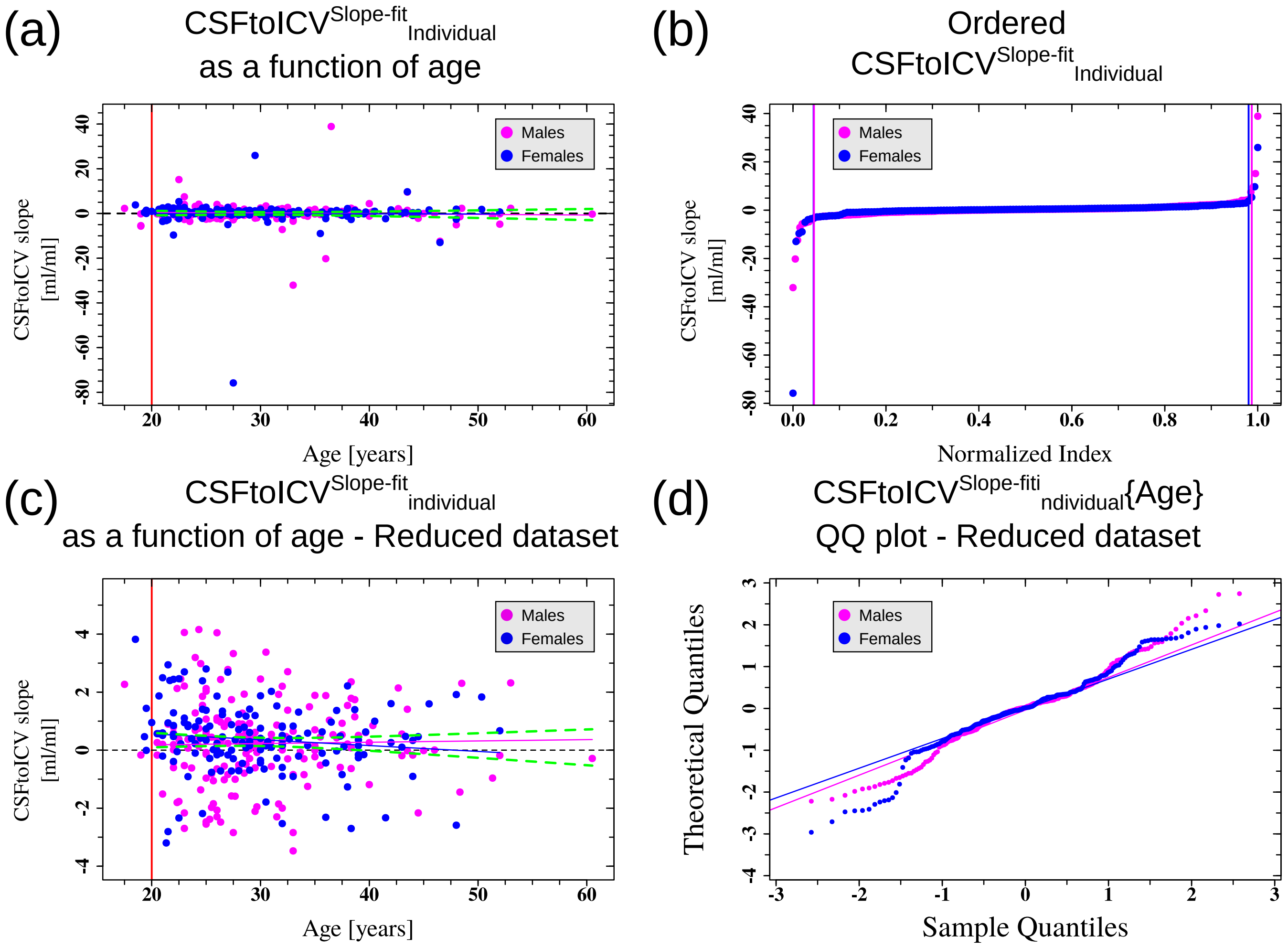


| | CSFtoICV$^{\text{Slope-fit}}_{\text{Individual}}$ as a function of age | | | | CSFtoICV$^{\text{Slope-fit}}_{\text{Individual}}$ as a function of age for the reduced dataset | | | |
|---|---|---|---|---|---|---|---|---|
| | Intercept | Slope [1/year] | $R^2$ | p | Intercept | Slope [1/year] | $R^2$ | p |
| Females | 0.2±2.4 | -0.3±0.12 | -0.007 | 0.995 | 0.9±0.43 | -0.02±0.01 | 0.008 | 0.148 |
| Males | 1.0±1.4 | -0.03±0.04 | -0.003 | 0.498 | 0.09±0.45 | 0.004±0.01 | -0.005 | 0.759 |
| Both | 0.5±1.3 | -0.02±0.04 | -0.002 | 0.692 | 0.5±0.32 | -0.01±0.03 | -0.002 | 0.542 |

Figure S26: **Slope of the CSF and ICV individual fits as a function of age.** Relationship between $\{\text{CSFtoICV}^{\text{Slope-fit}}_{\text{Individual i}}\}$ and age. (a) Linear fit for the complete dataset. (b) Cumulative plot of $\{\text{CSFtoICV}^{\text{Slope-fit}}_{\text{Individual i}}\}$ as a function of a normalized index between 0 and 1. Vertical lines represent the range of inclusion in the reduced dataset. (c) Linear fit for a reduced dataset that was constructed based on the criteria of panel (b). (d) Quantile-Quantile plot of the linear fit of the reduced dataset. Blue color is for females and pink for males. Dashed green lines are 95% confidence intervals of the fit for both females and males. The table below the graphs shows the fitting-values for the cases of females, males, and for the females and males together. For clarity, the fits for females and males together are not shown. $R^2$- goodness of fit. p-value - statistical significance of the fit. Error values - ±s.e.

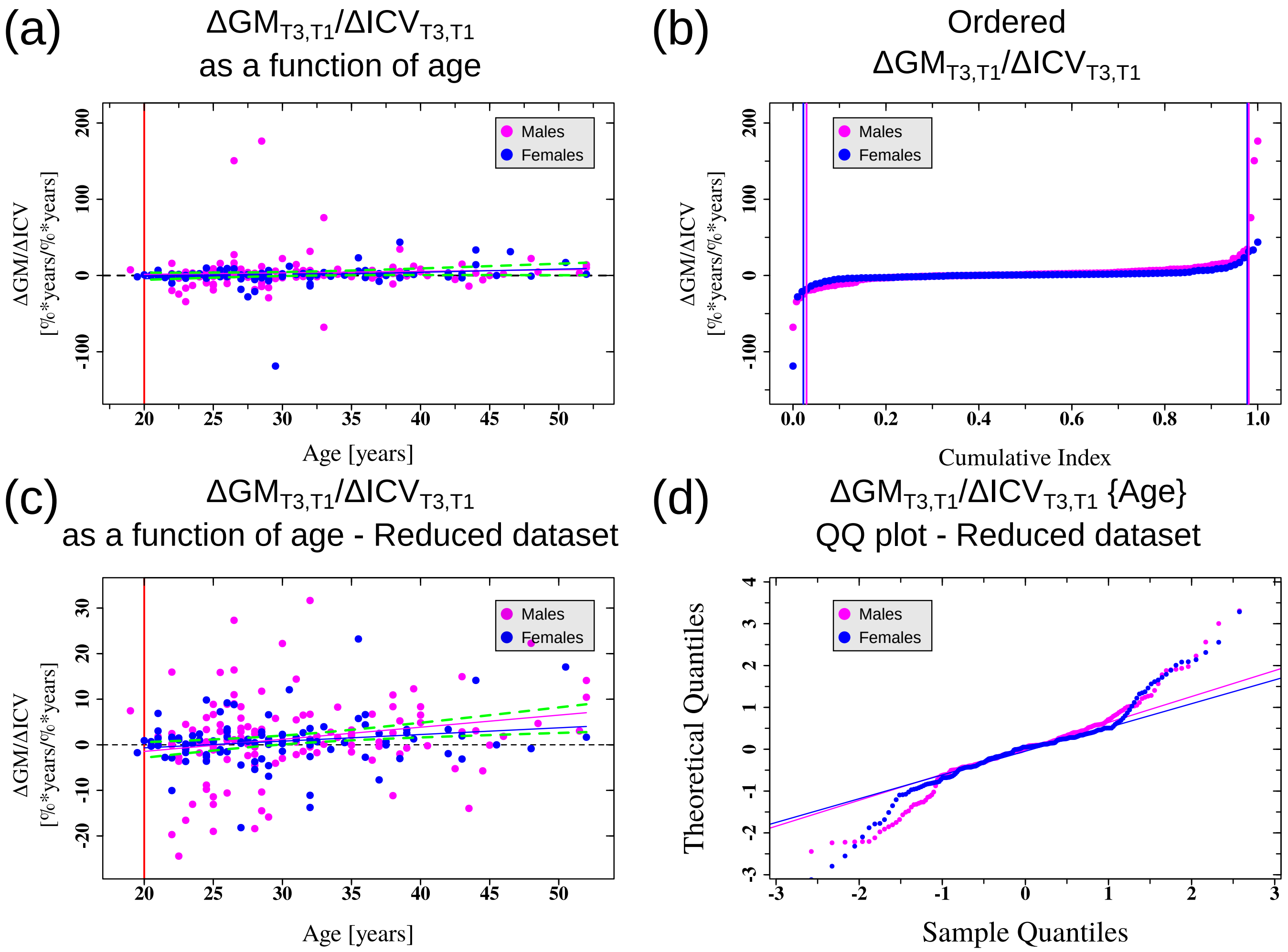


| | ΔGM/ΔICV as a function of age | | | | ΔGM/ΔICV as a function of age for the reduced dataset | | | |
|---|---|---|---|---|---|---|---|---|
| | Intercept | Slope [1/year] | $R^2$ | p | Intercept | Slope [1/year] | $R^2$ | p |
| Females | -13±6.2 | 0.4±0.20 | 0.322 | 0.040 | -4±2.4 | 0.15±0.08 | 0.026 | 0.064 |
| Males | -2±9 | 0.2±0.3 | -0.004 | 0.494 | -7±3.3 | 0.3±0.10 | 0.042 | 0.011 |
| Both | -7±5.8 | 0.3±0.18 | 0.007 | 0.107 | -5±2.1 | 0.2±0.07 | 0.004 | 0.002 |

Figure S27: **$\Delta$GM$_{T3,T1}$/$\Delta$ICV$_{T3,T1}$ as a function of age. Relationships between $\Delta$GM$_{T3,T1}$ and $\Delta$ICV$_{T3,T1}$ as a function of age.** (a) Linear fit for the complete dataset. (b) Cumulative plot of $\Delta\mathrm{GM}_{\mathrm{T3,T1}}/\Delta\mathrm{ICV}_{\mathrm{T3,T1}}$ as a function of a normalized index between 0 and 1. Vertical lines represent the range of inclusion in the reduced dataset. (c) Linear fit for a reduced dataset that was constructed based on the criteria of panel (b). (d) Quantile-Quantile plot of the linear fit of the reduced dataset. Blue color is for females and pink for males. Dashed green lines are 95% confidence intervals of the fit for both females and males. The table below the graphs shows the fitting-values for the cases of females, males, and for the females and males together. For clarity, the fits for females and males together are not shown. $R^2$- goodness of fit. p-value - statistical significance of the fit. Error values - ±s.e.

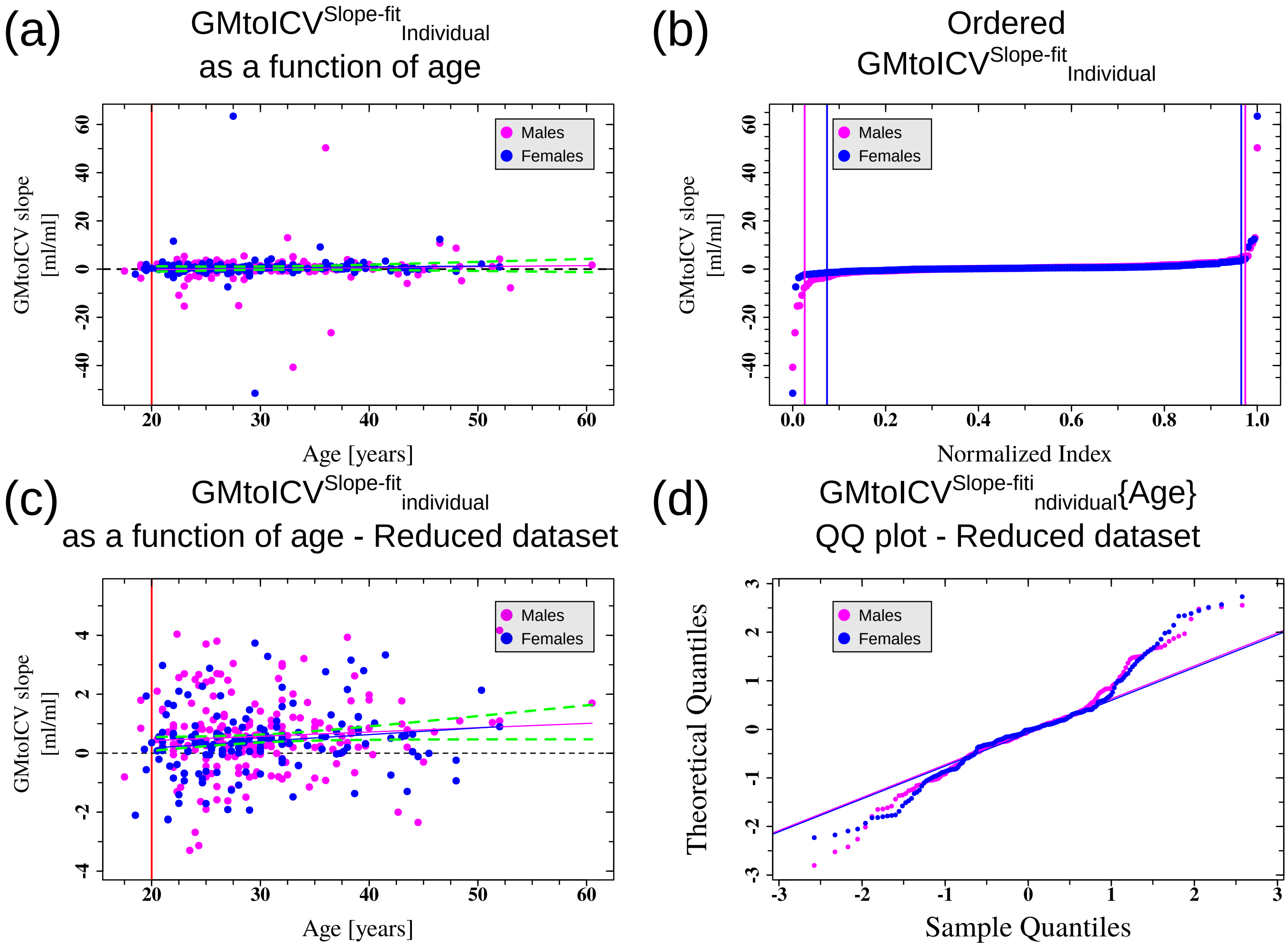


| | GMtoICV$^{\text{Slope-fit}}_{\text{Individual}}$ as a function of age | | | | GMtoICV$^{\text{Slope-fit}}_{\text{Individual}}$ as a function of age for the reduced dataset | | | |
|---|---|---|---|---|---|---|---|---|
| | Intercept | Slope [1/year] | $R^2$ | p | Intercept | Slope [1/year] | $R^2$ | p |
| Females | -0.3±2.5 | 0.03±0.08 | -0.006 | 0.690 | -0.3±0.40 | 0.02±0.01 | 0.015 | 0.080 |
| Males | -1.0±1.7 | 0.04±0.06 | -0.002 | 0.466 | 0.1±0.43 | 0.01±0.01 | 0.0005 | 0.297 |
| Both | -0.7±1.4 | 0.04±0.05 | -0.001 | 0.436 | -0.05±0.30 | 0.02±0.01 | 0.008 | 0.057 |

Figure S28: **Slope of the GM and ICV individual fits as a function of age.** Relationship between $\{\text{GMtoICV}^{\text{Slope-fit}}_{\text{Individual i}}\}$ and age. (a) Linear fit for the complete dataset. (b) Cumulative plot of $\{\text{GMtoICV}^{\text{Slope-fit}}_{\text{Individual i}}\}$ as a function of a normalized index between 0 and 1. Vertical lines represent the range of inclusion in the reduced dataset. (c) Linear fit for a reduced dataset that was constructed based on the criteria of panel (b). (d) Quantile-Quantile plot of the linear fit of the reduced dataset. Blue color is for females and pink for males. Dashed green lines are 95% confidence intervals of the fit for both females and males. The table below the graphs shows the fitting-values for the cases of females, males, and for the females and males together. For clarity, the fits for females and males together are not shown. $R^2$- goodness of fit. p-value - statistical significance of the fit. Error values - ±s.e.

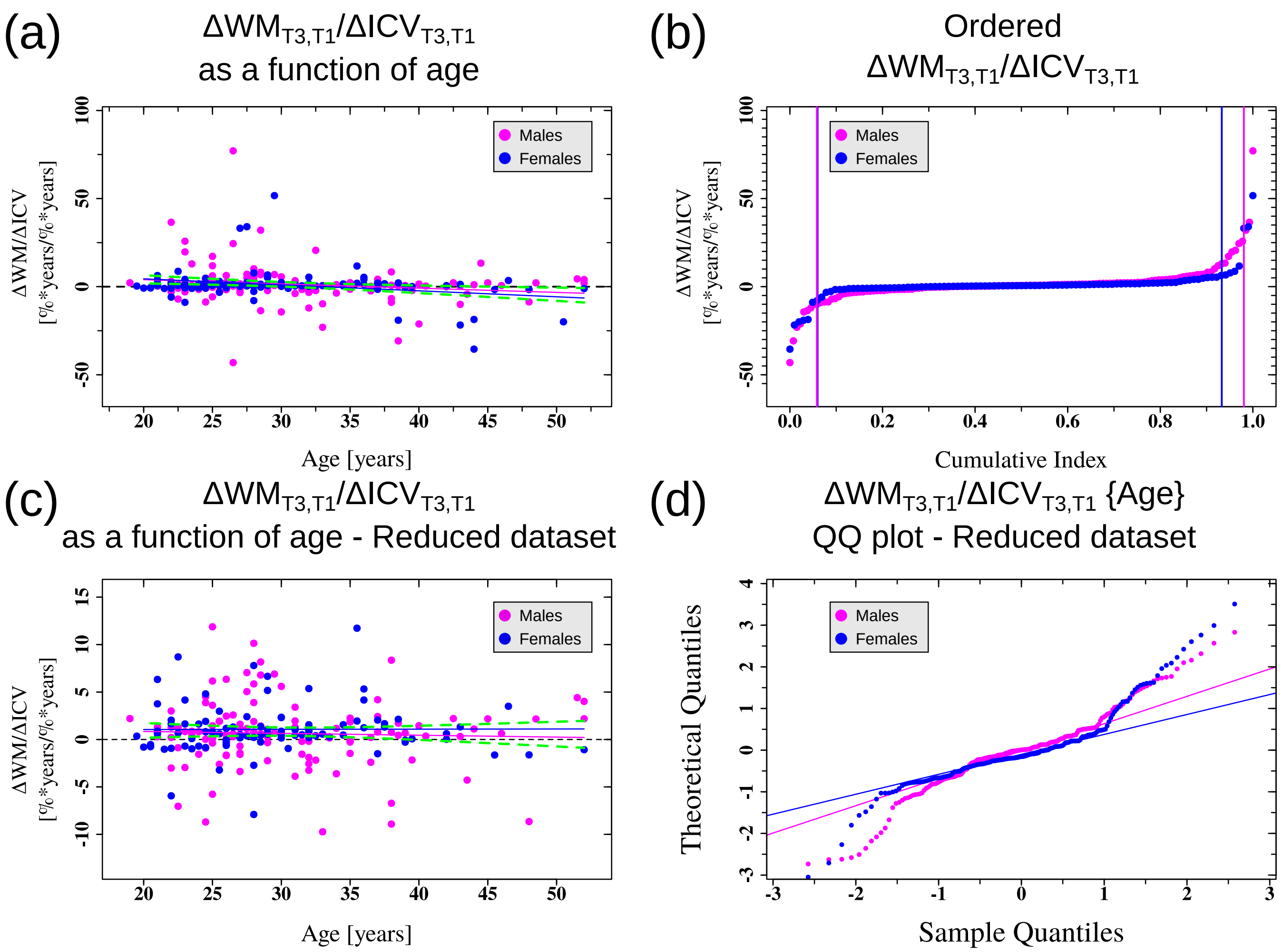


| | ΔWM/ΔICV as a function of age | | | | ΔWM/ΔICV as a function of age for the reduced dataset | | | |
|---|---|---|---|---|---|---|---|---|
| | Intercept | Slope [1/year] | $R^2$ | p | Intercept | Slope [1/year] | $R^2$ | p |
| Females | 11±3.8 | -0.3±0.12 | 0.063 | 0.007 | 1±1.3 | 0.002±0.04 | -0.011 | 0.965 |
| Males | 9±4.2 | -0.2±0.13 | 0.017 | 0.075 | 1±1.5 | -0.02±0.05 | -0.007 | 0.601 |
| Both | 10±2.9 | -0.3±0.09 | 0.035 | 0.002 | 1±1.0 | -0.01±0.03 | -0.004 | 0.679 |

Figure S29: **$\Delta$WM$_{T3,T1}$/$\Delta$ICV$_{T3,T1}$ as a function of age. Relationships between $\Delta$WM$_{T3,T1}$ and $\Delta$ICV$_{T3,T1}$ as a function of age.** (a) Linear fit for the complete dataset. (b) Cumulative plot of $\Delta\mathrm{WM}_{\mathrm{T3,T1}}/\Delta\mathrm{ICV}_{\mathrm{T3,T1}}$ as a function of a normalized index between 0 and 1. Vertical lines represent the range of inclusion in the reduced dataset. (c) Linear fit for a reduced dataset that was constructed based on the criteria of panel (b). (d) Quantile-Quantile plot of the linear fit of the reduced dataset. Blue color is for females and pink for males. Dashed green lines are 95% confidence intervals of the fit for both females and males. The table below the graphs shows the fitting-values for the cases of females, males, and for the females and males together. For clarity, the fits for females and males together are not shown. $R^2$- goodness of fit. p-value - statistical significance of the fit. Error values - ±s.e.

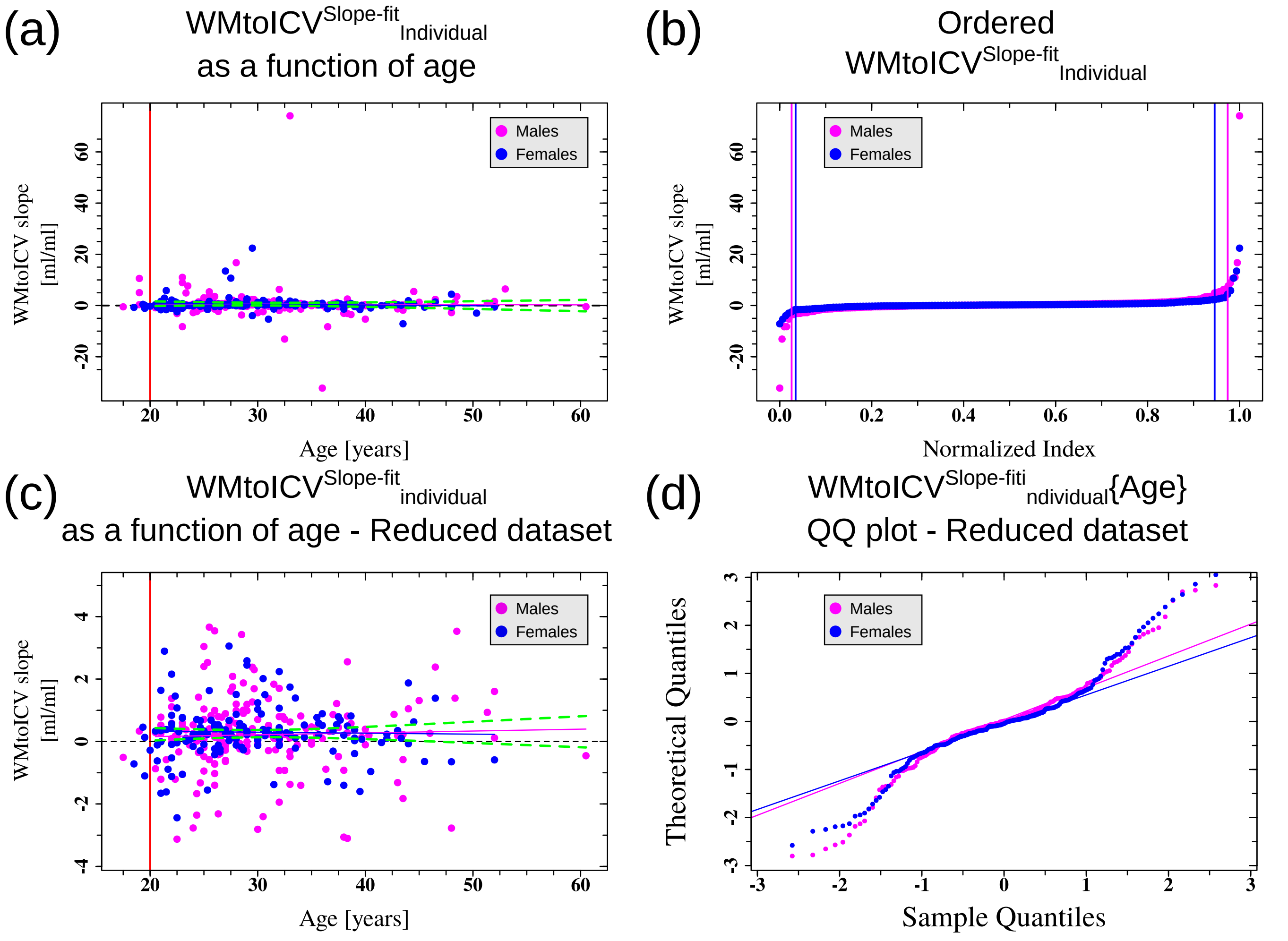


| | $WMtoICV^{Slope\text{-}fit}_{Individual}$ as a function of age | | | | $WMtoICV^{Slope\text{-}fit}_{Individual}$ as a function of age for the reduced dataset | | | |
|---|---|---|---|---|---|---|---|---|
| | Intercept | Slope [1/year] | $R^2$ | p | Intercept | Slope [1/year] | $R^2$ | p |
| Females | 1.5±0.9 | -0.03±0.03 | 0.0009 | 0.289 | 0.4±0.32 | -0.003±0.01 | -0.007 | 0.768 |
| Males | 0.9±1.9 | -0.01±0.06 | -0.005 | 0.865 | 0.04±0.38 | 0.006±0.01 | -0.004 | 0.621 |
| Both | 1.0±1.2 | -0.02±0.04 | -0.002 | 0.604 | 0.2±0.26 | 0.002±0.008 | -0.003 | 0.807 |

Figure S30: **Slope of the WM and ICV individual fits as a function of age.** Relationship between $\{WMtoICV^{Slope\text{-}fit}_{Individual\ i}\}$ and age. (a) Linear fit for the complete dataset. (b) Cumulative plot of $\{WMtoICV^{Slope\text{-}fit}_{Individual\ i}\}$ as a function of a normalized index between 0 and 1. Vertical lines represent the range of inclusion in the reduced dataset. (c) Linear fit for a reduced dataset that was constructed based on the criteria of panel (b). (d) Quantile-Quantile plot of the linear fit of the reduced dataset. Blue color is for females and pink for males. Dashed green lines are 95% confidence intervals of the fit for both females and males. The table below the graphs shows the fitting-values for the cases of females, males, and for the females and males together. For clarity, the fits for females and males together are not shown. $R^2$- goodness of fit. p-value - statistical significance of the fit. Error values - ±s.e.

### Females

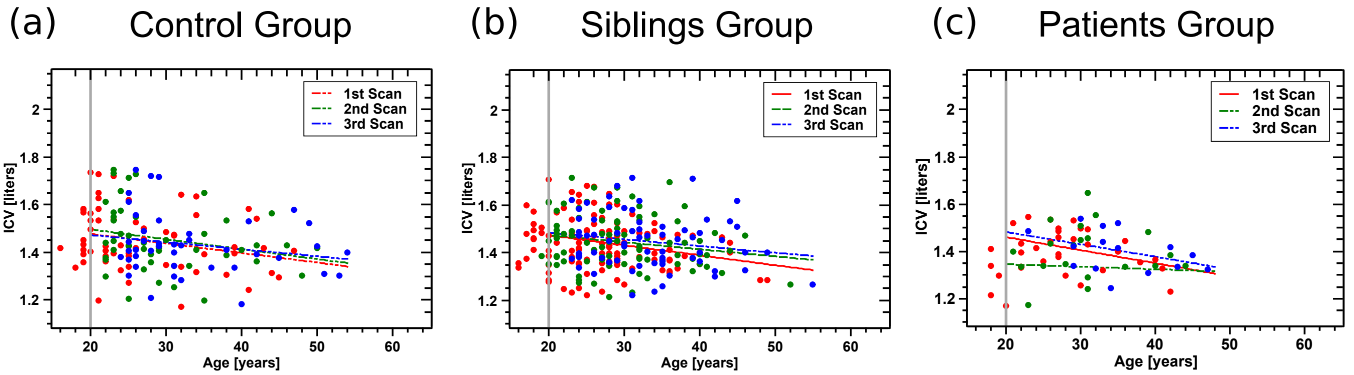


| | Control Group | | | | Siblings Group | | | | Patients Group | | | |
|---|---|---|---|---|---|---|---|---|---|---|---|---|
| | Intercept [liters] | Slope [liters/years] | $R^2$ | p (slope) | Intercept [liters] | Slope [liters/years] | $R^2$ | p (slope) | Intercept [liters] | Slope [liters/years] | $R^2$ | p (slope) |
| T1 | 1.56±0.07 | -0.004±0.002 | 0.035 | 0.103 | 1.56±0.05 | -0.004±0.001 | 0.049 | 0.019 | 1.57±0.07 | -0.006±0.002 | 0.120 | 0.037 |
| T2 | 1.58±0.08 | -0.004±0.003 | 0.037 | 0.106 | 1.53±0.05 | -0.003±0.002 | 0.022 | 0.092 | 1.37±0.13 | -0.001±0.004 | -0.051 | 0.793 |
| T3 | 1.53±0.08 | -0.003±0.002 | 0.019 | 0.189 | 1.54±0.06 | -0.003±0.002 | 0.017 | 0.138 | 1.59±0.11 | -0.005±0.003 | 0.102 | 0.122 |

### Males

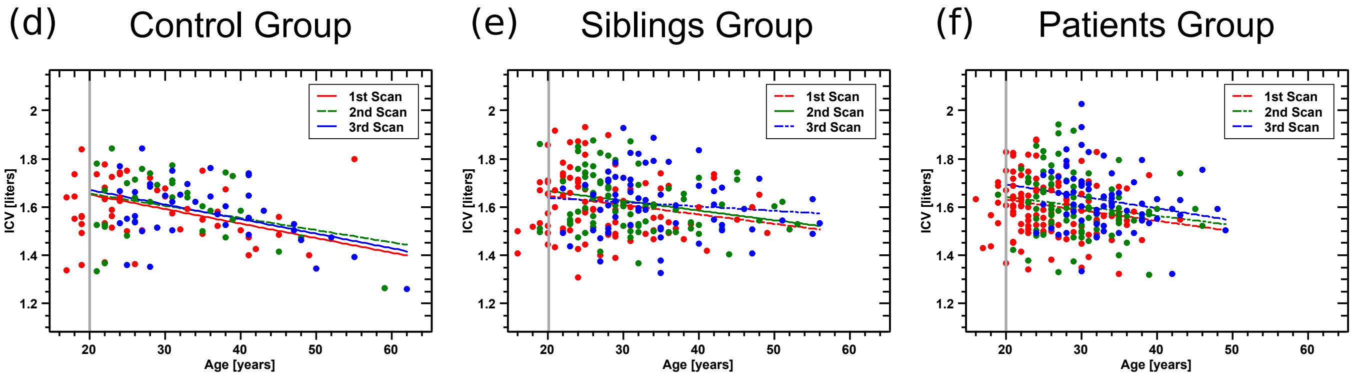


| | Control Group | | | | Siblings Group | | | | Patients Group | | | |
|---|---|---|---|---|---|---|---|---|---|---|---|---|
| | Intercept [liters] | Slope [liters/years] | $R^2$ | p (slope) | Intercept [liters] | Slope [liters/years] | $R^2$ | p (slope) | Intercept [liters] | Slope [liters/years] | $R^2$ | p (slope) |
| T1 | 1.77±0.07 | -0.006±0.002 | 0.149 | 0.008 | 1.72±0.06 | -0.003±0.002 | 0.029 | 0.065 | 1.72±0.06 | -0.004±0.002 | 0.027 | 0.052 |
| T2 | 1.76±0.08 | -0.005±0.003 | 0.076 | 0.060 | 1.75±0.06 | -0.004±0.002 | 0.060 | 0.019 | 1.72±0.07 | -0.004±0.002 | 0.017 | 0.116 |
| T3 | 1.79±0.07 | -0.006±0.002 | 0.180 | 0.004 | 1.67±0.07 | -0.002±0.002 | -0.003 | 0.374 | 1.79±0.09 | -0.005±0.003 | 0.037 | 0.062 |

Figure S31: **Cross-sectional results for the ICV behavior** (accompanying Fig. 5 of the main text). Cross-sectional ICV vs. age for the different groups in our study for (a-c) females and (d-e) males. Each dot represents the data for one individual. (a,d) control group; (b,e) group of siblings of subjects that were diagnosed with schizophrenia; (c,f) group of subjects that were diagnosed with schizophrenia. Red - first wave of the study, Green - second wave of the study, Blue - third wave of the study. In all panels, lines are fits of the different datasets to linear functions. Full lines - fits with a slope that is statistically significant at the $p < 0.05$ level. Striped lines - fits with a slope that is statistically significant at the $p < 0.1$ level. Striped-dotted lines - fits with a slope that is not statistically significant. Gray lines represent the minimum age for inclusion in the fits. The table below the graphs shows the fitting values for the cases of females, males. $R^2$- goodness of fit. p-value - statistical significance of the slope parameter of the fit. Error values - ±s.e.

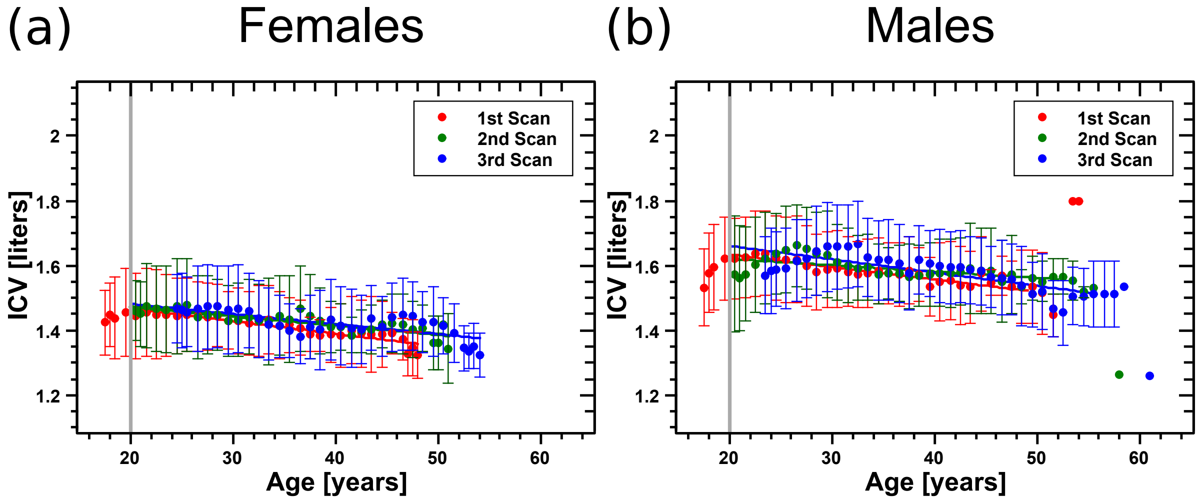


| | Females | | | | Males | | | |
|---|---|---|---|---|---|---|---|---|
| | Intercept [liters] | Slope [liters/years] | $R^2$ | p (slope) | Intercept [liters] | Slope [liters/years] | $R^2$ | p (slope) |
| T1 | 1.54±0.01 | -0.004±0.0003 | 0.85 | 3.1e-13 | 1.71±0.01 | -0.004±0.0004 | 0.79 | 6.6e-12 |
| T2 | 1.53±0.01 | -0.003±0.0003 | 0.69 | 6.8e-10 | 1.66±0.02 | -0.002±0.0004 | 0. 37 | 5.6e-05 |
| T3 | 1.54±0.02 | -0.003±0.0005 | 0.52 | 1.3e-06 | 1.74±0.02 | -0.004±0.0007 | 0.51 | 2.4e-06 |

Figure S32: **Running average analysis of the cross-sectional results** (accompanying Fig. 5 of the main text). Cross-sectional results for females (a) and males (b) separately were analyzed through a running average with a bin of 6 years. Dots and error bars - mean and s.d. of the running average analysis at the first (red), second (green) and third (blue) waves of the study. Full lines are fits of the running averages to the corresponding linear functions. Gray vertical lines - start age for the fits. End of lines - end age for the fits. The table below the graphs shows the fitting values for the cases of females or males. $R^2$- goodness of fit. p-value - statistical significance of the slope parameter of the fit. Error values - ±s.e.

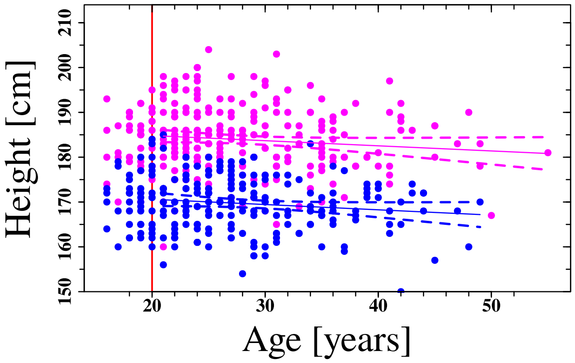


| | Intercept [cm] | Slope [cm/year] | $R^2$ | p |
|---|---|---|---|---|
| Females | 173±2.0 | -0.1±0.07 | 0.01 | 0.084 |
| Males | 187±2.0 | -0.1±0.07 | 0.007 | 0.094 |
| Both (mean adjusted) | 4±1.5 | -0.1±0.05 | 0.001 | 0.019 |

Figure S33: **Height as a function of age**. Height as a function of age for females (blue) and males (pink). Each dot represents the data for one individual. Straight lines are fits of the data to liner functions for males (pink) and females (blues). The table below the graphs shows the fitting values for the cases of females, males, and for the females and males together. Dashed lines are 95% confidence intervals of the fits. Red line represents the minimum age for inclusion in the fits. $R^2$- goodness of fit. p-value - statistical significance of the slope parameter of the fit. Error values - ±s.e.
